# The biophysical interaction network that couples zona occludens-1 to membrane proteins at tight junctions

**DOI:** 10.64898/2026.10.06.757067

**Authors:** Srajan Kapoor, Jagrity Choudhury, Alex J. Vecchio

**Affiliations:** Department of Structural Biology, Jacobs School of Medicine and Biomedical Sciences, University at Buffalo, Buffalo, NY 14203 USA

## Abstract

Tight junctions are dynamic membrane-bound protein assemblies that maintain barrier function for tissues as cells divide, grow, and move through development and throughout life. Maintenance of tissue barriers requires spatially separated intracellular and extracellular signaling pathways to be linked through novel and dynamic protein interaction networks. Tight junction intracellular signaling is governed by zona occludens-1 (ZO-1), a Membrane-Associated GUanylate Kinase (MAGUK) family protein that recruits and assembles transmembrane proteins and scaffolds them to the actin cytoskeleton through its own oligomerization, localization to membranes, and phase separation. Here we report comprehensive quantitative studies that nearly completely maps the intracellular interaction network between ZO-1 and tight junction transmembrane proteins that elucidates the biophysical principles governing their binding and a mechanism for how stimuli transmit through membranes. We establish where, how, and with whom each protein interacts, and identify how membrane proteins enable ZO-1 and greater tight junction function through specific interactions or by phase separation. Our findings reveal the intracellular interactome that triggers and regulates tight junction assembly, the active and direct roles each membrane protein plays in these processes, and provide a mechanism for how intra/extracellular signaling is coupled to transmit mechanochemical signals between cells via tight junctions. These insights provide a prospective blueprint for how other MAGUK/membrane protein interaction networks form and function across biology.

## Introduction

Membrane-Associated GUanylate Kinase (MAGUK) family proteins regulate numerous and critical biological processes that include neurotransmission, paracellular transport, and other cell signaling events where molecular communication between two disparate sites is required to occur.^<u>1-9</u>^ At unique and specific regions beneath the plasma membrane, MAGUK proteins act as protein-protein interaction hubs through their multi-modular organization that contains several folded structural domains linked together by intrinsically disordered regions (IDRs) with highly variable but low-complexity sequences. PDZ proteins constitute the structural domains of MAGUKs, which themselves are a ubiquitous protein subfamily that specializes in binding complementary peptide motifs on the intracellular termini of unrelated proteins.^<u>10</u>^ These PDZ domain/PDZ motif interactions drive assembly of large heterogeneous membrane-associated protein complexes that transmit their intended signals through spaces or junctions to adjacent complexes. The plethora of MAGUK and PDZ domain proteins, coupled with their at times promiscuous, or at others, selective binding to PDZ motifs of target and off-target proteins, has confounded understanding of individual systems and masked the general principles that guide their association.^<u>11-15</u>^

Tight junctions are a system that employs a MAGUK PDZ domain-containing protein to recruit and assemble integral plasma membrane proteins and scaffold these complexes to the actin cytoskeleton.^<u>16</u>^ This protein, zona occludens-1 (ZO-1), is one of the first identified MAGUKs, ∼200 kDa, and contains six distinct protein binding modules, three of which are PDZs.^<u>9</u>^ From N- to C-terminus ZO-1 contains the structured domains PDZ1, PDZ2, PDZ3, SH3 and GuK, which are linked together by IDRs called unique (U) motifs, and ultimately terminates with an actin binding domain (**Fig. 1a**). Using the first five domains, ZO-1 binds the intracellular C-termini of three distinct families of tight junction membrane proteins (TJMPs) called claudins, junctional adhesion molecules (JAMs), and the tight junction-associated Marvel proteins (TAMPs) occludin and tricellulin, amongst others.^<u>17-23</u>^ In a carefully orchestrated but unresolved network of interactions, these proteins facilitate the formation and function of tight junctions, which act as the primary barriers to paracellular molecular transport between individual cells in epithelial and endothelial tissues. The critical barrier function that tight junctions impart to tissues makes resolving this interaction network key to understanding tight junction assembly, disassembly, and regulation— and how these processes inform signal transduction through membranes and between cells.

**Fig. 1.**
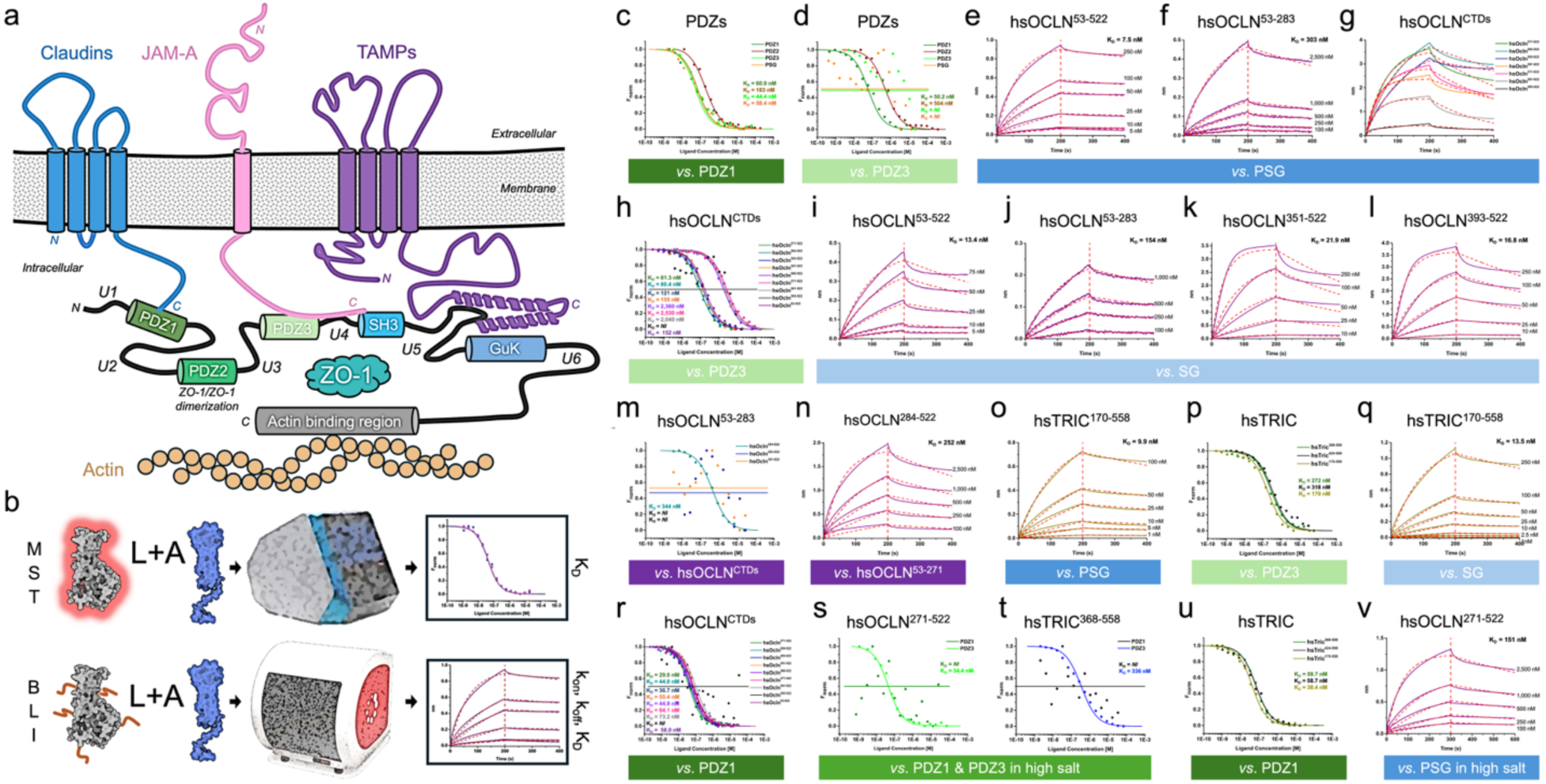
Existing model of ZO-1/TJMP assembly and quantification of biophysical interactions between ZO-1 subdomains and ZO-1 and TAMPs. (a) Schematic representation of previously determined interactions between ZO-1 and TJMPs. The three major TJMP families include claudins (blue), JAMs (pink), and TAMPs (purple). ZO-1s is organized from N- to C-term by PDZs −1 (dk green), −2 (green), −3 (lt. green), SH3 (blue), GuK (lt. blue), and actin binding region (grey) that interacts with actin (brown). Unstructured unique (U) motifs link ZO-1 structured domains. The C-terminal domains of TJMPs interact with their complementary ZO-1 domains. Note that PDZ2 functions as a dimerization domain for ZO-1 oligomerization. (b) Biophysical technique workflow. Chemically modified ligands (L) are incubated with unmodified analytes (A) at various concentrations in solution. Binding is detected by instruments, resulting in binding curves that yield thermodynamic affinities for both and kinetics for BLI. (c-v) Results from our biophysical interactions between ZO-1 subdomains and ZO-1 and TAMPs. Unmodified analyte is shown in top description and modified ligand in bottom description (colored boxes), with binding curves plotted between. See **Tables 1-2** for full results.

**Table 1.** PDZ Binding Studies.

| Ligand | Analyte | Fit | $K_D$ (nM) | Fig. |
| --- | --- | --- | --- | --- |
| PDZ1 | PDZ1 | 1:1 | 60.9 ± 31.3 | **1c** |
| PDZ1 | PDZ2 | 1:1 | 183.0 ± 51.0 | **1c** |
| PDZ1 | PDZ3 | 1:1 | 44.4 ± 13.0 | **1c** |
| PDZ1 | PSG | 1:1 | 58.4 ± 21.2 | **1c** |
| PDZ3 | PDZ1 | 1:1 | 50.2 ± 13.9 | **1d** |
| PDZ3 | PDZ2 | 1:1 | 504.0 ± 119.3 | **1d** |
| PDZ3 | PDZ3 | 1:1 | *NI* | **1d** |
| PDZ3 | PSG | 1:1 | *NI* | **1d** |
| PSG | PSG | 1:1 | *NI* | **N/A** |

*N/A* = Not Applicable
*NI* = No Interaction detected

**Table 2.**
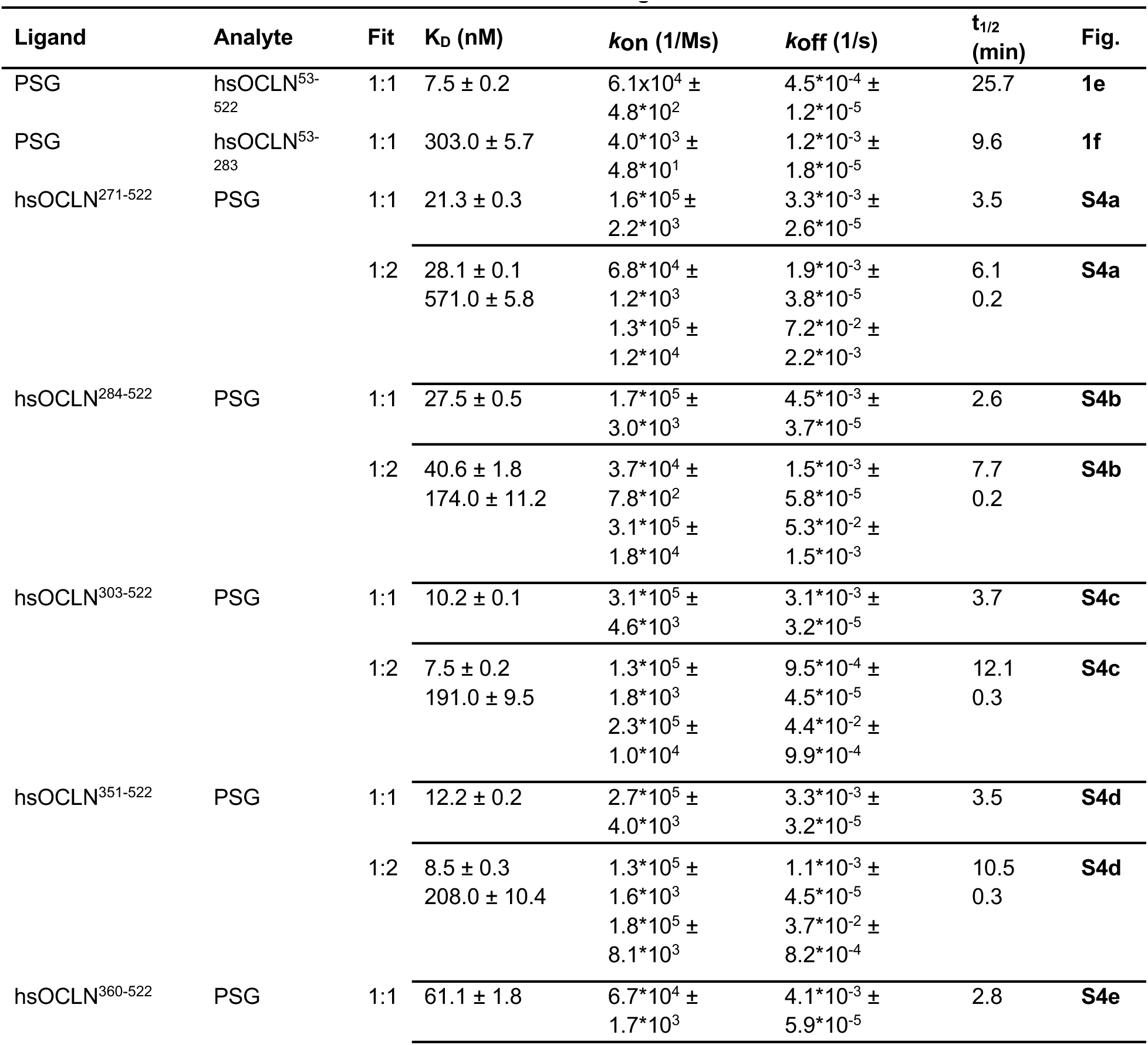
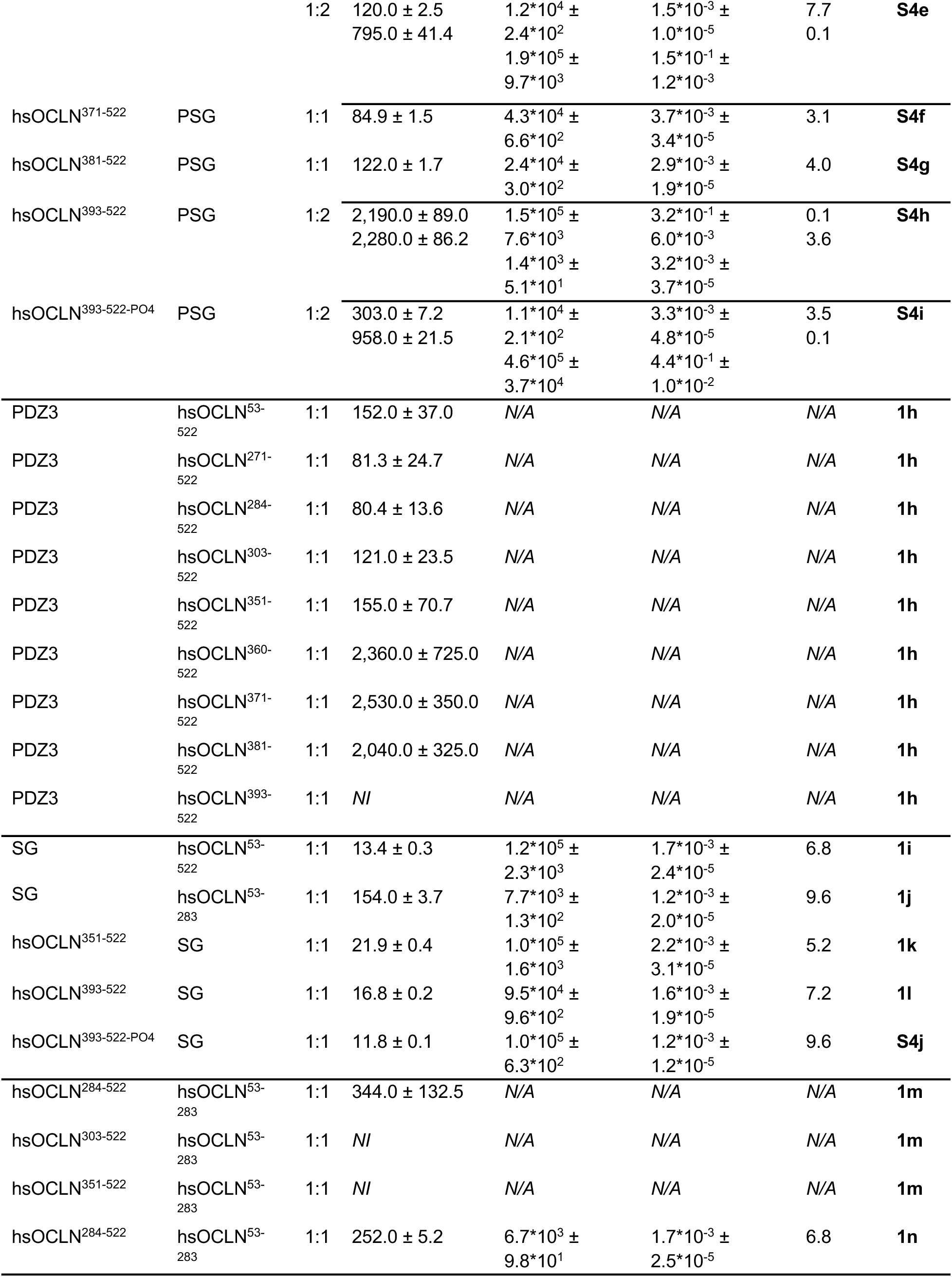
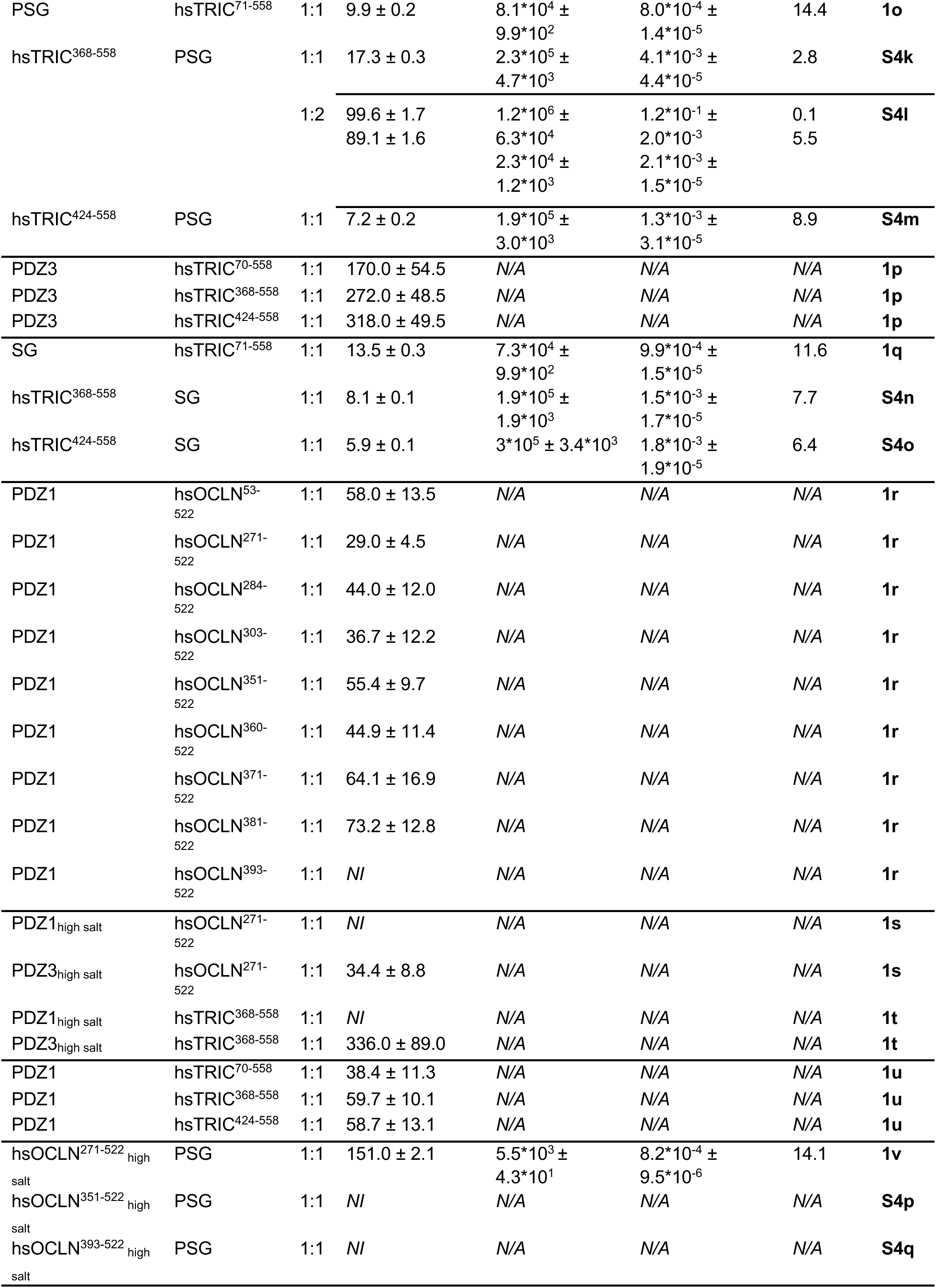
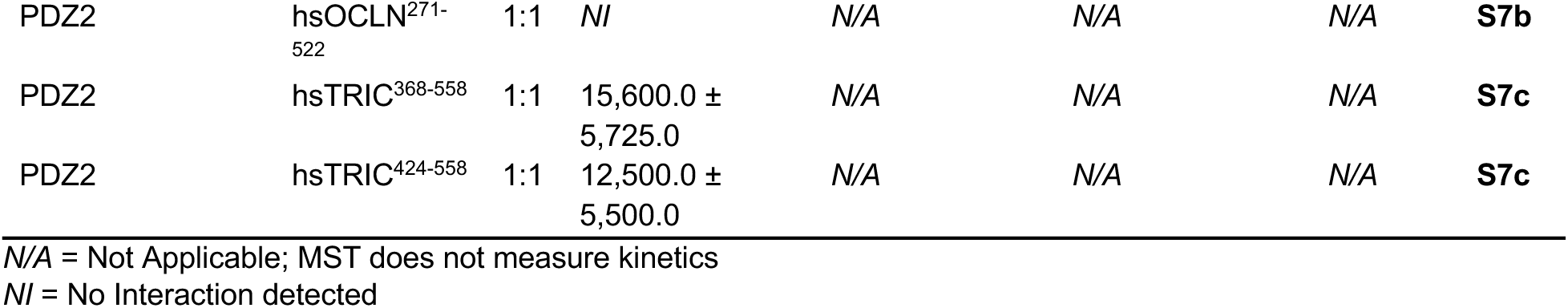
TAMP Binding Studies.

From outside the cell in, TJMPs possess overlapping and distinct topological and structural features that direct their specific functions at tight junctions (**Fig. 1a**). Claudins, JAMs, and TAMPs have extracellular segments (ECS) of varying structure and size that enable their lateral (cis) interactions within the same membrane and subsequent longitudinal (trans) interactions with their equivalents on neighboring cells across paracellular space.^<u>24-27</u>^ The cis interactions between TJMPs form tight junction strands that connect at branching points on the surfaces of cells, while the trans interactions adhere neighboring cells and regulate paracellular molecular transport between cells.^<u>27-30</u>^ The transmembrane (TM) domains of claudins and TAMPs contain four helices while JAMs traverse the membrane with one TM. The N- and C-termini of claudins and TAMPs are intracellular while for JAMs only the C-terminus resides inside the cell. The sequences, lengths, and structures of TJMP intracellular segments are extremely variable apart from claudins and JAMs possessing PDZ binding motifs at their extreme C-termini.^<u>19-21</u>^ TAMPs have no PDZ motif but instead have a conserved coiled-coil domain (CCD) at their C-terminus.^<u>31-33</u>^ Despite the diversity of TJMP intracellular segments, all are known to use it to bind specific PDZs of ZO-1. ^<u>20,21,23</u>^ Intracellular links to ZO-1 recruit TJMPs to tight junctions, stabilize greater assemblies through cytoskeletal scaffolding, and affect strand morphology and paracellular barrier function of tight junctions.^<u>16,34,35</u>^ For this to be true, the intracellular binding networks between ZO-1 and TJMPs on one cell must communicate with neighboring complexes on another through paracellular space. Elucidating the intracellular binding network can resolve the basis for how communication is initiated.

ZO-1’s PDZ domains are thought to have specific TJMP binding partners. The model in **Fig. 1a** summarizes our current understanding. Here, the PDZ motif of claudins interacts with PDZ1; the PDZ2 domain drives ZO-1 dimerization; JAM-A’s PDZ motif interacts with PDZ3 and requires the SH3 domain; and TAMPs use their CCD to bind the SH3/GuK domain.^<u>20,21,23</u>^ In this model, each TJMP interacts with a specific PDZ or module of ZO-1. However, PDZs are known to be promiscuous or selective depending on the physicochemical properties of their receptors.^<u>11-15</u>^ The current model of ZO-1/TJMP interactions thus defies typical PDZ complexity and diverges from how better characterized MAGUK proteins function. Because tight junctions are not static, other biochemical modifications or biophysical phenomena affect this system and the timescales of its association. Specifically, phosphorylation of residues on ZO-1 or TJMP intracellular segments modulate ZO-1 binding, which in turn alters tight junctions and barrier function.^<u>36,37</u>^ Moreover, the localization and lifespans of TJMPs at tight junctions are tightly controlled, where occludin is constitutively associated but dynamic within junctions while claudins are stable in junctions but continuously integrated at rates that differ by subtype.^<u>38,39</u>^ We sought to close the knowledge gaps that existed to identify the hidden biophysical characteristics and principles that govern TJMP association with ZO-1 through comprehensive quantitative studies.

Here, we elucidate this interaction network by validating established interactions and defining interactions that extend across the broader system. This was accomplished by developing biochemical methods to produce and isolate full-length, mutant, and truncated TJMPs, single PDZs, and multi-modular sub-domains of ZO-1; and the application of state-of-the-art biophysical techniques. Analyzing these 43 proteins—21 of which are membrane proteins and 13 full-length receptors—with this approach enabled robust, reproducible, and precise experimental determination of the thermodynamic binding affinities, and in some cases, the kinetics, of ZO-1/TJMP interactions. In full, we elucidate the intra- and inter-molecular ZO-1/TJMP interactome and the biophysical principles guiding their assemblies that in turn direct form and function of tight junctions. The work provides new paradigms to understand tight junction assembly and signaling that may apply generally to other MAGUK, PDZ, and membrane protein systems that perform diverse functions that span biology.

## Results

### Biochemical preparation of ZO-1 domains and TJMPs

We designed constructs of ZO-1 and TJMPs for expression and purification to assess their interactions that included full-length forms, isolated domains, truncations, or mutants (**Fig. S1**). These proteins were expressed in bacterial or insect cells depending on the complexity of their folding, then purified using affinity and size-exclusion chromatography (SEC) to isolate monodisperse samples using new or established methods (see **Methods**).^<u>40-44</u>^ We found that TJMPs with TM domains were not produced or folded using bacterial systems while their C-terminal domains were, as assessed by biochemical analyses, crystal structures, and validated previously by other labs (**Fig. S2**).^<u>20,21,31,32</u>^ We previously showed that insect cells produce folded and functional TJMPs.^<u>41</u>^ These folded, stable, and purified proteins were then used in biophysical studies to determine their interaction networks.

### Overview of biophysical interaction methods

Once proteins were produced and isolated, the post-affinity pure constructs were chemically modified for two distinct biophysical workflows (**Fig. 1b**). For microscale thermophoresis (MST), which requires a ligand to be fluorescently labeled, we did so using NHS-Cy5 dye at 2-fold molar excess of ligand. Free dye was removed via SEC then labeled ligand was used at a fixed concentration in MST experiments against a concentration range of unlabeled protein analytes. Preference for labeled ligand was given to soluble proteins over membrane proteins, whose stability and sizes were more varied. For bio-layer interferometry (BLI), which requires a ligand to be immobilized, we biotinylated the ligand using NHS-PEG4-biotin at 2-fold molar excess, removed free biotin using SEC, then immobilized the ligand at a fixed concentration on streptavidin biosensors and tested binding against a concentration range of protein analytes. In most cases, the smaller molecular weight soluble protein was immobilized. All subsequent BLI or MST binding analyses were performed in an identical buffer at physiological pH and room temperature.

### Structural homology of PDZ domains suggest ligand binding mode overlap

We first analyzed the structures of ZO-1’s three PDZ domains. We found that PDZ1, PDZ2, and PDZ3 were highly similar, with root mean square deviations of <1.0 Å, and had conserved PDZ motif binding grooves on their surfaces but varied outside of this core (**Fig. S3a**). Surface representation showed that the surface topology of PDZs were unique, but also that the PDZ motif of a TJMP peptide fit well within the ligand binding grooves of all three PDZs, indicating structural conservation (**Fig. S3b**).^<u>45-47</u>^ Despite a similar shaped ligand binding groove, the grooves were electrostatically charged in different ways. For PDZ1 and PDZ3, the proposed PDZ motif binding grooves were largely non-polar but are flanked by charged surfaces outside (**Fig. S3c**). Modeling the PDZ motif of a TJMP peptide into the PDZ binding grooves of PDZ1-3 showed that the peptide conforms to the surface groove, that side chains occupy small pockets within the groove, but also that the groove is continuous on both ends that the peptides bind. This indicated that a peptide could bind within these grooves at multiple points, as there was no clear beginning or end where only a C-terminal PDZ motif would fit. A highly basic pocket would be expected to bind the carboxyl termini of a TJMP, which was also not present. Based on this analysis, we speculated that there were numerous ways in which a TJMP terminal peptide could bind ZO-1 PDZs.

### PDZ domains within ZO-1 exhibit selective intra-molecular interplay

We next characterized intra-molecular interactions between ZO-1 domains to establish a baseline for our biophysical studies. Specifically, we tested PDZ1 and PDZ3 binding to PDZ1, PDZ2, PDZ3, and the tripartite PDZ3/SH3/GuK (PSG) module using MST. We found that PDZ1 bound itself with an equilibrium dissociation (K_D_) of 60.9 nM, while it bound PDZ2, PDZ3, and PSG with K_D’s_ of 183.0, 44.4 and 58.4 nM, respectively (**Fig. 1c** and **Table 1**). For PDZ3, we found it did not self-associate or bind PSG but did bind PDZ1 and PDZ2 with 50.2 and 504.0 nM affinities, respectively (**Fig. 1d** and **Table 1**). Consistent with this result, we found that PSG did not self-associate (**Table 1**). These results demonstrated that PDZ/PDZ intra-molecular interactions exist within ZO-1 and that their interplay is selective.

### Occludin interacts with high-affinity to the PSG module of ZO-1

Because interactions between occludin containing its TM domain and ZO-1 had not been tested *in vitro* and previous studies showed that occludin’s CCD interacts with ZO-1’s GuK domain and U5 motif with micromolar affinity, we sought to define and quantify the affinities of all interactions between these proteins using our cache of constructs.^<u>16,23</u>^ We first tested the binding between N-terminally truncated but otherwise full-length occludin (hsOCLN^53-522^) to PSG using BLI to establish their K_D_ and association and dissociation rate constants (k_on_ and k_off_). We found that hsOCLN^53-522^ bound to immobilized PSG with a K_D_ of 7.5 nM and used the k_off_ rate to measure that the half-life (t_1/2_) of the complex was 25.7 minutes (**Figs. 1e** and **Table 2**). These results established occludin binding to PSG and the high affinity of this interaction compared to that of CCD/GuK suggested that regions outside of the occludin’s CCD contribute to occludin/ZO-1 complex formation.

We next aimed to identify these regions by using occludin constructs that lacked a C-terminal domain or TM-embedded Marvel domain to test binding against PSG (**Fig. S1**). We isolated the Marvel domain of occludin (hsOCLN^53-283^) and found that it bound immobilized PSG with a K_D_ of 303.0 nM and t_1/2_ of 9.6 minutes (**Fig. 1f** and **Table 2**). We next used occludin constructs that lacked Marvel domains and only contained intracellular C-termini of variable lengths and found in single-point BLI experiments that all immobilized C-terminal domains of occludin bound PSG, albeit with different magnitudes (**Fig. 1g**). Specifically, we found that shortening occludin’s C-terminal domain from its N-terminus caused a progressive loss of PSG binding where the two shortest constructs, hsOCLN^381-522^ and hsOCLN^393-522^, resulted in significantly decreased binding. As single-point BLI analyses were qualitative, we performed quantitative multi-point analyses of immobilized occludin C-terminal domains to test binding against multiple concentrations of PSG (**Figs. S4a-h**). These results showed the same trends, that occludin with longer C-terminal domains bound PSG better than shorter constructs. We determined binding affinities of 21.3, 27.5, 10.2, 12.2, 61.1, 84.9, 122.0, and >2,000 nM for the hsOCLN^271-522^ through hsOCLN^393-522^ constructs, respectively (**Table 2**). Noticeably, occludin with longer C-terminal domains fit a 1:1 binding model at low concentrations of PSG (<100 nM) but a 2:1 binding model at concentrations >100 nM, indicating multiple interactions with PSG were occurring. Finally, because the region upstream of occludin’s CCD contains phosphorylation sites that attenuate its binding to ZO-1, we made a phosphorylation mimetic of the hsOCLN^393-522^ construct (hsOCLN^393-522-PO4^) by mutating in six glutamates.^<u>48-54</u>^ We found that hsOCLN^393-522-PO4^ bound PSG with higher affinity compared to native hsOCLN^393-522^ but had a nearly identical complex half-life (**Fig. S4i** and **Table 2**). This showed that phosphorylation affects association rates. These findings ascribed quantitative binding contributions of each major domain of occludin to ZO-1’s PSG module, which regions drive binding, and how phosphorylation modulates occludin interactions with ZO-1 to alter tight junction function.

### Occludin interacts with variable affinities to sub-domains of the PSG module

As our results suggested that regions outside of occludin’s CCD and PSG’s GuK domains play roles in complex formation, we sought to determine the roles of PSG’s PDZ3 and SH3 domains in binding to occludin. First, we used labeled PDZ3 and tested binding against occludin C-terminal constructs using MST (**Fig. 1h** and **Table 2**). We found that the longest C-terminal constructs of occludin, hsOCLN^271-522^ to hsOCLN^351-522^, bound PDZ3 with <155.0 nM affinity, while shorter constructs hsOCLN^360-522^ to hsOCLN^381-522^ bound with >2,000 nM affinity—the shortest construct, hsOCLN^393-522^ did not bind PDZ3. This experiment defined the region of occludin’s C-terminus that binds PDZ3.

We next determined occludin binding against the SH3/GuK (SG) domain using BLI. We first tested SG binding to occludin and to occludin’s Marvel domain and found the former bound with 13.4 nM and the latter with 154.0 nM, respectively (**Figs. 1i-j** and **Table 2**). These affinities appeared similar to our results with PSG, but kinetic analysis showed that the occludin/PSG complex t_1/2_ was 3.8-fold longer-lived than the occludin/SG complex at 25.7 vs. 6.8 minutes (**Table 2**). We then used C-terminal constructs hsOCLN^351-522^ and hsOCLN^393-522^, which showed high and no binding to PDZ3 using MST, and tested their interactions with SG (**Figs. 1k-l** and **Table 2**). We found that hsOCLN^351-522^ bound SG with 21.9 nM affinity while hsOCLN^393-522^ bound SG with 16.8 nM. A high affinity interaction between SG and hsOCLN^393-522-PO4^ was measured to be 11.8 nM (**Fig. S4j** and **Table 2**). These results differed from PDZ3 and PSG binding data. Specifically, the observation that hsOCLN^393-522^ bound SG with 129-fold higher affinity than PSG. These experiments provided a basis for occludin binding to specific sub-domains of the PSG module and isolated the regions driving occludin/PSG assembly.

### Occludin sub-domains interact, forming a multi-valent module

Because our finding that occludin bound PSG and SG better than the Marvel domain alone suggested that the membrane-embedded and C-terminal domains of occludin were in close proximity to each other, we tested this idea experimentally. Using the labeled Marvel domain of occludin and MST, we tested binding against progressively shorter C-terminal variants and found that only the longest variant hsOCLN^284-522^ bound Marvel (**Fig. 1m** and **Table 2**). We quantified this interaction using BLI with an immobilized C-terminal domain and showed that this complex interacted with a K_D_ of 252.0 nM and had a half-life of 6.8 minutes (**Fig. 1n** and **Table 2**). This result established that the membranous and intracellular domains of occludin are more than linked together through an unstructured peptide but are complexed to form a multivalent module to bind ZO-1 and also defined which region of occludin’s C-terminal domain is required to form this complex.

### Tricellulin also interacts with high affinity to PSG

As the two homologous TAMPs occludin and tricellulin integrate at bi-cellular or tri-cellular tight junctions, respectively, to perform overlapping yet distinct functions, we tested whether our findings for occludin were consistent in tricellulin.^<u>28,55</u>^ First, we performed sequence alignments of human occludin and tricellulin (**Fig. S5**). We found that the Marvel domains and CCD of TAMPs aligned well, which agreed with previous structural results.^<u>32,33</u>^ However, this analysis also showed that both TAMPs have highly varied N- and C-terminal domain sequence and length and specifically that tricellulin lacked 48 amino acids that were present in occludin between the Marvel and CCDs in an IDR. We tested whether PSG bound tricellulin differently using BLI and found, like occludin, that PSG bound tricellulin with high affinity using a K_D_ of 9.9 nM and a t_1/2_ of 14.4 minutes (**Fig. 1o** and **Table 2**). We next tested tricellulin C-terminal domain constructs that lacked a Marvel domain and were progressively shorter. We found hsTRIC^368-558^ and hsTRIC^424-558^ bound PSG with affinities of 17.3 nM and 7.2 nM, respectively (**Fig. S4k-m** and **Table 2**). Like with occludin, we then tested tricellulin interactions to PDZ3 and the SG domain alone. We found that tricellulin (hsTRIC^70-558^) and the two C-terminal domain constructs hsTRIC^368-558^ and hsTRIC^424-558^ bound PDZ3 with affinities of 170.0, 272.0 and 318.0 nM, respectively (**Fig. 1p** and **Table 2**). For tricellulin and its C-terminal domain against SG we found that tricellulin bound SG with a K_D_ of 13.5 nM and had a t_1/2_ of 11.6 minutes, while the two variants that lacked a Marvel domain bound with 5.9-8.1 nM to SG (**Figs. 1q** and **S4n-o** and **Table 2**). As with occludin, these affinities were similar to PSG. But unlike occludin, kinetic data showed that the tricellulin/PSG complex t_1/2_ was only 1.2-fold longer-lived than the tricellulin/SG complex at 14.4 vs. 11.6 minutes (**Table 2**). In full, these results showed that tricellulin’s C-terminal domain bound PDZ3 with 3.3-fold lower affinity than occludin’s equivalent domain, and that tricellulin’s affinity to PSG or SG was not significantly different whereas occludins was. These results explain the overlapping functions of TAMPs and how distinct TAMP functions may be attributed to alternate binding modes to ZO-1.

### PDZ1 and PDZ3 bind TAMPs differently and with distinct mechanisms

Although our binding data to PDZ3 was new, it was somewhat expected that TAMP C-terminal domains bind PSG and SG based on previously reported interactions with U5 and GuK.^<u>16,23,56-59</u>^ However, analysis of PDZ structures showed us that PDZ1, PDZ2, and PDZ3 of ZO-1 were highly similar and had conserved PDZ motif binding grooves (**Fig. S3**). This led us to wonder if ZO-1’s PDZs have overlapping inter-molecular interactions. We labeled PDZ1 and tested binding against occludin C-terminal constructs using MST to see. Unlike what was observed for PDZ3 (**Fig. 1h**), here we found that all but the shortest C-terminal construct of occludin, hsOCLN^393-522^, bound well to PDZ1 (**Fig. 1r**). Measurements showed that longer constructs bound PDZ1 with K_D’s_ <73.2 nM (**Table 2**). These results revealed differences in occludin binding to PDZ1 and PDZ3 that we analyzed in greater detail.

As PDZ3 binding to occludin C-terminal constructs showed an abrupt change in affinity between hsOCLN^351-522^ and hsOCLN^360-522^, and this region was not present in tricellulin, we surmised that the chemistry or length of TAMP C-terminal residues influence PDZ binding (**Figs. 1r** and **S5**). To test this, we performed binding studies in the presence of 500 mM salt, as structural analysis showed PDZs have variable charged surfaces (**Fig. S3c**). We found for both hsOCLN^271-522^ and hsTRIC^368-558^, which are the longest construct of each TAMP, that high salt abrogated interactions with PDZ1 but had no effect on PDZ3 binding (**Figs. 1s-t** and **Table 2**). Specifically, for hsOCLN^271-522^ and hsTRIC^368-558^, high salt decreased their affinities to PDZ1 from 29.0 nM and 59.7 nM to undetectable, respectively. While high salt altered their affinities to PDZ3 from 81.3 nM and 272.0 nM to 34.4 nM and 336.0 nM, respectively. To obtain this comparison, we had to perform binding studies of our three tricellulin constructs to PDZ1, which, like PDZ3 (**Fig. 1p**) showed that all three bound well with affinities between 38.4-59.7 nM (**Fig. 1u** and **Table 2**). We also tested occludin C-terminal domain interactions to PSG in high salt and found that hsOCLN^271-522^ decreased affinity from 21.3 nM to 151 nM but the complex half-life was extended 4-fold (**Fig. 1v** and **Table 2**). This showed that individual domains of PSG are driven by distinct chemical forces. For shorter constructs hsOCLN^351-522^ and hsOCLN^393-522^, no interactions with PSG were detected (**Figs. S4p-q** and **Table 2**). In full, these experiments showed that all tricellulin constructs bound both PDZ1 and PDZ3, all but occludin’s CCD bound PDZ1 and PDZ3, and that electrostatics govern interactions to PDZ1 and SG but not PDZ3. Comprehensively, they highlighted that TAMPs use overlapping yet distinct mechanisms to bind ZO-1 and that individual PDZ domains discern their binding sites through chemical properties inherent to their ligands.

### Claudins interact with PDZ1 and PDZ3 but do not use their PDZ motifs to do so

Previous studies showed that claudins bind PDZ1 of ZO-1 and likely do so using the PDZ motifs at their extreme C-termini.^<u>19,20,60</u>^ However, it has also been shown that deletion of a claudin’s PDZ motif had no effect on localization or assembly of claudins at tight junctions.^<u>61,62</u>^ It was thus largely assumed that loss of the PDZ motif caused loss of ZO-1 binding and that this loss was not detrimental to tight junction formation. Based on the continuous PDZ motif binding groove conserved in PDZs, coupled with no definitive basic pocket to bind the last carboxyl residue, we speculated that PDZs did not bind the C-terminal ends of claudins. To test this idea, we designed MST experiments to quantify interactions between 10 claudin subtypes against the labeled PDZ1 and PDZ3 domains of ZO-1. Claudins having high variability in sequence and length of their C-termini but also a conserved Tyr-Val terminus were chosen (**Fig. 2a**). We found that all claudins bound PDZ1 well, with affinities that ranged from 52.9 to 243.0 nM (**Fig. 2b** and **Table 3**). We next assayed PDZ1 binding to claudin-4 where 12 (hsCLDN-4^Δ12^) or 24 (hsCLDN-4^Δ24^) residues were removed from its C-terminus (**Fig. S1**). We found that full-length and hsCLDN-4^Δ12^ bound PDZ1 with ∼130.0 nM affinity but that hsCLDN-4^Δ24^ bound PDZ1 with >6,000 nM affinity—a 45-fold decrease (**Fig. 2c** and **Table 3**). These findings were consistent with the observation that claudins bind PDZ1 but also showed that a PDZ motif is not required for this interaction.

**Fig. 2.**
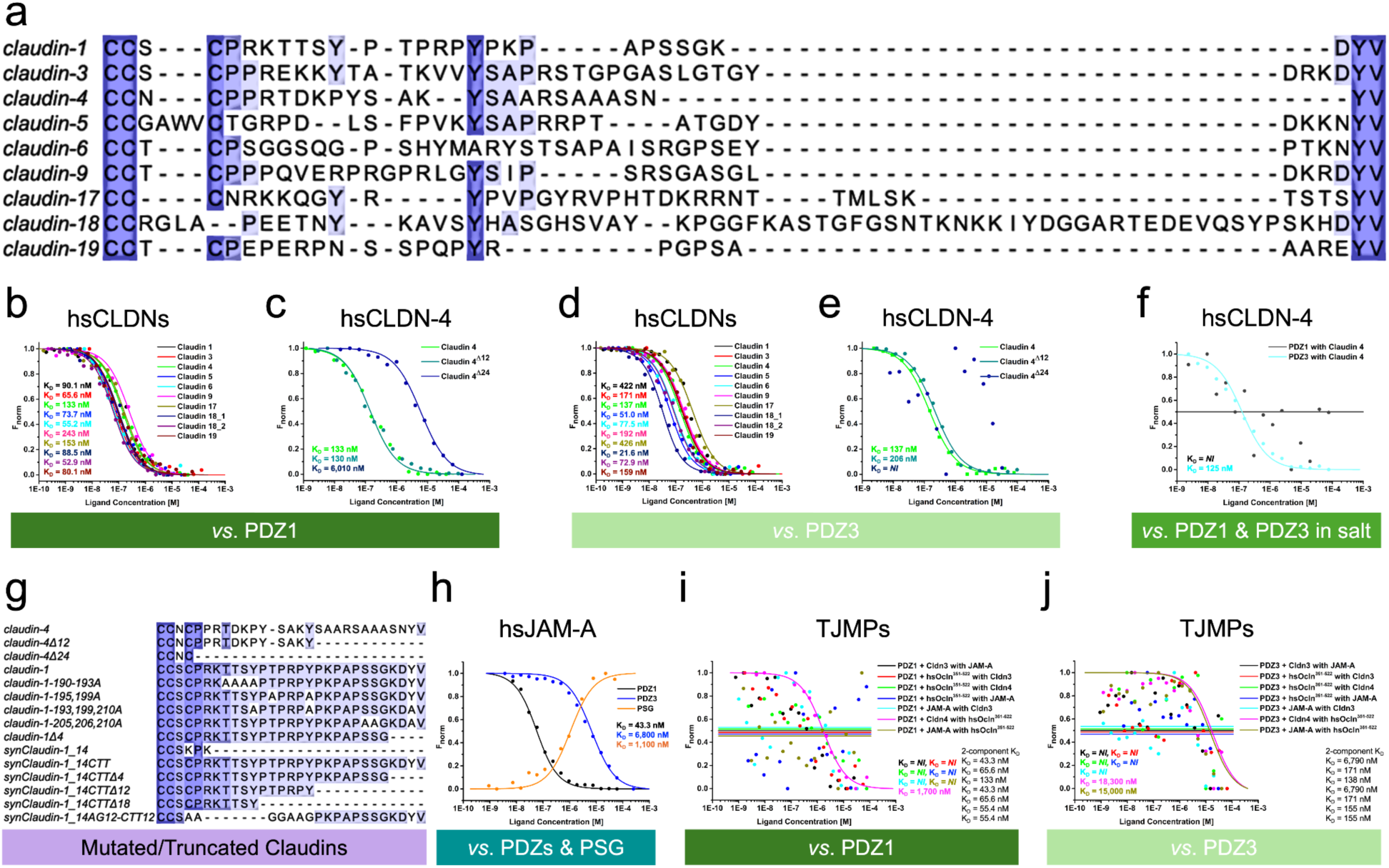
Quantification of biophysical interactions between ZO-1 and claudins, JAM-A, and competitive binding between TJMPs for PDZs. (a) Sequence alignment of the C-terminal domains of assayed human claudins with an added initial Cys to improve alignment. Intensity of shading (purple) indicated increased identity. (b-f) Results from our biophysical interactions between ZO-1 and claudins. See **Table 3** for full results. (g) Sequence alignments of the C-terminal domains of claudins −1, −4, and synClaudin-1 showing mutations and truncations to this region. Binding data for these analyses appear in **Figs. S6a-e**. (h) Biophysical interactions between ZO-1 and JAM-A. See **Table 3** for full results. (i-j) Three-component competition assays testing the ability of PDZ1 or PDZ3 mixed with a single TJMP to bind another TJMP. See **Table 4** for full results.

**Table 3.**
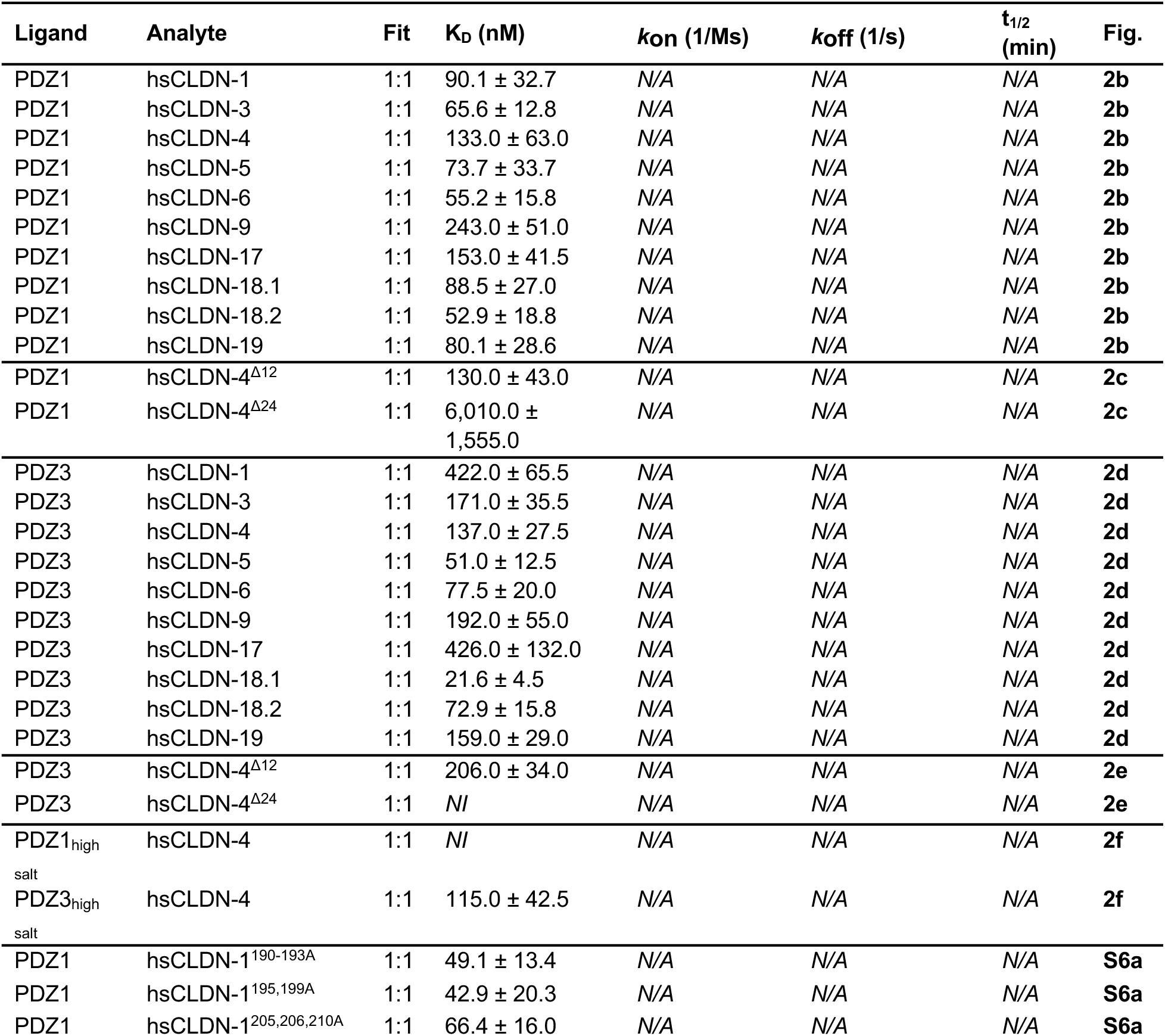
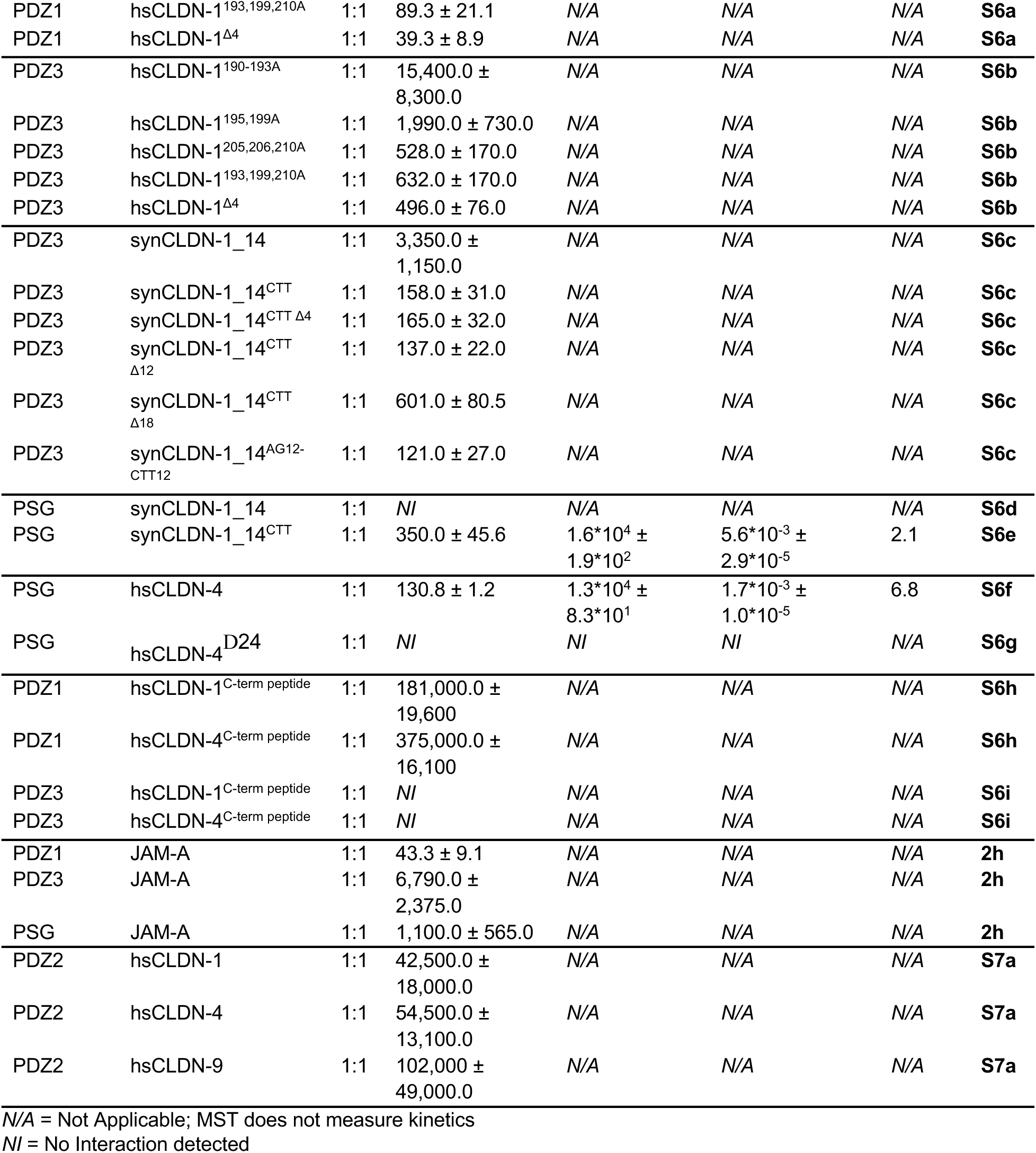
Claudin & JAM-A Binding Studies.

**Table 4.**
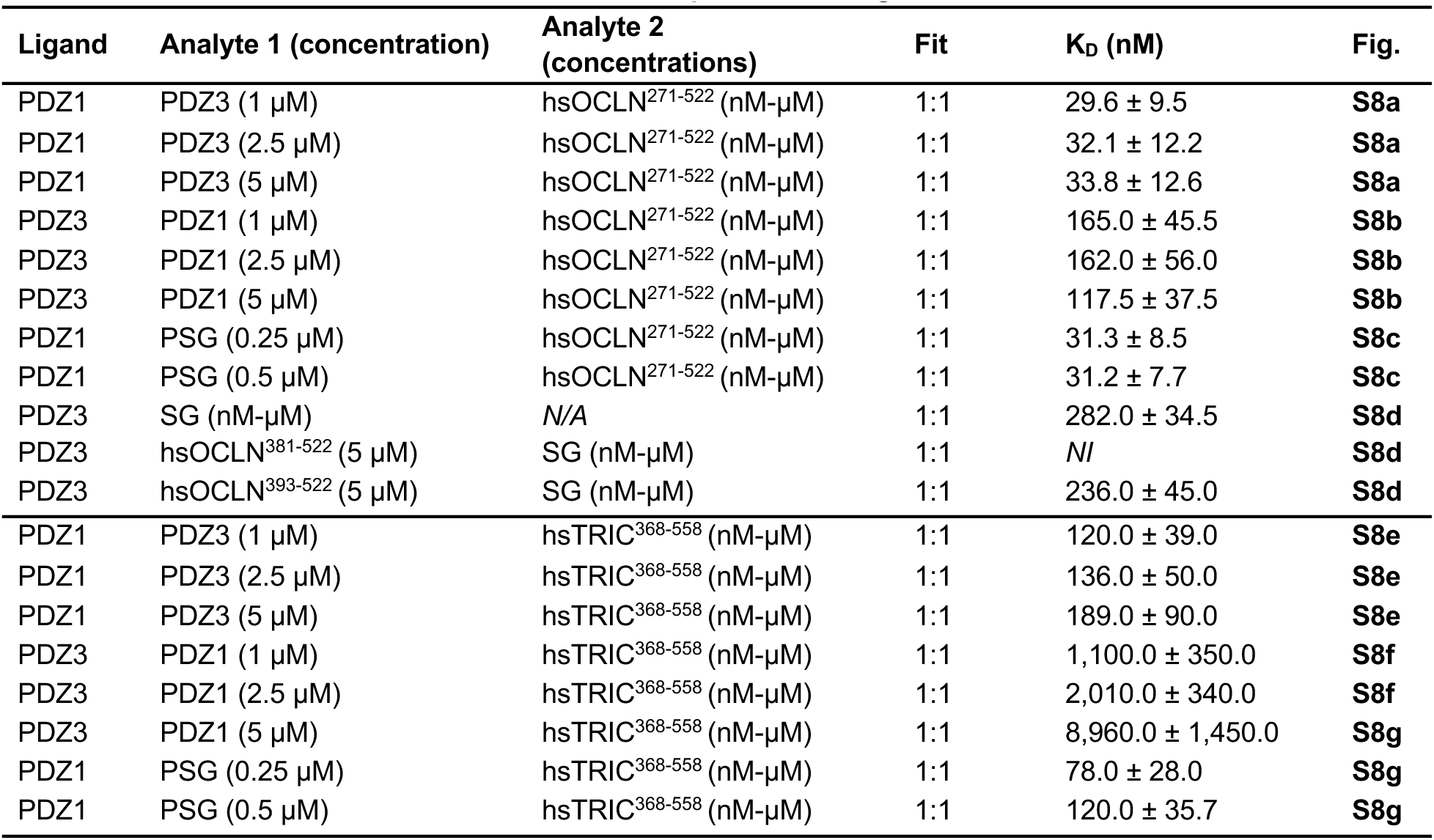
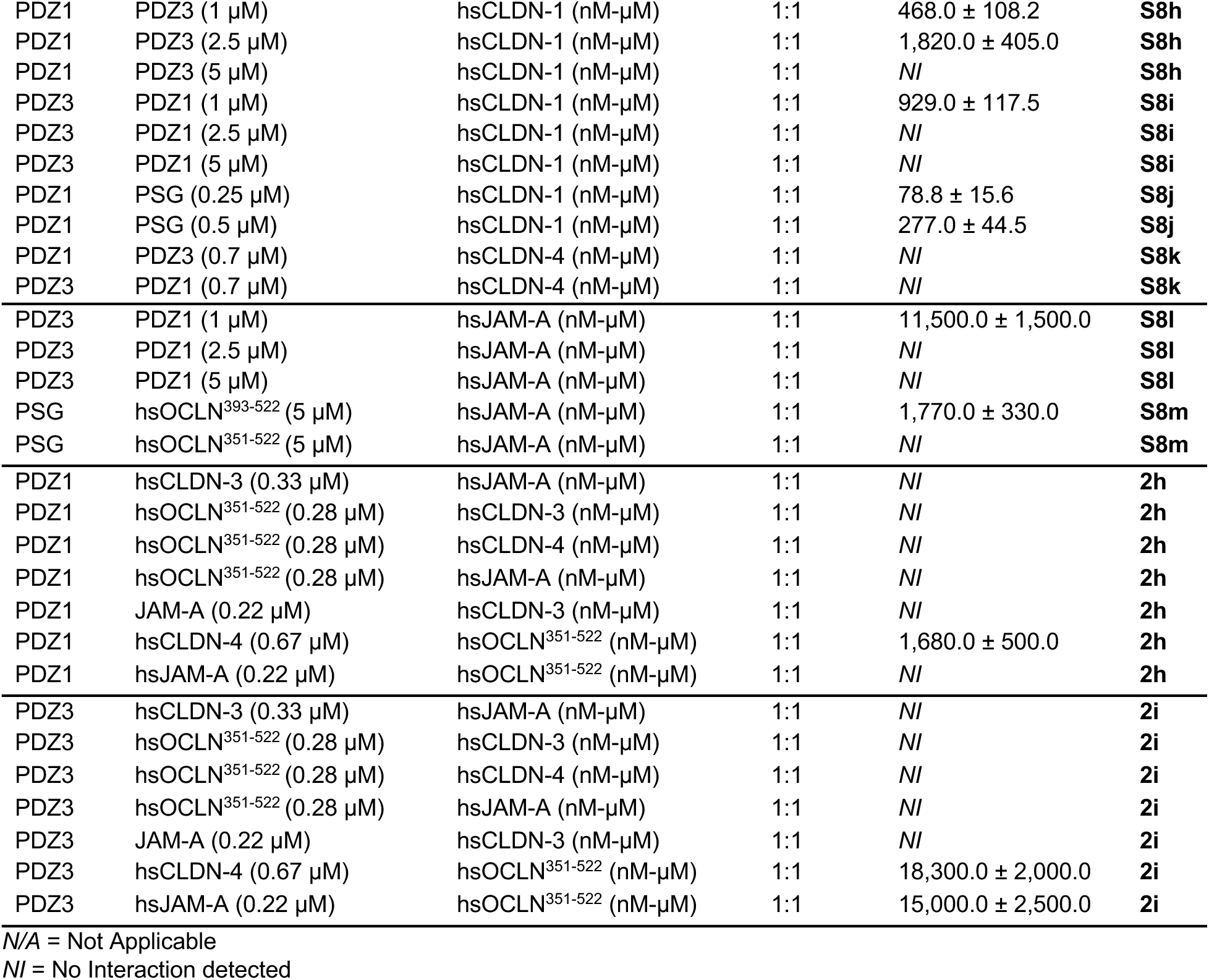
Three-Component Binding Studies.

We applied the same approach we used for TAMPs to resolve whether claudins also bind other PDZ domains of ZO-1. Like PDZ1, we found that PDZ3 bound to all 10 claudin subtypes, but also found that unlike PDZ1, there was a greater range of affinities between 21.6 to 426.0 nM (**Fig. 2d** and **Table 3**). Using claudin-4 truncations, we found that full-length and Δ12 claudin-4 bound PDZ3 with 137.0-206.0 nM affinity, but that the Δ24 truncation did not bind (**Fig. 2e** and **Table 3**). We tested again whether high salt affected PDZ1 and PDZ3 binding, this time to claudins, and found that 500 mM NaCl prevented claudin-4 binding to PDZ1 but only altered its affinity to PDZ3 1.2-fold compared to low salt (**Fig. 2f** and **Table 3**). These results established that the PDZ1 and PDZ3 domains of ZO-1 bind claudin C-terminal domains interchangeably as they do for TAMPS using a similar mode but unique chemical mechanism. Importantly, they again showed that claudin PDZ motifs are not needed for binding to PDZs.

Our observation that PDZ motifs are not required for PDZ binding to claudins had not been demonstrated before, so we tested this in another claudin subtype by creating: 1) mutants of claudin-1 C-termini at potential phosphorylation sites or deletion of its four-residue PDZ motif; and 2) a synthetic soluble analog of claudin-1 with no C-terminus, native C-terminus, or truncations to native C-terminus (**Fig. 2g**). First, we made four mutants with Ala substitutions to Ser, Thr, and Tyr residues in three regions along claudin-1’s C-terminus and one truncation of the last four residues (ΔKDYV—hsCLDN-1^Δ4^) constituting the PDZ motif. We tested binding of these mutants to labeled PDZ1 using MST and found that all claudin-1 variants bound with affinities that were slightly higher than wild type (**Fig. S6a** and **Table 3**). For PDZ3, we found that all claudin-1 variants bound with lower affinities than wild type. Specifically, the truncated hsCLDN-1^Δ4^ and phosphorylation site mutants hsCLDN-1^205,206,210A^ and hsCLDN-1^193,199,210A^ bound with 1.2-1.5-fold lower affinity compared to wild type claudin-1 (**Fig. S6b** and **Table 3**). However, the binding of mutants hsCLDN-1^190-193A^ and hsCLDN-1^195,199A^ to PDZ3 were significantly compromised with 36.5- and 4.7-fold lower affinities, respectively. The hsCLDN-1^190-193A^ variant mutations, which showed the largest change, were localized closer to TM4 than the PDZ motif (**Fig. 2g**). These results agreed with claudin-4 truncation data and validated that neither PDZ1 nor PDZ3 require the PDZ motif of claudins to bind. Additionally, our findings showed that identical changes to claudin C-terminal sequences affect PDZ1 and PDZ3 binding uniquely.

To validate further, we used soluble analogs of claudins designed in silico that we previously determined fold and function like native claudins.^<u>63</u>^ These analogs have their hydrophobic TM residues substituted with polar residues but have native ECS domain sequence and structure with no C-termini. We engineered the claudin-1 analog synClaudin-1_14 to have a native C-terminus (synClaudin-1_14^CTT^) then made variants of variable length (**Fig. 2g**) and tested their binding to PDZ3 via MST. We found that PDZ3 bound synClaudin-1_14 weakly with an affinity of 3,350.0 nM but bound with an affinity of 158.0 nM to synClaudin-1_14^CTT^ (**Fig. S6c** and **Table 3**). This indicated that the native C-terminus of claudin-1 directed binding. We then found that truncations of 4 (synClaudin-1_14^CTTΔ4^) and 12 (synClaudin-1_14^CTTΔ12^) amino acids had no effect on PDZ3 binding and only that an 18 (synClaudin-1_14^CTTΔ18^) amino acid truncation, which bound with 601.0 nM affinity, compromised PDZ3 binding ability (**Fig. S6c** and **Table 3**). Another mutant where the last 12 residues were native and the 7 upstream near TM4 were mutated to random amino acids (synClaudin-1_14^AG12_CTT12^) also bound PDZ3 with native-like affinity. Lastly, we tested synClaudin-1_14’s and synClaudin-1_14^CTTΔ4^’s abilities to bind PSG, which has a PDZ3. We found that synClaudin-1_14 failed to bind strongly even at 2,500 nM to PSG but that synClaudin-1_14^CTTΔ4^ with its PDZ motif removed, bound well with a K_D_ of 350.0 nM (**Fig. S6d-e** and **Table 3**). This was similar to the 422.0 nM affinity measured for wild type claudin-1 binding to PDZ3 (**Fig. 2d**). We also tested PSG binding to claudin-4 and hsCLDN-4^Δ24^ and found that the former bound with 130.8 nM but the latter did not bind (**Fig. S6f-g** and **Table 3**). These results indicated that there is length but not strong sequence dependency for PDZ binding and demonstrated clearly that the PDZ motifs of claudins are not required for binding by ZO-1 PDZs.

### Claudin transmembrane domains aid PDZ binding to their disordered C-termini

Because PDZ motifs of claudins were thought to be required for PDZ binding and our evidence countered this idea, it instigated us to reproduce what others had done to establish the precedent. The crystal structure of claudin-1’s C-terminal peptide SSGKDYV bound to PDZ1 was determined by fusing the peptide onto the C-terminus of PDZ1.^<u>20</u>^ The authors noted that co-complexing claudin C-terminal peptides that were unfused to PDZ1 failed to yield peptides bound to PDZ1 in crystals, indicating a very low affinity. We synthesized peptides encoding the C-termiil of claudin-1 (PRKTTSYPTPRPYPKPAPSSGKDYV) and −4 (PPRTDKPYSAKYSAARSAAASNYV) and tested binding to PDZ1 and PDZ3. We found that PDZ1 bound claudin-1 and −4 peptides with 181,000 and 375,000 nM affinities, respectively, but that PDZ3 did not bind either at even millimolar concentrations (**Fig. S6h-i** and **Table 3**). These binding affinities represented >2,000-fold decreases in affinity to PDZ1 and PDZ3 compared to native claudins (**Figs. 2b,d**). Data from that study found that unfused claudin peptides bound to PDZ1 in solution with >13,500 nM affinity, which agrees with our data.^<u>20</u>^ This experiment demonstrated that unstructured peptides from PDZ-binding receptors may be poor proxies to understand how the native receptors interact with PDZ domains. In biophysical context, our results demonstrate that the structured TM domains of membrane proteins like claudins aid the ability of PDZs to bind the largely disordered C-termini of receptors, which explains our finding that PDZ1 and PDZ3 bind TM4 proximal regions of claudins.

### JAM-A binds PDZ domains like a claudin, not like a TAMP

JAM-A is a single TM spanning TJMP that has a PDZ motif at the end of its intracellular C-terminus and is known to regulate the transport of large molecules through the tight junction paracellular barrier.^<u>64</u>^ It was thought that JAM-A’s PDZ motif binds PDZ3 and SH3 of PSG based on a structure of a C-terminal JAM-A peptide (EGEFKQTSSFLV) in complex with PSG.^<u>21,65</u>^ In that study, the JAM-A peptide, which constituted only 12 of 36 C-terminal residues, bound only to PSG with a K_D_ of 35,000 nM and not PDZ3, which suggested a role for the SG domain in binding. Because these findings have influenced subsequent understanding of JAM-A/ZO-1 complexes, and we showed the limits of using peptides to study PDZ interactions, we sought to reproduce or challenge this model. We fluorescently labeled PDZ1, PDZ3, and PSG, and tested binding against full-length JAM-A using MST. We measured an affinity of 43.3 nM for PDZ1 binding to JAM-A, while for PDZ3 and PSG we measured affinities of 6,790 nM and 1,100 nM, respectively (**Fig. 2h** and **Table 3**). Our data that JAM-A binds better to PSG than PDZ3 agreed with previous work, but our observation that it binds 25.4-fold better to PDZ1 than PSG had not been demonstrated. These results showed that JAM-A interactions with PDZs resemble claudins more than TAMPs, which bind best to PSG, and in full when combined with our other findings, provide the selective binding modes of TJMPs to three major PDZ-containing domains and modules of ZO-1.

### PDZ2 of ZO-1 has no relevant TJMP binding ability

Because PDZs are structurally homologous and PDZ1 and PDZ3 had overlapping TJMP binding abilities, we tested whether PDZ2 performed similarly (**Fig. S3**). It is thought that PDZ2’s primary function is to direct ZO-1 oligomerization through its own dimerization.^<u>46</u>^ Here, we fluorescently labeled PDZ2 and tested binding to the longest C-terminal domains of occludin and tricellulin, and claudins −1, −4, and −9. We found that PDZ2 bound to claudins very weakly with 42,500-102,000 nM affinity (**Fig. S7a** and **Table 3**). We found no binding of PDZ2 to hsOCLN^271-522^ and for tricellulin we found PDZ2 bound both C-terminal domain constructs but poorly, with 12,500-15,600 nM (**Figs. S7b-c** and **Table 2**). SEC validated that the PDZ2 being tested, like PDZ1 and PDZ3, was monomeric in solution, which indicated that weak binding was not a result of PDZ2 dimerization blocking access (**Fig. S2b**). Our findings that PDZ2 forms complexes with PDZ1 and PDZ3, coupled with PDZ2s weak affinity for claudins, hinted that intra-molecular interplay within ZO-1 PDZ domains could regulate their interactions with TJMPs.

### PDZs bind disparate sites on occludin but overlapping sites on other TJMPs

After establishing the biophysical rules of two-component binding interactions between sub-domains of ZO-1 and TJMPs, we tested the competitiveness of these interactions using three-component binding assays and MST. For these assays, a labeled ligand was pre-incubated with an unlabeled analyte at a single concentration (analyte 1) that exceeded its K_D_, then this mixture was incubated with another unlabeled analyte at increasing concentrations (analyte 2). The binding of analyte 1 to the ligand may or may not alter or prevent binding of analyte 2, and binding and affinity can be compared to two-component data to see (**Tables 2-3**).

First, we tested the ability of hsOCLN^271-522^ to bind PDZ1 or PDZ3 in the presence of increasing concentrations of the other. We found that PDZ3 had no effect on PDZ1 binding whereas PDZ1 had a marginal effect of PDZ3 binding—in the latter the affinity decreased ∼2.0-fold (**Figs. S8a,b** and **Table 4**). Non-competitive binding of PDZ1 and PDZ3 indicated non-overlapping binding sites on occludin. We next tested PDZ1 binding in the presence of PSG and found, like for PDZ3, that PSG had no effect of PDZ1 binding (**Fig. S8c** and **Table 4**). We then established that PDZ3 binds SG to form a PSG complex in a two-component assay where the two sub-domains bound with an affinity of 282.0 nM (**Fig. S8d** and **Table 4**). In the presence of hsOCLN^393-522^, which constitutes the CCD, PDZ3 binding to SG was not affected but with hsOCLN^381-522^ in solution, PDZ3 could not bind SG (**Fig. S8d** and **Table 4**). This indicated that PDZ3/SG complexation to form PSG can be altered or inhibited by occludin’s C-terminal domain.

We performed similar analyses of tricellulin using hsTRIC^368-558^. However, we found that PDZ3 attenuated PDZ1 binding moderately by decreasing affinity to hsTRIC^368-558^ 3.2-fold, and that PDZ1 significantly inhibited PDZ3 binding to hsTRIC^368-558^ up to 32.9-fold compared to PDZ3 alone (**Figs. S8e,f** and **Table 4**). We then tested PDZ1 binding to hsTRIC^368-558^ with PSG present and found that it decreased PDZ1 binding 2.0-fold (**Fig. S8g** and **Table 4**). Competitive binding of PDZ1 and PDZ3 indicated that they bind overlapping or partially overlapping regions of tricellulin. These results differed from occludin.

After analyzing TAMPs, we tested binding of PDZs to claudin-1. We found that PDZ3 prevented PDZ1 from binding claudin-1 in a concentration dependent manner and completely inhibited binding above 2,500 nM (**Figs. S8h** and **Table 4**). PDZ1 was a more effective inhibitor of PDZ3 binding to claudin-1 (**Figs. S8i** and **Table 4**). We also found that PSG at concentrations 10-fold lower than PDZ3 inhibited PDZ1 binding to claudin-1 and decreased their affinity 3.1-fold (**Fig. S8j** and **Table 4**). Because claudins have variable C-termini, we also tested competition with claudin-4 and found that both PDZ1 and PDZ3 in excess could block binding of the labeled PDZ to claudin-4 (**Fig. S8k** and **Table 4**). These results suggested that PDZ1 and PDZ3 compete for binding to claudin C-termini via overlapping binding sites.

Next, we performed experiments on JAM-A, which we showed binds PDZ1 much better than PDZ3 (**Fig. 2h**). We therefore did not test binding of PDZ1 with excess PDZ3 but did the inverse experiment. We found that PDZ1 at 1,000 nM inhibited PDZ3 binding to JAM-A and that at concentrations >2,5 00 nM, PDZ3 was unable to bind (**Fig. S8l** and **Table 4**). This result agreed with our finding that JAM-A binds PDZs more like a claudin but was surprising given JAM-A’s longer C-terminus compared to most claudins. Because JAM-A bound PDZ3 with low micromolar affinity we suspected it would not alter high affinity PSG binding to occludin even though JAM-A binding to PDZ3 is improved by SH3 (**Fig. 2h**). Indeed, we found that excess hsOCLN^393-522^ CCD had no effect on JAM-A binding to PSG but that the longer hsOCLN^351-522^ inhibited it completely (**Fig. S8m** and **Table 4**). These results showed that PDZ1 and PDZ3 compete for an overlapped binding region on JAM-A’s C-terminus and that JAM-A binding to PDZ3 cannot occur in the presence of occludin with a native length C-terminal domain.

### Only one TJMP can bind a single PDZ simultaneously

Our three-component competitive binding studies instigated us to design experiments to test whether one or more TJMP could bind a single PDZ domain simultaneously. These experiments were required because occludin had non-overlapping PDZ binding sites whereas PDZ binding sites partially or fully overlapped for tricellulin, claudins, and JAM-A. The results for occludin could be explained either by a single PDZ having two unique binding sites for a ligand, or by a single ligand having two unique and non-overlapping binding sites where different PDZs could bind. We hypothesized that the latter were true due to the differential effect of salt on PDZ1 and PDZ3 interactions with TJMP ligands (**Figs. 1q,r,t** and **2f**). Here, we pre-incubated an unlabeled TJMP (analyte 1) at a single concentration with labeled PDZ1 or PDZ3 then titrated increasing concentrations of a different unlabeled TJMP (analyte 2) and measured binding using MST. If the latter TJMP can bind, we would expect concentration dependent binding curves. We found that the presence of a claudin, occludin, or JAM-A completely inhibited the ability of another TJMP to bind PDZ1 (**Fig. 2i** and **Table 4**). Only very compromised binding (30.3-fold decrease) of hsOCLN^351-522^ to PDZ1 was detected in the presence of claudin-4. For PDZ3, we found the same, that a TJMP completely inhibited the ability of another to bind (**Fig. 2j** and **Table 4**). And again, we measured very weak binding of only hsOCLN^351-522^ to PDZ3. This finding was consistent with other data that showed that hsOCLN^351-522^ binds PDZs with high affinity (**Figs. 1h,r**). These results verified that occludin has two distinct PDZ binding sites and that a single PDZ domain has only one ligand binding site.

### Mapping of the ZO-1/TJMP interactome

In total, our biophysical studies, for the first time, quantitatively and nearly completely maps ZO-1’s intracellular interaction network with TJMPs to allow for elucidation of the hierarchy of binding partners and the principles that form and govern the ZO-1/TJMP interactome. We provide a summary of this network, along with its intrinsic hierarchies, in **Fig. 3**.

**Fig. 3.**
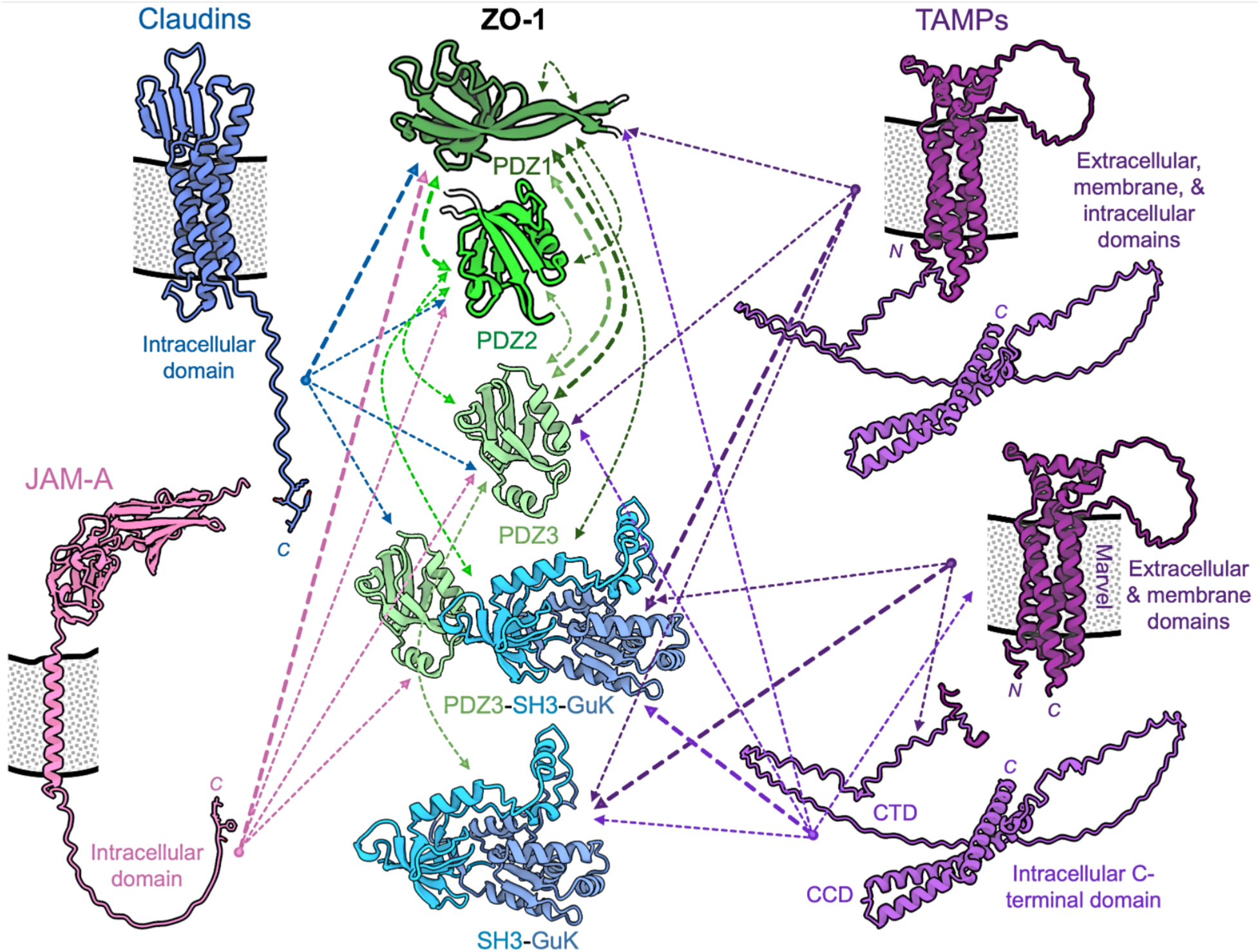
Updated model of the ZO-1/TJMP interactome. Schematic of intra- and inter-molecular interactions between ZO-1 and TJMPs elucidated by this study. Proteins are shown as cartoons based on AlphaFold^<u>95</u>^ predictions and colored as in Fig. 1a. Dashed lines represent experimentally verified biophysical interactions, with line thickness correlating to the primary highest affinity or longest-lived interactions.

### Rationale for phase separation studies of TAMPs and PSG

After establishing that TAMPs are the primary binders of the PSG module of ZO-1, we aimed to provide greater context to our biophysical studies by characterizing other physical behaviors. Because TAMPs have extensive IDRs and we determined that they employ multivalency to bind all sub-domains of PSG, we designed experiments to test whether TAMPs phase separate, as these traits are hallmarks of phase-separating proteins (**Figs. 1e,f, S4a-i** and **S5**). It had been shown previously that ZO-1 phase separates within cells, the PSG module is required for this physical transformation, phase separation is inhibited by PSG intra-molecular interactions with the U6 region downstream of it, and ZO-1 phase separation recruits TJMPs to condensates^<u>66-69</u>^. We therefore hypothesized that PSG binding proteins like TAMPs could modulate phase separation of ZO-1 to in turn modulate recruitment and coalescence of TJMPs.

In biophysical studies, we found that PSG interacts with occludin and tricellulin using their C-terminal domains. For occludin, residues 271-393 were required for high affinity interactions (**Fig. 1g**) while for tricellulin, high affinity interactions only require residues 424 onward (**Figs. S4k-m**). Other data showed that PDZ3 binding to the 271-360 region of occludin helped explain this difference (**Fig. 1h**). We used structural predictions that showed both TAMPs have IDRs between their folded Marvel and CCD domains and performed alignments that revealed the length and sequence of each TAMP’s IDR vary, with occludin having 72 residues more than tricellulin with one large 43 residue insert between residues 347-390 (**Figs. 4a** and **S5**). Given that the region PSG binds differed between occludin and tricellulin, and substantial sequence changes exist, we focused additional biophysical studies on TAMP C-terminal domains and specifically occludin’s IDR insert.

**Fig. 4.**
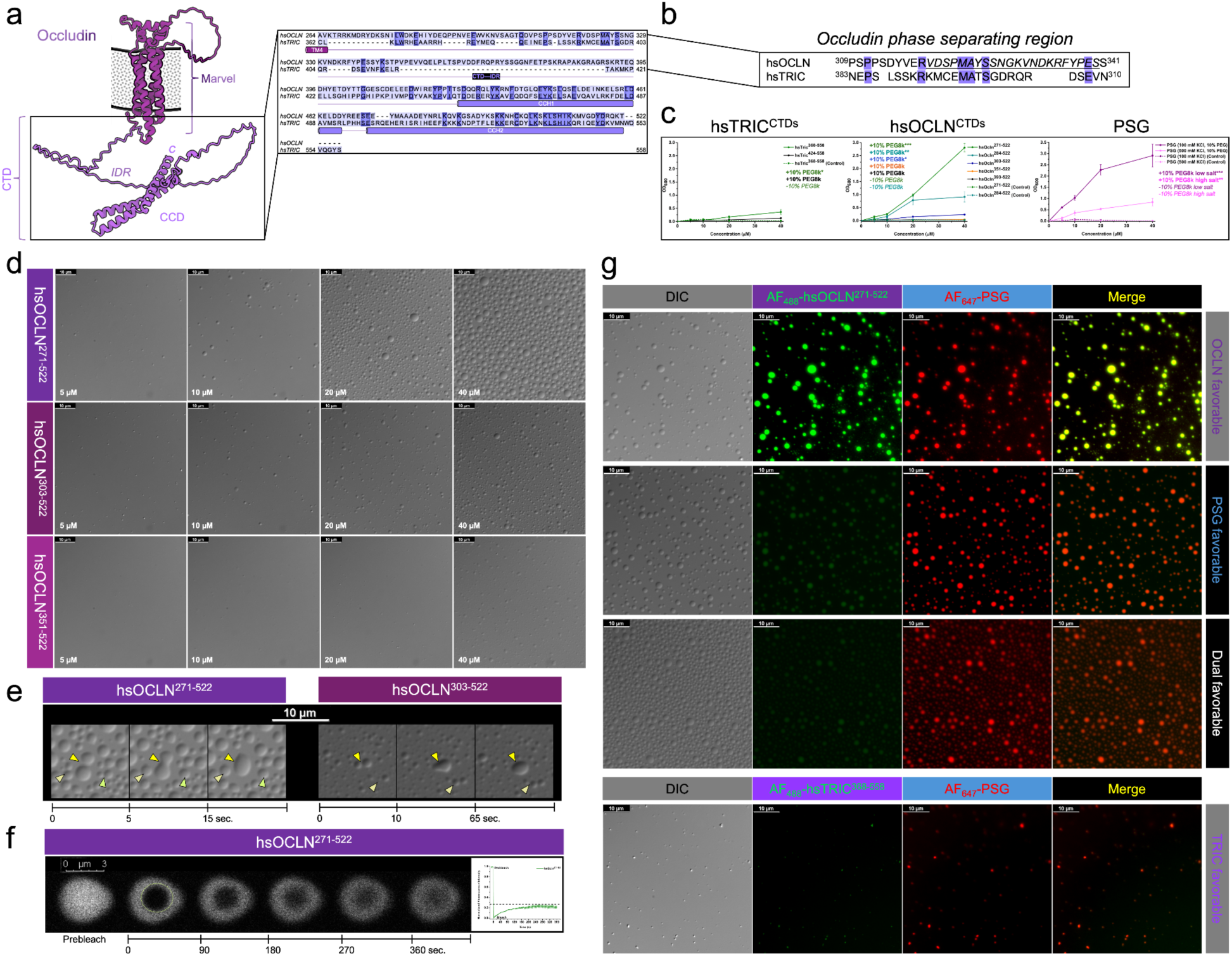
Phase separation of occludin and PSG. (a) Structural prediction of occludin depicting the extracellular and membrane-embedded Marvel domain and intracellular C-terminal domain (CTD) with IDR that includes the coil-coil domain (CCD); along with a sequence alignment of the intracellular domains of occludin and tricellulin. Note the 48 amino acid insertion in occludin that gives it a longer IDR. The coil-coil helix 1 (CCH1) and 2 within the CCD are highlighted in the sequence, as is TM4. (b) Intracellular sequences of occludin and tricellulin in the region predicted by ParSe to phase separate. Phase separating residues are italicized and underlined. Note that homology exists between both TAMPs here but only occludin is predicted to phase separate. (c) Turbidity assays of occludin and tricellulin CTDs benchmarked against the known phase separating protein PSG. Controls without phase separation molecules are shown as -% PEG 8k. * denotes relative turbidity quantity. (d) Phase separation abilities of three variable length occludin CTD constructs. Note that the shortest construct lacking the IDR and phase separating region only forms droplets at very high concentrations. (e) Droplet fusion studies using time-lapse DIC microscopy of two occludin CTDs. Fusion events are shown in triangles. (f) Time-lapse fluorescence recovery of a labeled hsOCLN^271-522^ droplet over 360 seconds showing liquid-liquid behavior but incomplete recovery. (g) Cohabitation after mixing of occludin CTD and PSG fluorescently labeled condensates. Occludin and tricellulin CTDs were labeled with alexa fluor 488 and PSG with alexa fluor 647 then mixed in buffers favorable to each and analyzed with fluorescence microscopy. Merged images show that PSG mixed with occludin CTD cohabit the same droplets while PSG mixed with tricellulin CTD we observe PSG droplets that sequester small amounts of tricellulin CTD within.

### The intracellular C-terminal domain of occludin, but not tricellulin, phase separates

First, we used bioinformatics to predict phase separation of TAMPs using the ParSe servers.^<u>70,71</u>^ ParSe predicted that two regions of occludin and three regions of tricellulin phase separated (**Fig. S9**). In the C-terminal domain, however, only occludin residues 319-340 were predicted to phase separate (**Fig. 4b**). Interestingly, the phase separating region of occludin was upstream and not contained within the IDR insert. Next, we tested TAMP phase separation using in vitro turbidity assays of our various TAMP C-terminal domain constructs by mixing them at various concentrations in phase separation-inducing buffer and measuring OD_600nm_.^<u>72,73</u>^ We found that turbidity of the two tricellulin constructs did not or minutely increased upon increasing concentrations up to 40 µM, whereas for occludin, the three longest constructs containing residues 271-350 increased turbidity dramatically and in concentration dependent ways (**Fig. 4c**). Importantly, without phase separation additive, occludin did not increase turbidity. We then benchmarked this result against PSG, a known phase separating protein, and found that PSG increased turbidity to the level of the longest occludin C-terminal domain construct, hsOCLN^271-522^ (**Fig. 4c**). PSG turbidity increased in low salt. In high salt, PSG turbidity was compromised and in no additive buffer no turbidity was detected. These results suggested that occludin and PSG, but not tricellulin, phase separate in vitro. However, because turbidity is not directly correlated to phase separation and could be attributed to protein aggregation or precipitation, we investigated this phenomenon further.

We next visualized the microscale physical properties of occludin using differential interference contrast (DIC) microscopy. First, we screened nine phase separation-inducing buffers for their ability to form droplets of the longest C-terminal domains of occludin (hsOCLN^271-522^) and tricellulin (hsTRIC^368-558^), and of our control, PSG, at 10 µM. We found robust droplet formation of hsOCLN^271-522^ in seven conditions that spanned low and high salt and precipitating polyethylene glycol 8,000 (PEG 8k) concentrations (**Fig. S10a**). For hsTRIC^368-558^, we found only two conditions yielded physical changes and that these droplets did not resemble the shape or size of droplets formed by occludin (**Fig. S10b**). For PSG, we found all nine conditions yielded physical changes, with droplets forming preferentially at high salt and PEG 8k concentrations (**Fig. S10c**). We attributed the alternate shaped forms at low salt, which are clustered and not spherical, to be aggregates. These results showed that the conditions at which phase separating droplets form and the resulting number, size, and morphology of these droplets were unique for occludin, tricellulin, and PSG but also suggested again that occludin and PSG phase separate in vitro.

Now that phase separating conditions were isolated and because only occludin C-terminal domain constructs containing residues 271-350 increased turbidity, we tested the abilities of three variable length occludin C-terminal domains to phase separate and compared them to PSG. First, we found that PSG formed droplets at >1 µM and did so robustly at 10 µM, which agreed with previous results using 5 µM (**Fig. S11a**).^<u>66</u>^ We then found with hsOCLN^271-522^, hsOCLN^303-522^, and hsOCLN^351-522^, that only the two longest constructs phase separated well above 5 µM (**Fig. 4d**). Full concentration screening showed that hsOCLN^271-522^ and hsOCLN^303-522^ formed droplets between 1.0-2.5 µM, whereas it required >20 µM of hsOCLN^351-522^ to form droplets (**Figs. S11b-d**). In absence of PEG 8k, no phase separation occurred for any construct, even at 40 µM. These results verified that PSG and occludin’s C-terminal domain phase separate and that residues 271-350 of occludin enable this physical transition.

### Occludin droplets form liquid-liquid condensates

To better classify the phase separation behaviors and physical states of hsOCLN^271-522^ and hsOCLN^303-522^ formed under these conditions we used time-lapse DIC microscopy and fluorescence recovery after photobleaching (FRAP). After forming droplets, we monitored droplet fusion and found that two independent drops fused into single larger droplets within 5 seconds for hsOCLN^271-522^, while for hsOCLN^303-522^, similar fusion occurred in 10 seconds (**Fig. 4e**). We photobleached droplets formed by fluorescently labeled hsOCLN^271-522^ and found that fluorescence recovered by 6 minutes, but that recovery was only ∼25% complete (**Fig. 4f**). The fusion of droplets and recovery of fluorescence within droplets suggested that occludin’s C-terminal domain undergoes liquid-liquid phase separation.

### PSG and occludin’s C-terminal domain cohabit phase separated droplets

Given that our biophysical studies showed that PSG interactions with the occludin C-terminal domain are high affinity but have short half-lives of <5 minutes, and that the isoelectric points of occludin’s C-terminal domain is 5.5 vs. 9.6 for PSG, we predicted that PSG and occludin phase separation was associative and driven by complimentary electrostatics. We tested whether PSG and hsOCLN^271-522^ labeled with unique fluorophores would cohabit the same drop and mix; partition separately within a single drop; or form homogeneous droplets that did not mix. In a buffer favorable to hsOCLN^271-522^ phase separation, we found that droplets formed, and that droplets were cohabited with both hsOCLN^271-522^ and PSG (**Fig. 4g**). In a buffer that was favorable for PSG or the phase separation of both proteins, we found the same, although the amount of hsOCLN^271-522^ contained in each droplet decreased despite there being a greater number of droplets under these conditions (**Fig. 4g**). To test whether cohabiting droplets was exclusive to occludin/PSG, we performed the same analysis using hsTRIC^368-558^, which we showed did not phase separate alone (**Fig. S10b**). The resulting droplets resembled those of PSG alone (**Figs. S10c** and **S11a**), which are small and infrequent (**Fig. 4g**). However, small amounts of fluorescently labeled hsTRIC^368-558^ appear in PSG driven condensates, indicating that PSG may sequester non-phase separating tricellulin to droplets, as was shown previously for claudins, JAM-A, and occludin.^<u>66,69</u>^ Experimental controls where only one protein is labeled in various buffers, both proteins labeled in different buffers, and their resulting microscopy images were also visualized for experimental robustness (**Figs. S12-15**). In full, our phase separation studies demonstrate that despite their conserved topology and homology, the intracellular C-terminal domain of occludin, but not tricellulin, phase separates, and that the droplets it forms coalesce and integrate into PSG droplets to establish a greater associative liquid-liquid condensate.

### Phase separation may be a select but prominent property of TJMPs

After demonstrating that occludin phase separates using residues 271-350, which were predicted by ParSe to contain the phase separating residues 319-340, we used both ParSe servers to predict the phase separation behaviors of other TJMPs (**Tables 5** and **S1)**. We found that claudins −3, −6, −15, and −18 contained regions in their intracellular C-termini that may phase separate, but no other claudin did. Moreover, we found that JAM-B and JAM-C may have phase separating regions that reside on opposite sides of the membrane and that JAM-A was not predicted to phase separate. For occludin and tricellulin, the predictions showed ECS1 phase separates for both TAMPs, which would be a conserved feature because only occludin was predicted to phase separate using its C-terminus and only tricellulin using its N-terminal domain—both domains reside inside cells. Interestingly, the other TAMP family members, two Marvel-D3 splice variants, which are known to interact with occludin and tricellulin at bi- and tri-cellular contacts, respectively, were also predicted to phase separate.^<u>55,74</u>^ These splice variants have 198 residues at their N-termini 100% conserved but share only 29% conservation in their remaining ∼200 residues. We found that Marvel-D3s were predicted to phase separate using a conserved region within their intracellular N-terminal domains but also uniquely with D3-1 residues 391-410 of its intracellular C-terminus and with D3-2 residues 222-250 of its ECS. This ParSe bioinformatic predicted behavior, strikingly, appears to be quite accurate. Because, in addition to the previously demonstrated phase separation of ZO-1^<u>66</u>^, and our data validating the phase separation of occludin’s C-terminal domain (**Fig. 4**), it has also been shown recently that occludin’s ECS1 phase separates (personal communication with <u>Martin Lehmann</u>). These results strongly validate ParSe’s ability to predict phase separation of TJMPs. Taken together, the predictions reveal that TJMPs may have unique phase separating regions and that closely related family members, such as TAMPs and JAMs, may have regions that activate or counteract the phase separation behaviors of other family members as they often reside on other sides of the membrane or opposite termini. In full, our analyses suggest that phase separation may be a fundamental property of TJMPs that may function to couple intracellular to extracellular signaling through membranes.

**Table 5.**
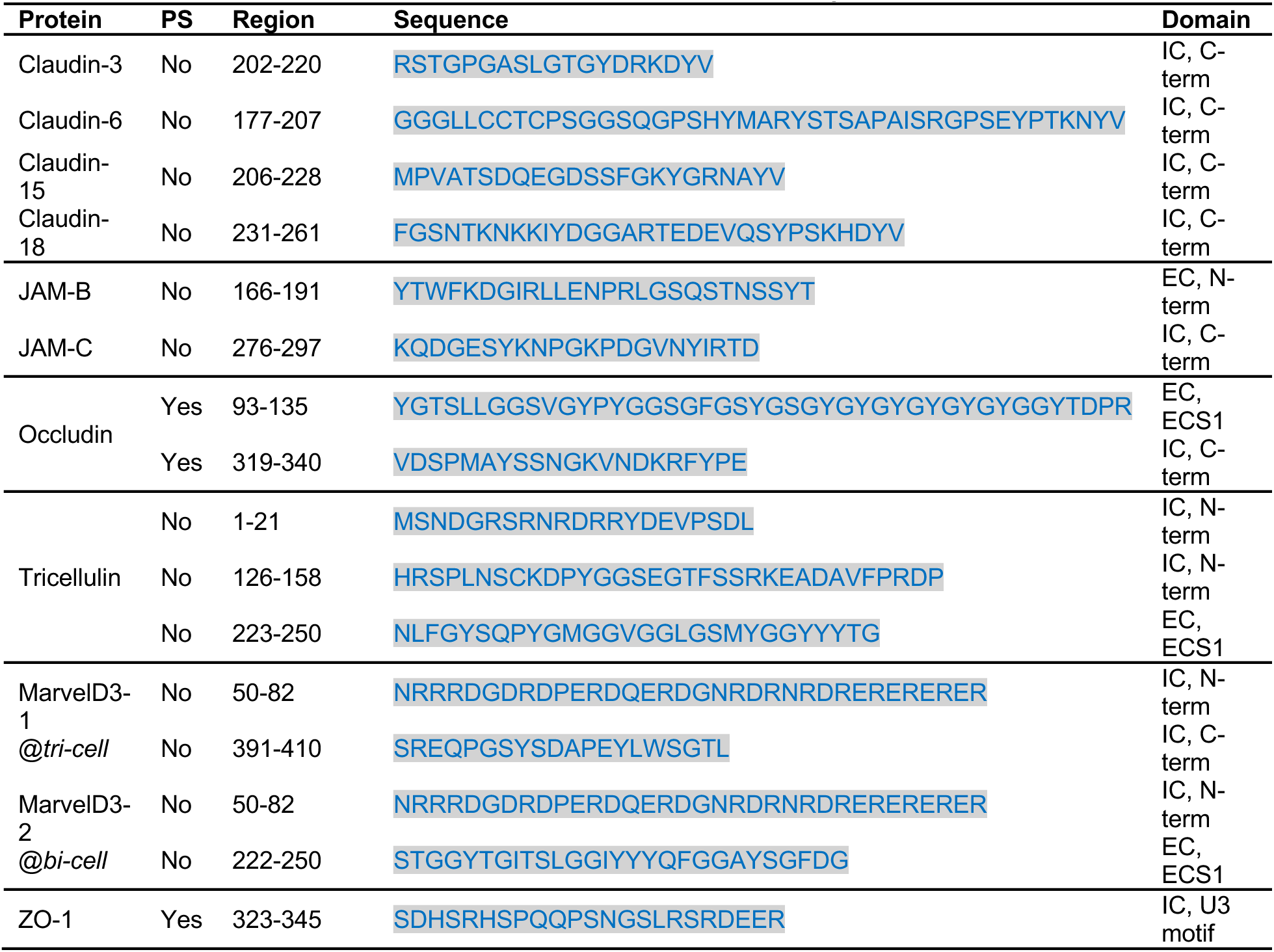
TJMPs Predicted to Phase Separate.

### Homology between ZO-1’s Unique motifs and TAMP C-terminal domains provides a mechanism to regulate phase separation

Between the structured domains of ZO-1 are unstructured unique (U) motifs with highly variable sequences and lengths (**Fig. 1a**). Two U motifs are potentially relevant to regulation, U5 between SH3 and GuK, which helps bind occludin and localize ZO-1 to tight junctions; and U6 downstream of PSG, which negatively regulates ZO-1 phase separation through intra-molecular interactions with PSG and at larger scales results in displaced tight junctions that contain ZO-1, occludin, and claudins when removed from ZO-1.^<u>22,56,66</u>^ Phosphorylation of residues in U5, U6, and occludin’s C-terminal domain are also known to regulate this system.^<u>22,37</u>^ Based on this knowledge, we hypothesized that unidentified sequence elements within these regions must exist that function to tune intra- and inter-molecular interactions. We performed sequence alignments between U5 and U6 individually against each of the C-terminal domains of occludin and tricellulin (**Fig. S16**). For occludin we found 22.2% sequence identity with U5 and 29.7% identity with U6 (**Fig. S16a,c**). Interestingly, both U motifs aligned well with occludin residues known to be phosphorylated. Moreover, U5 aligned to the IDR and CCD regions of occludin, while U6 aligned primarily to occludin’s IDR. For tricellulin, we found 25.8% sequence identity with U5 and 23.9% identity with U6 (**Fig. S16b,d**). Unlike occludin, only U6 aligned with tricellulin phosphorylation sites. Additionally, U5 aligned primarily to tricelluin’s TM4 proximal region while U6 aligned to TM4 proximal, IDR, and CCD regions. Lastly, because we showed that high salt abrogates occludin binding to PSG but not PDZ3 (**Figs. 1s,v** and **S4p,q**), we calculated isoelectric points (pI) of TAMP and PSG sub-domains (**Fig. S16e**). We found that global charge was highly correlated with the charges of the smaller IDR and U motifs. Specifically, occludin’s C-terminal domain is acidic with a pI of 5.5 largely due to its IDR region, whereas tricellulin’s C-terminal domain is basic with a pI of 8.7 due to its equivalent IDR. Similarly, PSG had a pI of 9.6 due to U5 having a pI of 11.3. U6 is very acidic, with a pI of 3.8. These analyses reveal previously hidden homology between ZO-1 U motifs and TAMP C-terminal domains, demonstrate how the IDR regions of TAMPs and U motifs of ZO-1 influence global protein chemistry, and suggest a mechanism for regulation of intra- and inter-molecular interactions between ZO-1 and TAMPs that at scale, controls tight junction organization.

## Discussion

Quantifying Interactions of MAGUK family proteins or their associated PDZ domains with ligand receptor proteins is notoriously challenging, thus comprehensive biophysical studies were mostly non-existent, leaving major gaps in our understanding of the many vital biological systems that employ these complex interaction networks. Here, for the first time, we quantified the intra- and inter-molecular interactions between ZO-1’s sub-domains and full-length, truncated, and mutated forms of the TJMPs claudins, occludin, tricellulin, and JAM-A to provide new insights into the system that operates at tight junctions. We found that the PDZ domains of ZO-1 exhibit intra-molecular interplay where PDZ1 binds all other PDZs while PDZ3 is more selective and cannot self-associate. We also discovered intra-molecular interplay within TAMPs, where the membrane-embedded Marvel domain interacts with the intracellular C-terminal domain to form multivalent ZO-1 binding hubs. Further, we revealed that inter-molecular interactions between claudins and PDZs do not require a PDZ motif and instead bind TM4 proximal regions that overlap where only a single PDZ can bind at once. Overlapping binding sites for PDZs also occur in JAM-A and partially for tricellulin, but not for occludin. We showed that occludin has distinct PDZ1 and PDZ3 binding sites and can bind both simultaneously. Moreover, we determined the unique mechanisms of ligand recognition of PDZs by showing that PDZ1 binds ligands promiscuously via electrostatics while PDZ3 binding is selective and not affected by charge. In full, our biophysical studies elucidate the selective binding hierarchy between ZO-1 and TJMPs where claudins and JAM-A compete for binding to PDZ1 while occludin, and to a lesser extent, tricellulin, are the primary binding partners for the PDZ3-SH3-GuK (PSG) module. We also show here, originally, that the intracellular C-terminal domain of occludin but not tricellulin undergoes liquid-liquid phase separation and that this physical transition is enabled by 21 residues. We characterize occludin phase separation to be associative and electrostatically complementary to PSG phase separation, where both condensates cohabit the same droplets. Lastly, we reveal how phase separation could be a prominent property of TJMPs and how it could be regulated by PSG binders like ZO-1’s own U5 and U6 motifs or newly identified U motif homologous regions within occludin and tricellulin. In total, we uncover the interactome between ZO-1 and TJMPs and elucidate the mechanisms that these proteins use to recognize, bind, assemble, and organize tight junctions at mesoscales, which we believe has relevance to signal transduction within and between epithelial cells.

Binding between ZO-1 and TJMPs had been previously described and a general outline for this network was used to interpret subsequent results and biology (**Fig. 1a**).^<u>16,18,21,23,31,34-36,45,47</u>^ However, the use of non-quantitative techniques or peptides obscured a full understanding of this system. Our biophysical interaction studies elucidate inter-molecular interactions between TJMPs and ZO-1 more completely and with higher precision and represent the first extensive and quantitative study of a MAGUK family protein with native full-length receptors that we know of.

Deciphering the intra-molecular interplay within ZO-1 and TAMPs is critical to understand how each initiate inter-molecular interactions as well as how each interaction type may be regulated. Our finding that PDZ1 binds all other PDZs and self-associates highlights its promiscuity, while our finding that PDZ3 only binds PDZ1, PDZ2, and SG shows it is selective and unable to self-associate. This suggests that PDZ1 or PDZ2 could bind to PDZ3 when it is bound to SG to form the PSG module. In the context of TJMP assembly, this provides a means for TAMP/PSG complexes to recruit claudin/PDZ1 complexes into larger assemblies. Indeed, it is known that ZO-1 recruits TJMPs and that assembly of TJMPs enriches ZO-1’s presence at tight junctions.^<u>66,69</u>^ Our work provides a potential mechanism for recruitment, whereby intra-molecular interactions between ZO-1 PDZ domains carry their bound TJMP cargo toward a coalesced TJMP/ZO-1 cluster. Moreover, our work ascribes a function for occludin’s long disordered C-terminal region that resides between its Marvel and CCD domains that was previously unknown. We show that occludin’s Marvel domain interacts with its C-terminal domain via this unstructured region, using residues 284-303. How the Marvel/C-terminal domain complex is formed is not understood by our work and could be the result of interactions between the intracellular loop connecting TM2 to TM3 or the ends of TMs themselves with the 284-303 region (**Fig. S5**). The intra-molecular interplay within TAMPs that we identify, coupled with our binding data, suggests that TAMPs form a multivalent interaction hub poised to bind ZO-1’s PSG module and explains how ZO-1 associates with membranes to recruit TJMP intracellular components into localized clustered assemblies.

The relevance of JAM-A binding to ZO-1 becomes complicated in light of previous studies and our data for only a single construct.^<u>21</u>^ Specifically, our results are limited by a construct that contains the full native protein with a five residue C-terminal protease scar downstream of the canonical PDZ motif. If a native PDZ motif is required for PDZ binding, this scar may alter native affinities. However, our binding data for JAM-A to PDZ3 and PSG agree with those from previous reports that show low affinities and that SH3 improves JAM-A binding to PDZ3, indicating that the scar has little effect.^<u>21,75</u>^ This makes sense in light of the PDZ binding pocket structural analyses we performed, which show no clear pocket to fit a terminal residue but rather a continuous groove, our demonstration that PDZ3 ligand binding has no electrostatic dependence, and evidence from other systems that show a terminal Val charged carboxylate did not improve PDZ3 binding.^<u>76</u>^ In light of the weak affinity of JAM-A to PSG, coupled with our finding that JAM-A binds well to PDZ1, we suggest that like claudins, PDZ1 binds JAM-A at a region upstream and disparate of its PDZ motif as PDZ1 affinities for JAM-A are markedly improved over PDZ3, which would not be true if they bound identically. However, PDZ1 and PDZ3 cannot bind simultaneously to JAM-A, as is true for claudins, indicating the binding sites overlap or the binding of one PDZ alters the binding of another. In the context of tight junction assembly, JAM-A binding to PDZ1 or ZO-1 appears only somewhat relevant, as it has been shown that PSG plays a greater role than PDZ1, and that ZO-2 is JAM-A’s primary MAGUK scaffold.^<u>21,77</u>^ Nonetheless, our data support the idea that PDZs are promiscuous for TJMP intracellular ligands and that individual PDZs can bind different regions depending on chemical or physical environments. JAM-A, and its intracellular domain have been shown to bind with variable affinities to PDZs upon changing mechanical forces.^<u>75,78</u>^ It is conceivable then that JAM-A interactions with specific ZO-1 and ZO-2 PDZ domains could be based on whether high affinity binding to internal sequences or low affinity binding to PDZ motifs are required for cellular mechanotransduction. Two binding modes may exist for functional flexibility.

Claudins were known to interact with PDZ1 but binding was thought to be low affinity, transient, and require a PDZ motif.^<u>19,20</u>^ We show that claudins interact with high affinity to PDZ1 and PDZ3 at their C-termini despite high divergence in sequence and charge but do not require their PDZ motifs to do so (**Fig. S17**). We validated this finding several ways, and data support the idea that TM4 proximal regions of claudins are the PDZ binding sites and that TM4 influences the selectivity and affinity of PDZ binding through PDZ-specific entropic/enthalpic contributions. For instance, although PDZ1 and PDZ3 overlap in claudin binding ability, they do not overlap in binding mechanism. Instead PDZ1 is ligand promiscuous and employs electrostatic interactions, whereas PDZ3 is ligand selective and binds in absence of charge. The diversity of PDZs and PDZ ligands in biology makes elucidating general principles a challenge. Briefly, we surmise that PDZs can bind either TM4 proximal (primary and high affinity) or PDZ motifs (secondary and low affinity) of claudin C-termini and that binding to either region can be altered by phosphorylation, electrostatic environment, or local structural changes, to make binding to one region preferential. Altering PDZ binding sites would alter the affinities and time scales of claudin/PDZ complexes and the organization and stability of larger ZO-1-bound complexes to have greater effects on tight junction assembly. Indeed, modifications of the C-termini through these means has been demonstrated to alter binding to PDZs and/or ZO-1 to effect tight junction assembly, structure, and epithelial permeability.^<u>79</u>^ The at times promiscuous, and at other, selective, modes of PDZ binding to claudins explains the often contradictory or construct-/cell-specific findings reported in the literature.^<u>19,62</u>^ Further studies are needed to demonstrate that the PDZ motif is not required for every claudin subtype, as our results focus on two classic claudins. We acknowledge that other subtypes may exclusively employ low affinity peptide-like interactions because ZO-1 interactions could be disfavored while interactions with other claudins are favorable.

Occludin and tricellulin form long-lived complexes with PSG through multivalent interactions where their CCD and Marvel domains interact with SG and unstructured IDRs with PDZ3. For this reason, the full-length TAMP constructs containing both a Marvel and C-terminal domain bound PSG best, explaining why peptides or domains alone have not yielded equivalent insights. Occludin uses residues 351-360 to bind PDZ3. Although occludin and tricellulin bind PSG with similar affinity, PDZ3 cannot bind tricellulin well because it lacks an equivalent binding site due to a 48 amino acid deletion. Instead, PDZ3 binds closer to tricellulin’s CCD. The sequence and length differences between TAMP C-terminal domains influences how and where interactions with individual PSG sub-domains occur, which suggests that the organization of PSG sub-domains differs when bound to occludin *versus* tricellulin. Altered organization is also demonstrated by our finding that occludin can bind PSG and PDZ1 simultaneously, while tricellulin has a limited ability to do so (**Fig. S8**). This organization could leave PDZ1 or PDZ3 free in tricellulin bound complexes with one PDZ. The literature presents conflicting reports regarding tricellulin/ZO-1 interactions, with some suggesting no interactions occur while others report interactions are required for biological function.^<u>23,55,80-82</u>^ Our data shows that tricellulin can interact with the PSG module of ZO-1 with biologically relevant affinities and time scales. At the same time, the tricellulin/ZO-1 complexes are not as long-lived as occludin/ZO-1 complexes, likely due to differences in PDZ3/C-terminal domain interactions that impart occludin with higher PSG binding affinity and slower complex dissociation rates *versus* tricellulin. These results explain several aspects of TAMP function at tight junctions. First, occludin regulates tricellulin cellular localization by “occluding” it from bicellular contacts, which can be explained by its higher affinity to ZO-1.^<u>28,55</u>^ Second, tricellulin resides at bicellular contacts in the absence of occludin and compensates for occludin function to maintain tight junction strand branching points as well as paracellular barriers— removal of either tricellulin or occludin, but not both, had no significant effect of tight junction ultrastructure or barrier function^<u>28</u>^. The conserved ability of TAMPs to bind PSG using both their Marvel and C-terminal domains explains functional overlaps and their abilities to compensate for one another in cells. Interestingly, this points to the clustering of ZO-1-bound TAMPs as being important contributors to the development of tight junction branching points. TAMPs mediate formation of tight junction strand branching points and the removal of both occludin and tricellulin is required to decrease the number of branching points and concomitantly, paracellular barrier integrity.^<u>28</u>^ Our data provides a mechanism for TAMP-enabled branching, where binding to PSG leaves PDZ1 free to bind claudin C-terminal domains and sequester claudins into larger ZO-1-bound complexes to reroute claudin-made strands. PDZ1 bound claudins may organize uniquely compared to unbound claudins to link converging strands and form new branches. In this way, one of the functions of TAMP/ZO-1 complexation could be to recruit claudins at the terminal ends of strands to these sites to reorient them to form and reinforce branches. However, given that occludin integrates directly to the most apical cell junctions, is found throughout strands, and dynamically diffuses in and out of tight junctions, we speculate that TAMPs enable ZO-1 function outside of solely branching points.^<u>38,39</u>^ Indeed, occludin dynamics at tight junctions correlates strongly with ZO-1’s that take it to and from membranes to intracellular pools.^<u>38</u>^ Coordinating ZO-1 trafficking to membranes, another product of TAMP/ZO-1 complexation, is explained by the inter-molecular interactions elucidated by this work. In these ways, ZO-1 and occludin are structurally linked to enable or disable key functionalities of the other during tight junction assembly and remodeling.

In the absence of other proteins, the ZO-1/TJMP interactome we uncover can be broadly summarized as occludin and tricellulin interact with high affinity and multivalency to the PSG module of ZO-1; claudins and JAM-A bind preferentially to PDZ1 over other ZO-1 sub-domains; while PDZ2 forms no comparable interactions with TJMPs (**Fig. 3**). This interactome would enable: i) localization of ZO-1 via PSG to membranes; ii) sequestering of claudin- or JAM-A-bound PDZ1 to TAMP/PSG complexes via PDZ1/PDZ3 complex formation; and iii) PDZ2 free for self-association to dimerize ZO-1—phenomena known to be critical for tight junction structure-function. Based on these findings, we propose a model of TJMP-enabled ZO-1 function in **Fig. 5**. However, our data also show that two TJMPs cannot bind a single PDZ at one time but that occludin, and to a lesser extent, tricellulin, can bind PDZ1 when bound to PDZ3 of PSG but claudins and JAM-A cannot (**Fig. S8**). This adds complexity and levels of regulation to the ZO-1/TJMP interactome that are not clearly translatable to our current understanding of tight junction form and function. In a system with such high overlap in interactions and functions, localization and abundance of TJMPs and ZO-1 at given times become primary determinants of binding. Further studies that pinpoint with high spatial and temporal resolutions the locations and amounts of these proteins are needed to elucidate a fuller understanding of the ZO-1/TJMP interactome when many are present.

**Fig. 5.**
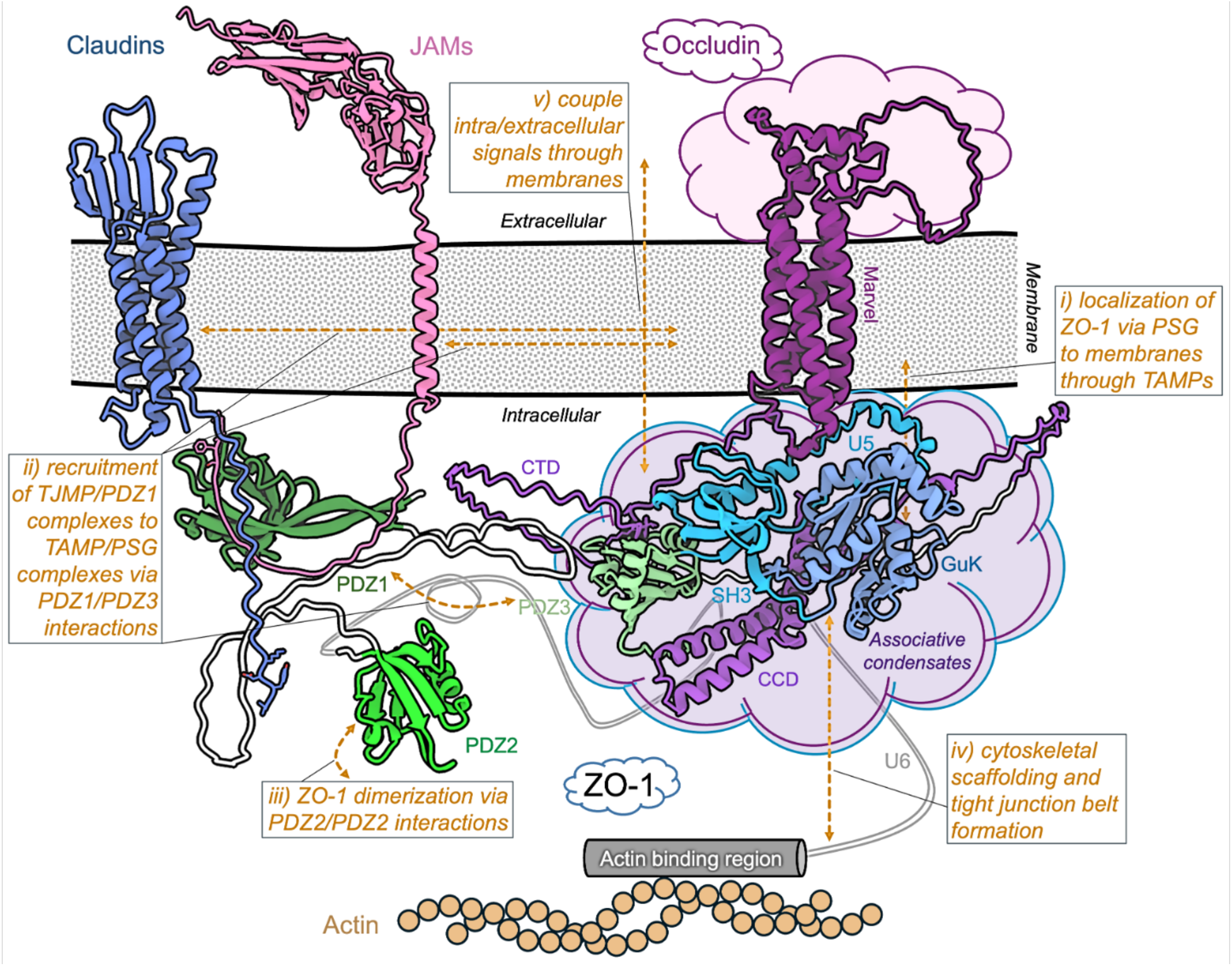
Model of TJMP-enabled ZO-1 function at tight junctions. Based on results from our biophysical and phase separation studies and those done previously, we propose a model where TAMPs (purple) are ZO-1 recruiting and membrane localizing TJMPs (i). This occurs through high affinity multivalent interactions to PSG (lt.green/lt.blue/blue) via TAMP’s Marvel and intracellular domains (CTD and CCD). TAMP/PSG complexes recruit PDZ1 (dk.green) bound to claudins (blue) or JAMs (pink) into larger assemblies via PDZ1/PD3 intra-molecular interactions (ii). PDZ2 (green) is not involved and instead homo-dimerizes ZO-1 (iii). TAMP/PSG complexation associates individual condensates (PSG, blue; occludin, purple; indigo clouds) and occludes the U6 motif from auto-inhibiting ZO-1 phase separation, allowing it and the nearby actin binding domain to bind actin to scaffold the associated complexes to the cytoskeleton and trigger tight junction belt formation (iv). The intracellular phase separation of occludin and PSG, and extracellular phase separation of TAMPs (purple cloud) and/or other TJMPs, couples intra/extracellular signaling, providing a mechanism for cell-cell communication via tight junctions (v). Binding and unbinding of ZO-1’s U5 to TAMPs or U6 to PSG can regulate this system, as can phosphorylation of these motifs and/or the U5/U6 homology regions of TAMPs, which would shift preferential binding from intra-molecular (tight junction disassembly) to inter-molecular (tight junction assembly) interactions or *vice versa*. Claudin and JAM sequestering to condensates can be regulated by phosphorylation of their intracellular domains, which alters binding to PDZ1. The role of the membrane or whether membrane microdomain formation (lipid rafts) plays additional regulatory roles should be a focus of future research.

Moreover, the interactome we uncover needs to be considered in the context of our discovery that occludin phase separates, and predictions that other TJMPs undergo phase separation within and outside of cells. Occludin but not tricellulin uses a 22 amino acid region of its C-terminal domain that is unique in sequence but not presence compared to tricellulin, to form liquid-like condensates in vitro. Occludin phase separates in chemical environments distinct from PSG, but both proteins cohabit the same condensates under conditions favorable to one, indicating that their phase separation is associative and driven by complimentary opposing charges. The global charges of TAMP intracellular domains is driven by the IDRs between the Marvel and CCD domains. Despite both occludin and tricellulin binding PSG well, only occludin has complementary electrostatics with PSG, indicating that occludin may play a larger role in regulating ZO-1 phase separation. Regulation of ZO-1/TAMP interactions and phase separation becomes a broader point of interest with our discovery of homology between ZO-1’s U5 and U6 motifs with TAMP intracellular domains. These motifs regulate interactions with TJMPs to alter tight junction organization or ZO-1 phase separation in vivo and in vitro via intra-molecular interactions with PSG.^<u>66</u>^ In this light, we speculate that attractive or repulsive charges between PSG and TAMPs define and regulate ZO-1/TAMP interactions and phase separation. This is validated by our and others results that show salt abrogates TAMP interactions with PSG as well as phase separation of occludin and PSG.^<u>66</u>^ Occludin binding to PSG would free U6 and ZO-1’s actin binding domain for cytoskeletal scaffolding that directs tight junction belt formation^<u>68,69</u>^, while U6 binding to PSG would favor auto-inhibition and ZO-1 uncoupling from both membranes (via loss of occludin binding) and cytoskeletal scaffolding (via actin binding domain).^<u>35,56,66</u>^ This system could be regulated further by post-translational modifications of residues within either ZO-1’s U5 and U6 motifs or occludin’s U5 and U6 homology regions. Indeed, the U5 and U6 homologous regions of occludin are rich in phosphorylation sites known to affect interactions with ZO-1^<u>66,83,84</u>^ and are now validated by our studies to interact with distinct domains of the PSG module; and phosphorylation of ZO-1 inhibits its phase separation ability.^<u>66</u>^ Taken together, this suggests that occludin interactions with PSG through unmodified residues would promote associative phase separation of both proteins, subsequent TJMP sequestering, and cytoskeletal scaffolding; while phosphorylation of key residues in these regions could either inhibit PSG binding to promote intra-molecular auto-inhibition of ZO-1 phase separation by U6 or alter occludin’s mode of PSG binding to disable the phase separation abilities of occludin and/or ZO-1—both being disabling to TJMP sequestering and tight junction belt formation. Further studies are needed to verify this and identify other inter-molecular regulatory mechanisms of phase separation.

The model of TJMP-enabled ZO-1 function provided in **Fig. 5** accounts for occludin and PSG phase separation that would, in addition to TJMP-enabled ZO-1 functions i-iii, further enable: iv) tight junction belt formation and cytoskeletal scaffolding through disabling U6 auto-inhibition; and v) coupling of intra/extracellular cell-cell communication via phase separation of associated occludin or other TJMP intra- or extra-cellular domains. In full, this work defines the interactions and mechanisms used by TJMPs to associate with ZO-1 and elucidates the regulatory consequences that they have on greater ZO-1 function. We identify occludin as a key interactor and enabler/regulator of ZO-1 functions known to direct tight junction assembly and dynamics, and TJMP phase separation as a means to transmit mechanochemical signals between adjacent cells via tight junctions.

## Methods

### Sequence alignments

Sequences of proteins were extracted from Uniprot^<u>85</u>^ and input manually, aligned using Clustal Omega^<u>86</u>^ then visualized using Jalview.^<u>87</u>^

### Protein expression and purification

All full length, truncated, or mutated membrane proteins were expressed and purified using methods previously described.^<u>41,43</u>^ Briefly, all genes were cloned into the vector pFastBac1 (ThermoFisher Scientific) and recombinant baculoviruses were generated and used to infect insect cells from *Trichoplusia ni* (Expression Systems). Cells were harvested using centrifugation, resuspended in Lysis buffer (50 mM Tris pH 7.4, 150 mM NaCl, 1 mM phenylmethylsulfonyl fluoride, and EDTA-free SigmaFast protease tablets (Sigma), then frozen and stored at −80°C until use. For purification, membranes were isolated by lysing cells using sonication followed by ultracentrifugation at 100,000 xg for 1 hour. Pellets were resuspended in Lysis buffer, dounce homogenized, then 1% n-dodecyl-β-D-maltopyranoside (DDM) and 0.04% cholesteryl hemisuccinate (CHS) (Anatrace) were added and allowed to rapidly mix overnight at 4°C. For JAM-A, 2% n-decyl-β-D-maltopyranoside (DM) and 0.04% CHS was used. Soluble material was separated using ultracentrifugation at 100,000 xg for 35 minutes and supernatants were mixed with HisPur affinity resin (ThermoFisher Scientific) overnight at 4°C with gentle mixing. Resin was collected in a bead capture column (Bio-Rad) and washed with 5 column volumes Wash A (50 mM Tris pH 7.4, 500 mM NaCl, 20 mM imidazole, and 10x critical micelle concentration (CMC) detergent) and 5 column volumes Wash B (50 mM Tris pH 7.4, 300 mM NaCl, 40 mM imidazole, and 10x CMC detergent) buffers. TJMPs were released from resins by treatment with bovine thrombin (MP Biomedical) in Wash T (50 mM Tris pH 8.0, 150 mM NaCl, and 5x CMC detergent) overnight at 4°C and collected in conicals. Purity and homogeneity were assessed by SDS-PAGE and size-exclusion chromatography (SEC), respectively. The latter required concentrating the proteins using 50 kDa MWCO concentrators (Millipore) and passing the solution through a 0.2 µm filter (Millipore) prior to injection on SEC using a Superdex 200 increase 10/300 column (Cytiva) equilibrated in SEC buffer (20 mM Hepes pH 7.4, 100 mM NaCl, 1% glycerol and 5x CMC detergent). Peaks corresponding to monomeric TJMPs were collected, pooled, concentrated, and flash frozen in liquid nitrogen and stored in −80°C then used in further studies.

All non-membrane proteins, which include ZO-1 subdomains PDZ2, PDZ3, SG, PSG, and the intracellular domains of TAMPs, were expressed and purified as follows. Constructs were cloned into the vector pET-14b (Novagen) and transformed into *Escherichia coli* BL21(DE3) (ThermoFisher Scientific) cells. Cells were grown in Terrific broth supplemented with ampicillin at 37°C and then induced with 1 mM isopropyl β-D-1-thiogalactopyranoside (IPTG, Inalco) once the OD_600_ reached 1.0 and then grown at 16°C for 18 hours. For PSG and SG, cells were induced with 0.1 mM IPTG once the OD_600_ reached 0.6 and then grown at 22°C for 18 hours. Cells were harvested by centrifugation and resuspended in Lysis buffer; then cells were lysed using sonication and ultracentrifuged at 100,000 xg for 1 hour to remove insoluble debris. Supernatants were mixed with HisPur resin and gently rocked at 4°C for 1 hour. Resin was washed in Wash A and B then proteins were eluted in Elution buffer (50 mM Tris pH 7.5, 150 mM NaCl, 10-500 mM imidazole, 10% glycerol). SDS-PAGE was used to check the fractions containing the purest protein and these fractions were pooled, concentrated using a 10 kDa MWCO concentrator (Millipore), filtered and purified further by SEC in SEC buffer. Buffers did not contain detergents for these constructs and SG and PSG buffers contained 500 mM NaCl to maintain protein stability. The SEC fractions containing monomers of these proteins were collected, pooled, concentrated, and flash frozen in liquid nitrogen and stored in −80°C then used in further studies.

For the non-membrane protein PDZ1, the insect cell methods for TJMPs were used for expression but purification methods used for other non-membrane proteins employed to produce and isolate this construct.

### X-ray Crystallography

Crystals of hsOCLN^393-522^ and hsOCLN^393-522-PO4^ were obtained using sitting-drop method at 18°C by mixing 1.5 µL of 3 mg/mL protein with 1.5 µL of 2.0 M Ammonium sulfate, 7.5% glycerol and 0.1 M sodium citrate pH 5.6. Crystals obtained were flash frozen and sent for data collection at APS beamline 23-ID-D. Data were integrated using iMosflm^<u>88</u>^ and scaled in SCALA.^<u>89</u>^ Molecular replacement was then performed using phenix.MR^<u>90</u>^ by searching for one copy of 1WPA.^<u>33</u>^ Both the structures were solved in space group P212121. For hsOCLN^393-522^ cell parameters were a=32.49 Å, b=35.83 Å, c=107.91 Å, α,β,γ=90°; while for hsOCLN^393-522-PO4^ cell parameters were a=32.89 Å, b=35.44 Å, c=107.47 Å, α,β,γ=90°. The structures after molecular replacement were refined using phenix.refine^<u>91,92</u>^ using multiple rounds of manual model building in coot.^<u>93,94</u>^ Final R_work_/R_free_ for were 26%/29% and 24%/27% for hsOCLN^393-522^ and hsOCLN^393-522-PO4^, respectively.

### Microscale Thermophoresis (MST)

Protein preparation for MST consisted of taking 20 µM SEC purified protein ligand, adding 2 molar excess of Cy5 dye (Cytiva), and incubating it at 4°C protected from light with gentle and occasional mixing for 2 hours. Free dye was removed using a PD-10 buffer exchange column (Bio-Rad) equilibrated in Biophysical buffer (20 mM Hepes pH 7.4, 150-500 mM NaCl, 1% glycerol and 0.04% DDM). Labelled ligand proteins were snap frozen in liquid nitrogen then stored at −80°C in small aliquots for downstream analyses.

Prior to complex formation for two-component binding studies, serial dilutions of analyte proteins were made from post-SEC concentrated stocks to prepare concentration series in low protein binding microtubes (Corning). Then 40 nM fluorescent ligand was added to each tube of the concentration series containing the analyte at 1:1 ratios by volume, mixed gently, then incubated at 25°C for 10 minutes. ∼5 µL of this mixture was then loaded into standard MST capillaries (NanoTemper) through capillary action and loaded into a Monolith X (NanoTemper) instrument for analysis at 25°C. Each capillary was scanned at 650 and 670 nm and binding response was evaluated by plotting the normalized fluorescence at 670 nM (Fnorm) against ligand concentration. Data fits were performed in K_D_-mode using the MO Control 2 software provided by the instrument then exported, normalized and plotted using Origin 9.0 to obtain K_D_. Data from two-component studies are reported in **Tables 1-3**.

For three-component binding studies, analytes at a single concentration (1-5 µM, analyte 1) were added to 40 nM fluorescent ligand, mixed, and incubated for 10 minutes at 25°C. Analytes that were prepared in serial dilutions as before (called analyte 2) were then added to these mixtures and incubated for another 10 minutes at 25°C. As before, ∼5 µL of mixtures were loaded into standard MST capillaries, scanned using a Monolith X. Binding curves to obtain K_D_ were acquired as in two-component studies. Data from three-component studies are reported in **Table 4**.

### Bio-Layer Interferometry (BLI)

Protein preparation for BLI consisted of taking 20 µM SEC purified protein ligand, adding 2 molar excess of EZ-link NHS-PEG4-Biotin (ThermoFisher Scientific), and incubating it at 4°C protected from light with gentle and occasional mixing for 2 hours. Free biotin was removed using a PD-10 buffer exchange column (Bio-Rad) equilibrated in Biophysical buffer (20 mM Hepes pH 7.4, 150-500 mM NaCl, 1% glycerol and 0.04% DDM). Labelled ligand proteins were snap frozen in liquid nitrogen then stored at −80°C in small aliquots for downstream analyses.

Biotinylated ligands were immobilized onto Octet streptavidin SA (Sartorious) biosensors after scouting experiments demonstrated that 100 nM of TAMPs and 75 nM of ZO-1 sub-domains were optimally loaded on the biosensor tip in 96-well flat bottom black plates (Greiner). Analytes at concentrations ranging between 0-5 µM were transferred to individual wells. No significant non-specific binding of analytes to naked SA sensors were observed at the highest concentrations tested. A typical experiment comprised 4 steps; Loading at 60 seconds, Baseline at 60 seconds; Association at 200 seconds; and Dissociation in the Baseline well at 200 seconds. All experiments were performed using an Octet R8 (Sartorius) instrument at 25°C using basic kinetic parameters and acquisition rates of 5 Hz averaged by 20 set within the embedded Octet Discovery 12.2 software.

Data were analyzed and fit using Octet Analysis Studio 12.2 software. Ligand-loaded sensors were subtracted from reference wells containing no analyte and for low affinity interactions, double referencing was used to subtract minor non-specific binding signals of the analyte to naked sensors at very high concentrations. The response units (nm) were aligned to the baseline for the last 5 seconds of measurement, and an inter-step correction was applied aligned to dissociation, then processed using Savitzky-Golay filtering. These processed data were fitted using a 1:1 Langmuir or 2:1 heterogeneous binding model with global full fits applied and R_max_ unlinked by sensor. Curves fit with 2:1 binding model included long constructs of occludin and tricellulin to PSG were justified because multivalent interactions were likely occurring based on other data. Data from BLI studies are reported in **Tables 2-3**.

### Turbidity assay

40 µM of SEC purified proteins were used straight or diluted in Phase Separation buffer (20 mM Hepes pH 7.4, 100-500 mM KCl, and 10% PEG 8,000) into low protein binding microtubes (Corning) and OD_600_ was measured after 10 minutes at 25°C using the nanophotometer NP80 (Implen). Control samples were tested in a buffer without PEG 8,000. Readings were taken at four concentrations that included 0, 5, 10, and 40 µM and done in triplicate then plotted using Origin 9.0 (OriginLab).

### Differential Interference Contrast (DIC) Microscopy

DIC microscopy was used to screen the phase separation abilities of occludin, tricellulin and PSG in other Phase Separation buffers with varied KCl (100, 250, and 500 mM) or PEG 8,000 (2, 5, and 10% w/v) concentrations. Protein concentrations ranging from 0.1-40 µM were tested across these buffers where proteins were diluted to their appropriate concentrations in Phase Separation buffer at 25°C. 6 µL of this mixture were placed into wells formed by adhesive SecureSeal Imaging spacers 9×0.12 mm (Grace Bio-Labs) then placed onto clean glass slides (VWR) and immediately covered with 61×0.17×1.5 mm 1 ounce coverglass (Grace Bio-Labs). Phase separation droplets were visualized using a Leica DM6B upright microscope (Leica Microsystems) and images were captured using 20x, 40x, and 100x objectives under oil. LAS X Lifescience Microscope software (Leica Microsystems) was used for image generation.

### Fluorescence Recovery after Photobleaching (FRAP) using Confocal Microscopy

Mixtures containing 99 µL of 10 µM unlabeled hsOCLN^271-522^ was mixed with 1 µL of 10 µM hsOCLN^271-522^ labeled with Cy5 as in MST studies in Phase Separation buffer to produce phase separation. The sample was transferred to the glass slide as for DIC microscopy and droplets were imaged using a Leica TCS SP8 confocal microscope system (Leica Microsystems), installed on a Leica DMi 8 inverted stand with 40x oil immersion objective and figures generated using the LAS X software. A circular region in the center of a droplet was bleached with a 404 nm laser at 100% and 4 pre-bleach, and 100 post-bleach frames were acquired with frame rate of 5.3 seconds corresponding to ∼10 minutes of recovery. Data were analyzed using ImageJ (Fuji). FRAP recovery was normalized to maximal pre-bleach and minimal post-bleach intensity. 15 droplets were photobleached and the data were plotted by averaging their intensities. The data were fit using a one-phase decay model and errors represent the standard deviation observed in intensity from data collection.

Client-partition assay using DIC microscopy. The hsOCLN^271-522^ and hsTRIC^368-558^ proteins were labeled with Alexa Fluor 488 (AF_488nm_) and PSG labeled with Alexa Fluor 647 (AF_647nm_, ThermoFisher Scientific) using the protocol as described above for MST. 20 µM of hsOCLN^271-522^ or hsTRIC^368-558^ containing 99% unlabeled and 1% labelled protein were mixed with 1 or 5 µM 100% labelled PSG in Phase Separation buffers. Phase separation was observed using a Leica DM6B upright microscope (Leica Microsystems) with 100x oil immersion objective and images were taken for DIC, AF_488nm_ (Ex_470±20nm_/Em_525±25nm_), and AF_647nm_ (Ex_620±30nm_/Em_700±38nm_) using the LAS X software. Controls with hsOCLN^271-522^, hsTRIC^368-558^, or PSG alone and without PEG 8,000 were performed.

## Acknowledgements

Research reported in this publication was supported by the National Institute of General Medical Sciences of the National Institutes of Health under Award Number: NIH NIGMS R35GM138368 (AJV). The content is solely the responsibility of the authors and does not necessarily represent the official views of the National Institutes of Health. We acknowledge the Pathology and Anatomical Sciences Research Service Center for providing DIC and Fluorescence Microscopy instrumentation and services used in the course of this study. Additionally, we would also like to acknowledge the use of the Advanced Photon Source GM/CA beamline. GM/CA@APS has been funded by the National Cancer Institute (ACB-12002) and the National Institute of General Medical Sciences (AGM-12006, P30GM138396). This research used resources of the Advanced Photon Source, a U.S. Department of Energy (DOE) Office of Science User Facility operated for the DOE Office of Science by Argonne National Laboratory under Contract No. DE-AC02-06CH11357. The Eiger 16M detector at GM/CA-XSD was funded by NIH grant S10OD012289.

## References

1. Woods, D.F., and Bryant, P.J. (1991). The discs-large tumor suppressor gene of Drosophila encodes a guanylate kinase homolog localized at septate junctions. Cell 66, 451–464. 10.1016/0092-8674(81)90009-x.

2. Anderson, J.M. (1996). Cell signalling: MAGUK magic. Curr Biol 6, 382–384. 10.1016/s0960-9822(02)00501-8.

3. Funke, L., Dakoji, S., and Bredt, D.S. (2005). Membrane-associated guanylate kinases regulate adhesion and plasticity at cell junctions. Annu Rev Biochem 74, 219–245. 10.1146/annurev.biochem.74.082803.133339.

4. Gonzalez-Mariscal, L., Betanzos, A., and Avila-Flores, A. (2000). MAGUK proteins: structure and role in the tight junction. Semin Cell Dev Biol 11, 315–324. 10.1006/scdb.2000.0178.

5. Hsueh, Y.P., Wang, T.F., Yang, F.C., and Sheng, M. (2000). Nuclear translocation and transcription regulation by the membrane-associated guanylate kinase CASK/LIN-2. Nature 404, 298–302. 10.1038/35005118.

6. Montgomery, J.M., Zamorano, P.L., and Garner, C.C. (2004). MAGUKs in synapse assembly and function: an emerging view. Cell Mol Life Sci 61, 911–929. 10.1007/s00018-003-3364-5.

7. Oliva, C., Escobedo, P., Astorga, C., Molina, C., and Sierralta, J. (2012). Role of the MAGUK protein family in synapse formation and function. Dev Neurobiol 72, 57–72. 10.1002/dneu.20949.

8. Gumbiner, B., Lowenkopf, T., and Apatira, D. (1991). Identification of a 160-kDa polypeptide that binds to the tight junction protein ZO-1. Proc Natl Acad Sci U S A 88, 3460–3464. 10.1073/pnas.88.8.3460.

9. Stevenson, B.R., Siliciano, J.D., Mooseker, M.S., and Goodenough, D.A. (1986). Identification of ZO-1: a high molecular weight polypeptide associated with the tight junction (zonula occludens) in a variety of epithelia. J Cell Biol 103, 755–766. 10.1083/jcb.103.3.755.

10. Hung, A.Y., and Sheng, M. (2002). PDZ domains: structural modules for protein complex assembly. J Biol Chem 277, 5699–5702. 10.1074/jbc.R100065200.

11. Lee, H.J., and Zheng, J.J. (2010). PDZ domains and their binding partners: structure, specificity, and modification. Cell Commun Signal 8, 8. 10.1186/1478-811X-8-8.

12. Basdevant, N., Weinstein, H., and Ceruso, M. (2006). Thermodynamic basis for promiscuity and selectivity in protein-protein interactions: PDZ domains, a case study. J Am Chem Soc 128, 12766–12777. 10.1021/ja060830y.

13. Gerek, Z.N., Keskin, O., and Ozkan, S.B. (2009). Identification of specificity and promiscuity of PDZ domain interactions through their dynamic behavior. Proteins 77, 796–811. 10.1002/prot.22492.

14. te Velthuis, A.J., Sakalis, P.A., Fowler, D.A., and Bagowski, C.P. (2011). Genome-wide analysis of PDZ domain binding reveals inherent functional overlap within the PDZ interaction network. PLoS One 6, e16047. 10.1371/journal.pone.0016047.

15. Ivarsson, Y. (2012). Plasticity of PDZ domains in ligand recognition and signaling. FEBS Lett 586, 2638–2647. 10.1016/j.febslet.2012.04.015.

16. Fanning, A.S., Jameson, B.J., Jesaitis, L.A., and Anderson, J.M. (1998). The tight junction protein ZO-1 establishes a link between the transmembrane protein occludin and the actin cytoskeleton. J Biol Chem 273, 29745–29753. 10.1074/jbc.273.45.29745.

17. Schlingmann, B., Overgaard, C.E., Molina, S.A., Lynn, K.S., Mitchell, L.A., Dorsainvil White, S., Mattheyses, A.L., Guidot, D.M., Capaldo, C.T., and Koval, M. (2016). Regulation of claudin/zonula occludens-1 complexes by hetero-claudin interactions. Nat Commun 7, 12276. 10.1038/ncomms12276.

18. Umeda, K., Ikenouchi, J., Katahira-Tayama, S., Furuse, K., Sasaki, H., Nakayama, M., Matsui, T., Tsukita, S., Furuse, M., and Tsukita, S. (2006). ZO-1 and ZO-2 independently determine where claudins are polymerized in tight-junction strand formation. Cell 126, 741–754. 10.1016/j.cell.2006.06.043.

19. Itoh, M., Furuse, M., Morita, K., Kubota, K., Saitou, M., and Tsukita, S. (1999). Direct binding of three tight junction-associated MAGUKs, ZO-1, ZO-2, and ZO-3, with the COOH termini of claudins. J Cell Biol 147, 1351–1363. 10.1083/jcb.147.6.1351.

20. Nomme, J., Antanasijevic, A., Caffrey, M., Van Itallie, C.M., Anderson, J.M., Fanning, A.S., and Lavie, A. (2015). Structural Basis of a Key Factor Regulating the Affinity between the Zonula Occludens First PDZ Domain and Claudins. J Biol Chem 290, 16595–16606. 10.1074/jbc.M115.646695.

21. Nomme, J., Fanning, A.S., Caffrey, M., Lye, M.F., Anderson, J.M., and Lavie, A. (2011). The Src homology 3 domain is required for junctional adhesion molecule binding to the third PDZ domain of the scaffolding protein ZO-1. J Biol Chem 286, 43352–43360. 10.1074/jbc.M111.304089.

22. Bewley, M.C., Tash, B.R., Tian, F., and Flanagan, J.M. (2013). A complex affair: Attraction and repulsion make occludin and ZO-1 function! Tissue Barriers 1, e23496. 10.4161/tisb.23496.

23. Tash, B.R., Bewley, M.C., Russo, M., Keil, J.M., Griffin, K.A., Sundstrom, J.M., Antonetti, D.A., Tian, F., and Flanagan, J.M. (2012). The occludin and ZO-1 complex, defined by small angle X-ray scattering and NMR, has implications for modulating tight junction permeability. Proc Natl Acad Sci U S A 109, 10855–10860. 10.1073/pnas.1121390109.

24. Van Itallie, C.M., and Anderson, J.M. (2013). Claudin interactions in and out of the tight junction. Tissue Barriers 1, e25247. 10.4161/tisb.25247.

25. Sasaki, H., Matsui, C., Furuse, K., Mimori-Kiyosue, Y., Furuse, M., and Tsukita, S. (2003). Dynamic behavior of paired claudin strands within apposing plasma membranes. Proc Natl Acad Sci U S A 100, 3971–3976. 10.1073/pnas.0630649100.

26. Cording, J., Berg, J., Kading, N., Bellmann, C., Tscheik, C., Westphal, J.K., Milatz, S., Gunzel, D., Wolburg, H., Piontek, J., et al. (2013). In tight junctions, claudins regulate the interactions between occludin, tricellulin and marvelD3, which, inversely, modulate claudin oligomerization. J Cell Sci 126, 554–564. 10.1242/jcs.114306.

27. Furuse, M., Sasaki, H., and Tsukita, S. (1999). Manner of interaction of heterogeneous claudin species within and between tight junction strands. J Cell Biol 147, 891–903. 10.1083/jcb.147.4.891.

28. Saito, A.C., Higashi, T., Fukazawa, Y., Otani, T., Tauchi, M., Higashi, A.Y., Furuse, M., and Chiba, H. (2021). Occludin and tricellulin facilitate formation of anastomosing tight-junction strand network to improve barrier function. Mol Biol Cell 32, 722–738. 10.1091/mbc.E20-07-0464.

29. Nusrat, A., Brown, G.T., Tom, J., Drake, A., Bui, T.T., Quan, C., and Mrsny, R.J. (2005). Multiple protein interactions involving proposed extracellular loop domains of the tight junction protein occludin. Mol Biol Cell 16, 1725–1734. 10.1091/mbc.e04-06-0465.

30. Monteiro, A.C., Luissint, A.C., Sumagin, R., Lai, C., Vielmuth, F., Wolf, M.F., Laur, O., Reiss, K., Spindler, V., Stehle, T., et al. (2014). Trans-dimerization of JAM-A regulates Rap2 and is mediated by a domain that is distinct from the cis-dimerization interface. Mol Biol Cell 25, 1574–1585. 10.1091/mbc.E14-01-0018.

31. Peng, B.H., White, M.A., Campbell, G.A., Robert, J.J., Lee, J.C., and Sutton, R.B. (2005). Crystallization and preliminary X-ray diffraction of the ZO-binding domain of human occludin. Acta Crystallogr Sect F Struct Biol Cryst Commun 61, 369–371. 10.1107/S1744309105007475.

32. Schuetz, A., Radusheva, V., Krug, S.M., and Heinemann, U. (2017). Crystal structure of the tricellulin C-terminal coiled-coil domain reveals a unique mode of dimerization. Ann N Y Acad Sci 1405, 147–159. 10.1111/nyas.13408.

33. Li, Y., Fanning, A.S., Anderson, J.M., and Lavie, A. (2005). Structure of the conserved cytoplasmic C-terminal domain of occludin: identification of the ZO-1 binding surface. J Mol Biol 352, 151–164. 10.1016/j.jmb.2005.07.017.

34. Van Itallie, C.M., Fanning, A.S., Bridges, A., and Anderson, J.M. (2009). ZO-1 stabilizes the tight junction solute barrier through coupling to the perijunctional cytoskeleton. Mol Biol Cell 20, 3930–3940. 10.1091/mbc.e09-04-0320.

35. Haas, A.J., Zihni, C., Krug, S.M., Maraspini, R., Otani, T., Furuse, M., Honigmann, A., Balda, M.S., and Matter, K. (2022). ZO-1 Guides Tight Junction Assembly and Epithelial Morphogenesis via Cytoskeletal Tension-Dependent and -Independent Functions. Cells 11. 10.3390/cells11233775.

36. Kuo, W.T., Odenwald, M.A., Turner, J.R., and Zuo, L. (2022). Tight junction proteins occludin and ZO-1 as regulators of epithelial proliferation and survival. Ann N Y Acad Sci 1514, 21–33. 10.1111/nyas.14798.

37. Van Itallie, C.M., and Anderson, J.M. (2018). Phosphorylation of tight junction transmembrane proteins: Many sites, much to do. Tissue Barriers 6, e1382671. 10.1080/21688370.2017.1382671.

38. Shen, L., Weber, C.R., and Turner, J.R. (2008). The tight junction protein complex undergoes rapid and continuous molecular remodeling at steady state. J Cell Biol 181, 683–695. 10.1083/jcb.200711165.

39. Van Itallie, C.M., Lidman, K.F., Tietgens, A.J., and Anderson, J.M. (2019). Newly synthesized claudins but not occludin are added to the basal side of the tight junction. Mol Biol Cell 30, 1406–1424. 10.1091/mbc.E19-01-0008.

40. Erramilli, S.K., Dominik, P.K., Ogbu, C.P., Kossiakoff, A.A., and Vecchio, A.J. (2024). Structural and biophysical insights into targeting of claudin-4 by a synthetic antibody fragment. Commun Biol 7, 733. 10.1038/s42003-024-06437-6.

41. Ogbu, C.P., de Las Alas, M., Mandriota, A.M., Liu, X., Kapoor, S., Choudhury, J., Ruma, Y.N., Goodman, M.C., Sanders, C.R., Gonen, T., et al. (2025). Biophysical basis of tight junction barrier modulation by a pan-claudin-binding molecule. PNAS Nexus 4, pgaf189. 10.1093/pnasnexus/pgaf189.

42. Rathnayake, S.S., Erramilli, S.K., Kossiakoff, A.A., and Vecchio, A.J. (2024). Cryo-EM structures of Clostridium perfringens enterotoxin bound to its human receptor, claudin-4. Structure 32, 1936–1951 e1935. 10.1016/j.str.2024.09.015.

43. Vecchio, A.J., Rathnayake, S.S., and Stroud, R.M. (2021). Structural basis for Clostridium perfringens enterotoxin targeting of claudins at tight junctions in mammalian gut. Proc Natl Acad Sci U S A 118. 10.1073/pnas.2024651118.

44. Vecchio, A.J., and Stroud, R.M. (2019). Claudin-9 structures reveal mechanism for toxin-induced gut barrier breakdown. Proc Natl Acad Sci U S A 116, 17817–17824. 10.1073/pnas.1908929116.

45. Appleton, B.A., Zhang, Y., Wu, P., Yin, J.P., Hunziker, W., Skelton, N.J., Sidhu, S.S., and Wiesmann, C. (2006). Comparative structural analysis of the Erbin PDZ domain and the first PDZ domain of ZO-1. Insights into determinants of PDZ domain specificity. J Biol Chem 281, 22312–22320. 10.1074/jbc.M602901200.

46. Fanning, A.S., Lye, M.F., Anderson, J.M., and Lavie, A. (2007). Domain swapping within PDZ2 is responsible for dimerization of ZO proteins. J Biol Chem 282, 37710–37716. 10.1074/jbc.M707255200.

47. Pan, L., Chen, J., Yu, J., Yu, H., and Zhang, M. (2011). The structure of the PDZ3-SH3-GuK tandem of ZO-1 protein suggests a supramodular organization of the membrane-associated guanylate kinase (MAGUK) family scaffold protein core. J Biol Chem 286, 40069–40074. 10.1074/jbc.C111.293084.

48. Cordenonsi, M., Turco, F., D’Atri, F., Hammar, E., Martinucci, G., Meggio, F., and Citi, S. (1999). Xenopus laevis occludin. Identification of in vitro phosphorylation sites by protein kinase CK2 and association with cingulin. Eur J Biochem 264, 374–384. 10.1046/j.1432-1327.1999.00616.x.

49. Chen, Y.H., Lu, Q., Goodenough, D.A., and Jeansonne, B. (2002). Nonreceptor tyrosine kinase c-Yes interacts with occludin during tight junction formation in canine kidney epithelial cells. Mol Biol Cell 13, 1227–1237. 10.1091/mbc.01-08-0423.

50. McKenzie, J.A., Riento, K., and Ridley, A.J. (2006). Casein kinase I epsilon associates with and phosphorylates the tight junction protein occludin. FEBS Lett 580, 2388–2394. 10.1016/j.febslet.2006.03.048.

51. Andreeva, A.Y., Krause, E., Muller, E.C., Blasig, I.E., and Utepbergenov, D.I. (2001). Protein kinase C regulates the phosphorylation and cellular localization of occludin. J Biol Chem 276, 38480–38486. 10.1074/jbc.M104923200.

52. Yamamoto, M., Ramirez, S.H., Sato, S., Kiyota, T., Cerny, R.L., Kaibuchi, K., Persidsky, Y., and Ikezu, T. (2008). Phosphorylation of claudin-5 and occludin by rho kinase in brain endothelial cells. Am J Pathol 172, 521–533. 10.2353/ajpath.2008.070076.

53. Du, D., Xu, F., Yu, L., Zhang, C., Lu, X., Yuan, H., Huang, Q., Zhang, F., Bao, H., Jia, L., et al. (2010). The tight junction protein, occludin, regulates the directional migration of epithelial cells. Dev Cell 18, 52–63. 10.1016/j.devcel.2009.12.008.

54. Nunbhakdi-Craig, V., Machleidt, T., Ogris, E., Bellotto, D., White, C.L., 3rd, and Sontag, E. (2002). Protein phosphatase 2A associates with and regulates atypical PKC and the epithelial tight junction complex. J Cell Biol 158, 967–978. 10.1083/jcb.200206114.

55. Raleigh, D.R., Marchiando, A.M., Zhang, Y., Shen, L., Sasaki, H., Wang, Y., Long, M., and Turner, J.R. (2010). Tight junction-associated MARVEL proteins marveld3, tricellulin, and occludin have distinct but overlapping functions. Mol Biol Cell 21, 1200–1213. 10.1091/mbc.e09-08-0734.

56. Fanning, A.S., Little, B.P., Rahner, C., Utepbergenov, D., Walther, Z., and Anderson, J.M. (2007). The unique-5 and −6 motifs of ZO-1 regulate tight junction strand localization and scaffolding properties. Mol Biol Cell 18, 721–731. 10.1091/mbc.e06-08-0764.

57. Muller, S.L., Portwich, M., Schmidt, A., Utepbergenov, D.I., Huber, O., Blasig, I.E., and Krause, G. (2005). The tight junction protein occludin and the adherens junction protein alpha-catenin share a common interaction mechanism with ZO-1. J Biol Chem 280, 3747–3756. 10.1074/jbc.M411365200.

58. Schmidt, A., Utepbergenov, D.I., Krause, G., and Blasig, I.E. (2001). Use of surface plasmon resonance for real-time analysis of the interaction of ZO-1 and occludin. Biochem Biophys Res Commun 288, 1194–1199. 10.1006/bbrc.2001.5914.

59. Schmidt, A., Utepbergenov, D.I., Mueller, S.L., Beyermann, M., Schneider-Mergener, J., Krause, G., and Blasig, I.E. (2004). Occludin binds to the SH3-hinge-GuK unit of zonula occludens protein 1: potential mechanism of tight junction regulation. Cell Mol Life Sci 61, 1354–1365. 10.1007/s00018-004-4010-6.

60. Rodgers, L.S., Beam, M.T., Anderson, J.M., and Fanning, A.S. (2013). Epithelial barrier assembly requires coordinated activity of multiple domains of the tight junction protein ZO-1. J Cell Sci 126, 1565–1575. 10.1242/jcs.113399.

61. Ruffer, C., and Gerke, V. (2004). The C-terminal cytoplasmic tail of claudins 1 and 5 but not its PDZ-binding motif is required for apical localization at epithelial and endothelial tight junctions. Eur J Cell Biol 83, 135–144. 10.1078/0171-9335-00366.

62. Fujiwara, S., Nguyen, T.P., Furuse, K., Fukazawa, Y., Otani, T., and Furuse, M. (2022). Tight junction formation by a claudin mutant lacking the COOH-terminal PDZ domain-binding motif. Ann N Y Acad Sci 1516, 85–94. 10.1111/nyas.14881.

63. Goverde, C.A., Pacesa, M., Goldbach, N., Dornfeld, L.J., Balbi, P.E.M., Georgeon, S., Rosset, S., Kapoor, S., Choudhury, J., Dauparas, J., et al. (2024). Computational design of soluble and functional membrane protein analogues. Nature 631, 449–458. 10.1038/s41586-024-07601-y.

64. Otani, T., Nguyen, T.P., Tokuda, S., Sugihara, K., Sugawara, T., Furuse, K., Miura, T., Ebnet, K., and Furuse, M. (2019). Claudins and JAM-A coordinately regulate tight junction formation and epithelial polarity. J Cell Biol 218, 3372–3396. 10.1083/jcb.201812157.

65. Severson, E.A., Jiang, L., Ivanov, A.I., Mandell, K.J., Nusrat, A., and Parkos, C.A. (2008). Cis-dimerization mediates function of junctional adhesion molecule A. Mol Biol Cell 19, 1862–1872. 10.1091/mbc.e07-09-0869.

66. Beutel, O., Maraspini, R., Pombo-Garcia, K., Martin-Lemaitre, C., and Honigmann, A. (2019). Phase Separation of Zonula Occludens Proteins Drives Formation of Tight Junctions. Cell 179, 923–936 e911. 10.1016/j.cell.2019.10.011.

67. Kinoshita, N., Yamamoto, T.S., Yasue, N., Takagi, C., Fujimori, T., and Ueno, N. (2022). Force-dependent remodeling of cytoplasmic ZO-1 condensates contributes to cell-cell adhesion through enhancing tight junctions. iScience 25, 103846. 10.1016/j.isci.2022.103846.

68. Pombo-Garcia, K., Adame-Arana, O., Martin-Lemaitre, C., Julicher, F., and Honigmann, A. (2024). Membrane prewetting by condensates promotes tight-junction belt formation. Nature 632, 647–655. 10.1038/s41586-024-07726-0.

69. Sun, D., Zhao, X., Wiegand, T., Martin-Lemaitre, C., Borianne, T., Kleinschmidt, L., Grill, S.W., Hyman, A.A., Weber, C., and Honigmann, A. (2025). Assembly of tight junction belts by ZO1 surface condensation and local actin polymerization. Dev Cell 60, 1234–1250 e1236. 10.1016/j.devcel.2024.12.012.

70. Wilson, C., Lewis, K.A., Fitzkee, N.C., Hough, L.E., and Whitten, S.T. (2023). ParSe 2.0: A web tool to identify drivers of protein phase separation at the proteome level. Protein Sci 32, e4756. 10.1002/pro.4756.

71. Paiz, E.A., Allen, J.H., Correia, J.J., Fitzkee, N.C., Hough, L.E., and Whitten, S.T. (2021). Beta turn propensity and a model polymer scaling exponent identify intrinsically disordered phase-separating proteins. J Biol Chem 297, 101343. 10.1016/j.jbc.2021.101343.

72. Huang, Y., Bai, Y., Jin, W., Shen, D., Lyu, H., Zeng, L., Wang, M., and Liu, Y. (2021). Common Pitfalls and Recommendations for Using a Turbidity Assay to Study Protein Phase Separation. Biochemistry 60, 2447–2456. 10.1021/acs.biochem.1c00386.

73. Van Lindt, J., Bratek-Skicki, A., Nguyen, P.N., Pakravan, D., Duran-Armenta, L.F., Tantos, A., Pancsa, R., Van Den Bosch, L., Maes, D., and Tompa, P. (2021). A generic approach to study the kinetics of liquid-liquid phase separation under near-native conditions. Commun Biol 4, 77. 10.1038/s42003-020-01596-8.

74. Steed, E., Rodrigues, N.T., Balda, M.S., and Matter, K. (2009). Identification of MarvelD3 as a tight junction-associated transmembrane protein of the occludin family. BMC Cell Biol 10, 95. 10.1186/1471-2121-10-95.

75. Vachharajani, V.T., DeJong, M.P., Dutta, S., Chapman, J., Ghosh, E., Singharoy, A., and Dunn, A.R. (2025). PDZ Domains from the Junctional Proteins Afadin and ZO-1 Act as Mechanosensors. bioRxiv. 10.1101/2023.09.24.559210.

76. Harris, B.Z., Lau, F.W., Fujii, N., Guy, R.K., and Lim, W.A. (2003). Role of electrostatic interactions in PDZ domain ligand recognition. Biochemistry 42, 2797–2805. 10.1021/bi027061p.

77. Monteiro, A.C., Sumagin, R., Rankin, C.R., Leoni, G., Mina, M.J., Reiter, D.M., Stehle, T., Dermody, T.S., Schaefer, S.A., Hall, R.A., et al. (2013). JAM-A associates with ZO-2, afadin, and PDZ-GEF1 to activate Rap2c and regulate epithelial barrier function. Mol Biol Cell 24, 2849–2860. 10.1091/mbc.E13-06-0298.

78. Scott, D.W., Tolbert, C.E., and Burridge, K. (2016). Tension on JAM-A activates RhoA via GEF-H1 and p115 RhoGEF. Mol Biol Cell 27, 1420–1430. 10.1091/mbc.E15-12-0833.

79. Suzuki, T. (2013). Regulation of intestinal epithelial permeability by tight junctions. Cell Mol Life Sci 70, 631–659. 10.1007/s00018-012-1070-x.

80. Riazuddin, S., Ahmed, Z.M., Fanning, A.S., Lagziel, A., Kitajiri, S., Ramzan, K., Khan, S.N., Chattaraj, P., Friedman, P.L., Anderson, J.M., et al. (2006). Tricellulin is a tight-junction protein necessary for hearing. Am J Hum Genet 79, 1040–1051. 10.1086/510022.

81. Ikenouchi, J., Furuse, M., Furuse, K., Sasaki, H., Tsukita, S., and Tsukita, S. (2005). Tricellulin constitutes a novel barrier at tricellular contacts of epithelial cells. J Cell Biol 171, 939–945. 10.1083/jcb.200510043.

82. Cho, Y., Haraguchi, D., Shigetomi, K., Matsuzawa, K., Uchida, S., and Ikenouchi, J. (2022). Tricellulin secures the epithelial barrier at tricellular junctions by interacting with actomyosin. J Cell Biol 221. 10.1083/jcb.202009037.

83. Staddon, J.M., Herrenknecht, K., Smales, C., and Rubin, L.L. (1995). Evidence that tyrosine phosphorylation may increase tight junction permeability. J Cell Sci 108 (Pt 2), 609–619. 10.1242/jcs.108.2.609.

84. Balda, M.S., Anderson, J.M., and Matter, K. (1996). The SH3 domain of the tight junction protein ZO-1 binds to a serine protein kinase that phosphorylates a region C-terminal to this domain. FEBS Lett 399, 326–332. 10.1016/s0014-5793(96)01352-x.

85. UniProt, C. (2025). UniProt: the Universal Protein Knowledgebase in 2025. Nucleic Acids Res 53, D609–D617. 10.1093/nar/gkae1010.

86. Madeira, F., Madhusoodanan, N., Lee, J., Eusebi, A., Niewielska, A., Tivey, A.R.N., Lopez, R., and Butcher, S. (2024). The EMBL-EBI Job Dispatcher sequence analysis tools framework in 2024. Nucleic Acids Res 52, W521–W525. 10.1093/nar/gkae241.

87. Waterhouse, A.M., Procter, J.B., Martin, D.M., Clamp, M., and Barton, G.J. (2009). Jalview Version 2--a multiple sequence alignment editor and analysis workbench. Bioinformatics 25, 1189–1191. 10.1093/bioinformatics/btp033.

88. Battye, T.G., Kontogiannis, L., Johnson, O., Powell, H.R., and Leslie, A.G. (2011). iMOSFLM: a new graphical interface for diffraction-image processing with MOSFLM. Acta Crystallogr D Biol Crystallogr 67, 271–281. 10.1107/S0907444910048675.

89. Evans, P.R. (2011). An introduction to data reduction: space-group determination, scaling and intensity statistics. Acta Crystallogr D Biol Crystallogr 67, 282–292. 10.1107/S090744491003982X.

90. Adams, P.D., Afonine, P.V., Bunkoczi, G., Chen, V.B., Davis, I.W., Echols, N., Headd, J.J., Hung, L.W., Kapral, G.J., Grosse-Kunstleve, R.W., et al. (2010). PHENIX: a comprehensive Python-based system for macromolecular structure solution. Acta Crystallogr D Biol Crystallogr 66, 213–221. 10.1107/S0907444909052925.

91. Afonine, P.V., Grosse-Kunstleve, R.W., Echols, N., Headd, J.J., Moriarty, N.W., Mustyakimov, M., Terwilliger, T.C., Urzhumtsev, A., Zwart, P.H., and Adams, P.D. (2012). Towards automated crystallographic structure refinement with phenix.refine. Acta Crystallogr D Biol Crystallogr 68, 352–367. 10.1107/S0907444912001308.

92. Headd, J.J., Echols, N., Afonine, P.V., Grosse-Kunstleve, R.W., Chen, V.B., Moriarty, N.W., Richardson, D.C., Richardson, J.S., and Adams, P.D. (2012). Use of knowledge-based restraints in phenix.refine to improve macromolecular refinement at low resolution. Acta Crystallogr D Biol Crystallogr 68, 381–390. 10.1107/S0907444911047834.

93. Emsley, P., and Cowtan, K. (2004). Coot: model-building tools for molecular graphics. Acta Crystallogr D Biol Crystallogr 60, 2126–2132. 10.1107/S0907444904019158.

94. Emsley, P., Lohkamp, B., Scott, W.G., and Cowtan, K. (2010). Features and development of Coot. Acta Crystallogr D Biol Crystallogr 66, 486–501. 10.1107/S0907444910007493.

95. Jumper, J., Evans, R., Pritzel, A., Green, T., Figurnov, M., Ronneberger, O., Tunyasuvunakool, K., Bates, R., Zidek, A., Potapenko, A., et al. (2021). Highly accurate protein structure prediction with AlphaFold. Nature 596, 583–589. 10.1038/s41586-021-03819-2.

