## Supplementary material for "The biophysical interaction network that couples zona occludens-1 to membrane proteins at tight junctions": SI

**Figures S1-17**
**Table S1**

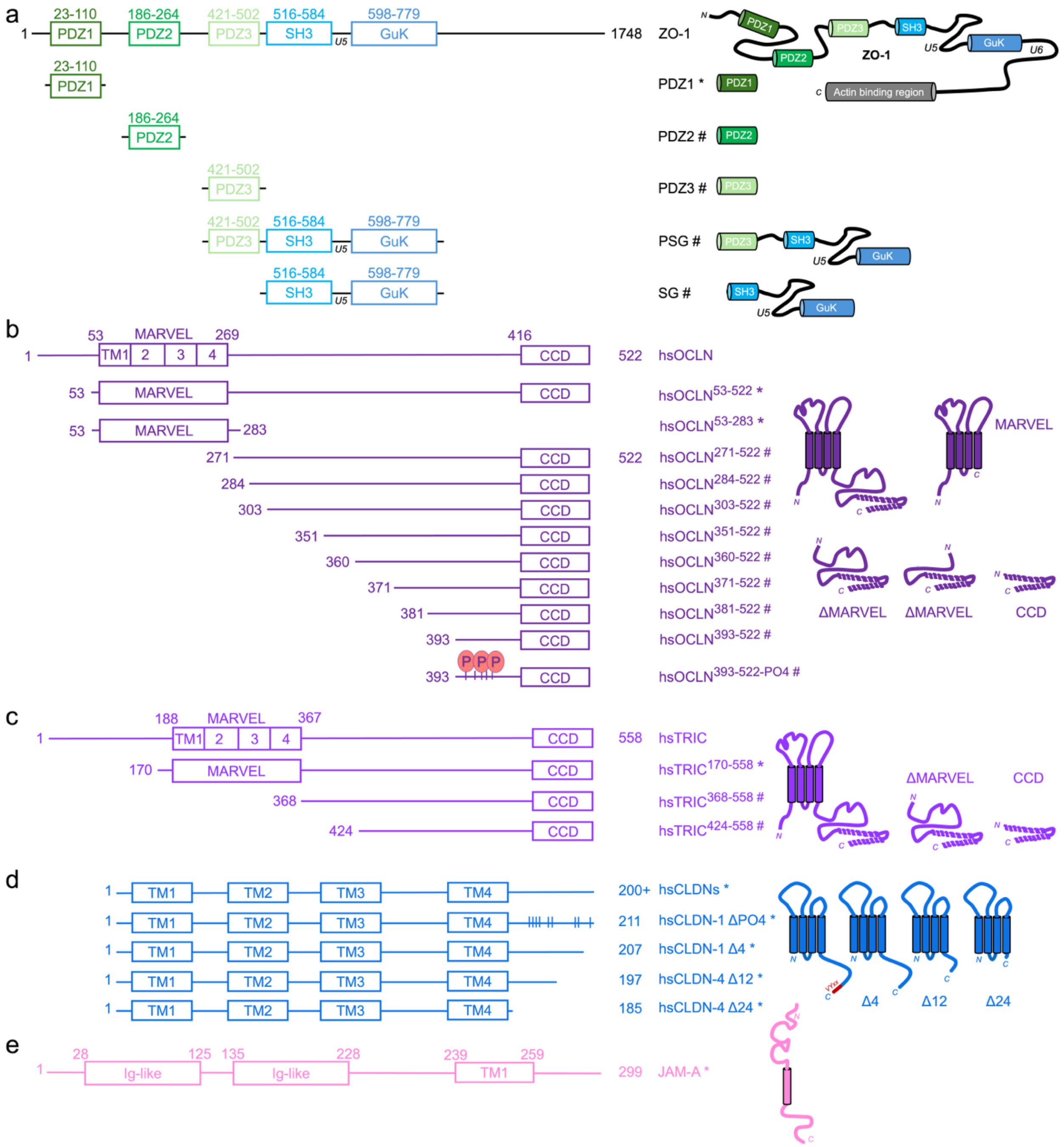


**Fig. S1. Overview of major constructs.** Schematic and topological overviews of ZO-1 and TJMPs used in this study include (a) ZO-1. (b) Occludin. (c) Tricellulin. (d) Claudins. (e) JAM-A. Proteins are colored as in **Fig. 1**. # represent constructs expressed in bacteria while * demark those expressed in insect systems. Unlabeled proteins represent full length wild type protein.

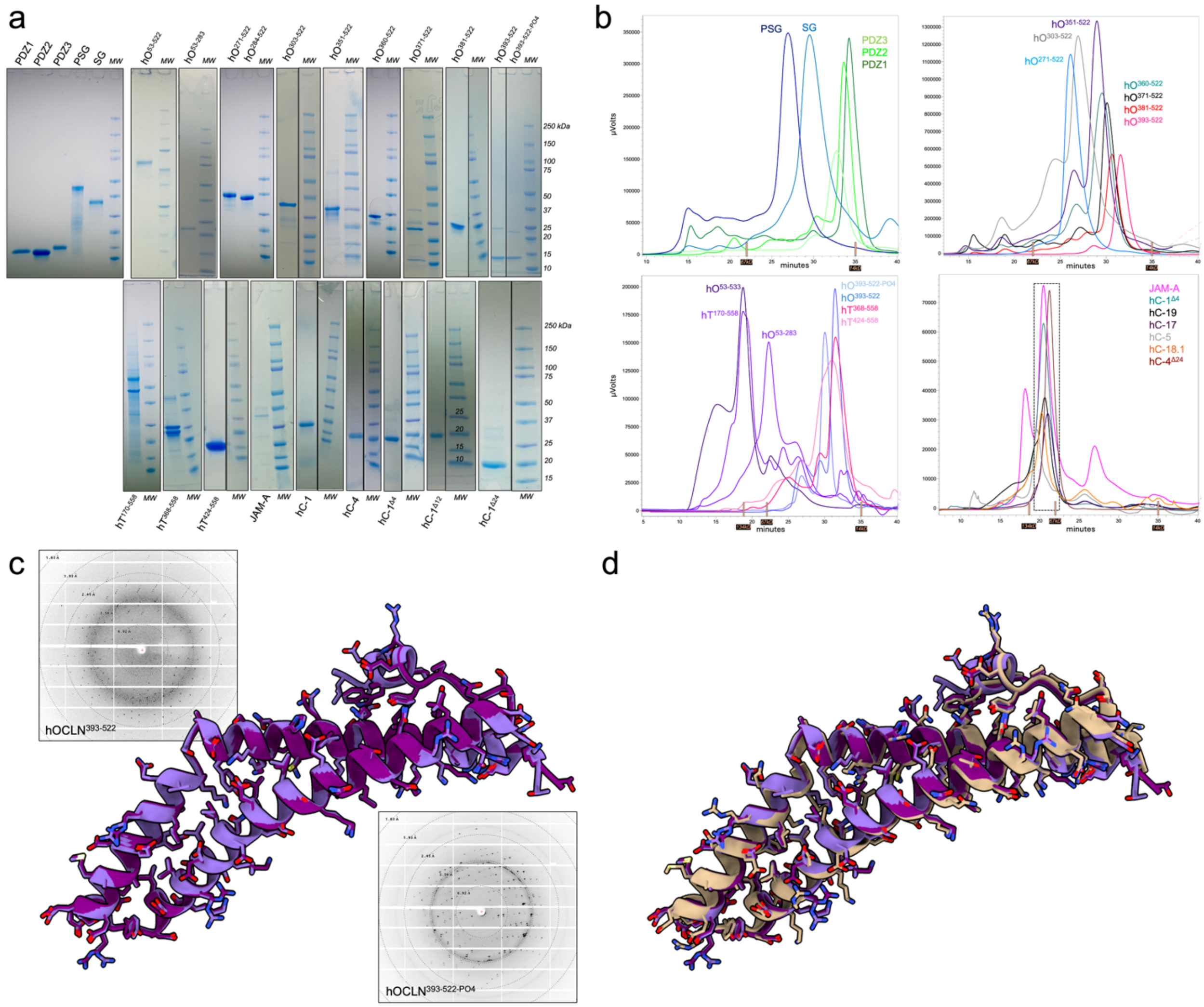

**Fig. S2. Biochemical and biophysical preparation of ZO-1 and TJMP constructs.** (a) SDS-PAGE analysis of protein purity post-affinity and size-exclusion chromatography (SEC). Each construct is labeled next to molecular weight standards. (b) SEC profiles of post-affinity purified proteins. Molecular weight standards are labeled on the x axis. (c) X-ray crystal structures and associated diffraction patterns of hsOCLN$^{393\text{-}533}$ (dk purple) and hsOCLN$^{393\text{-}533\text{-}PO4}$ (lt purple). (d) Structural overlap of hsOCLN$^{393\text{-}533}$ (dk purple) and hsOCLN$^{393\text{-}533\text{-}PO4}$ (lt purple) onto the crystal structure of occludin coiled-coil domain (PDB ID 1xaw, tan) determined by Li et al.[1]

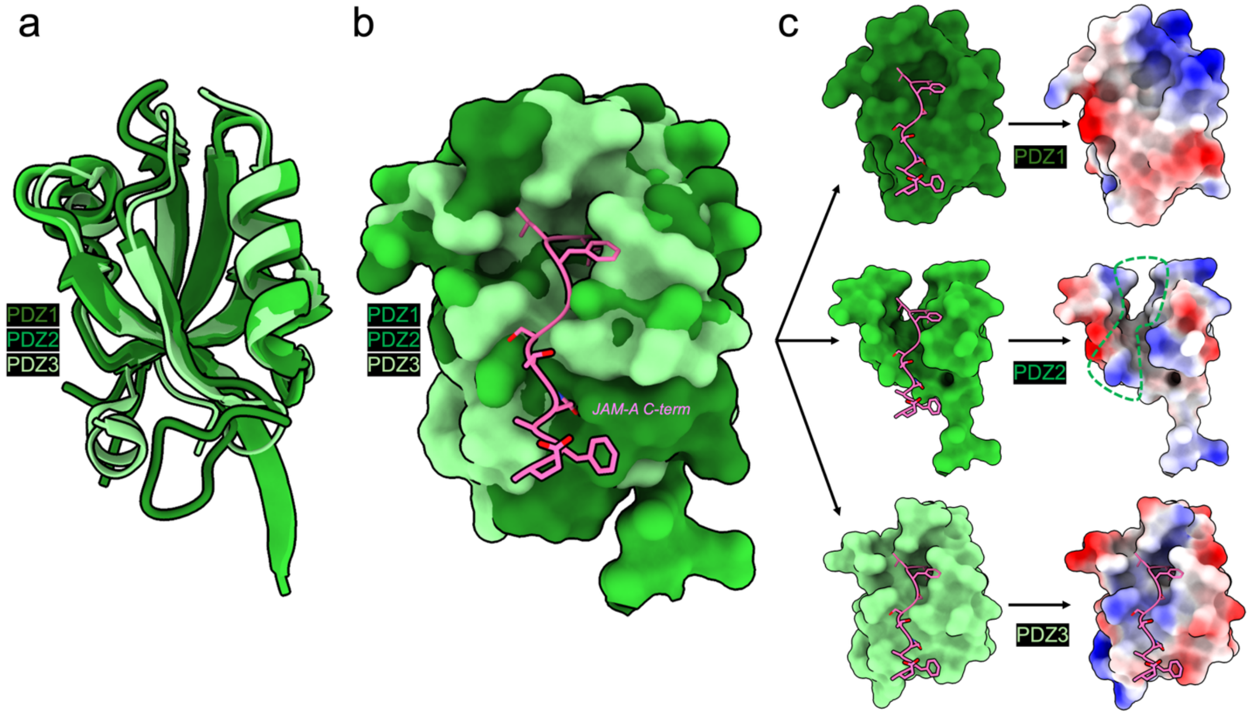


**Fig. S3. Structural homology of ZO-1 PDZ domains.** (a) Structural overlays of PDZ1 (dk green, PDB ID 2h3m), PDZ2 (green, PDB ID 2rcz), and PDZ3 (lt green, PDB ID 3shu) from experimentally determined crystal structures shown as cartoons. (b) Same overlays as in a but shown as surfaces. The C-terminal PDZ motif peptide from JAM-A (pink, cartoon) is shown in its experimentally determined pose when bound to the PDZ3 domain of PSG (PDB ID 3tsz) for reference. (c) Surface representations of all three PDZs bound to the JAM-A peptide from PDB ID 3tsz depicting the approximate binding pose for PDZ1 and PDZ2. Note the complex with PDZ3 is experimentally verified. Electrostatic surfaces are shown for each PDZ to highlight the differences in surface charge distribution despite structural conservation.

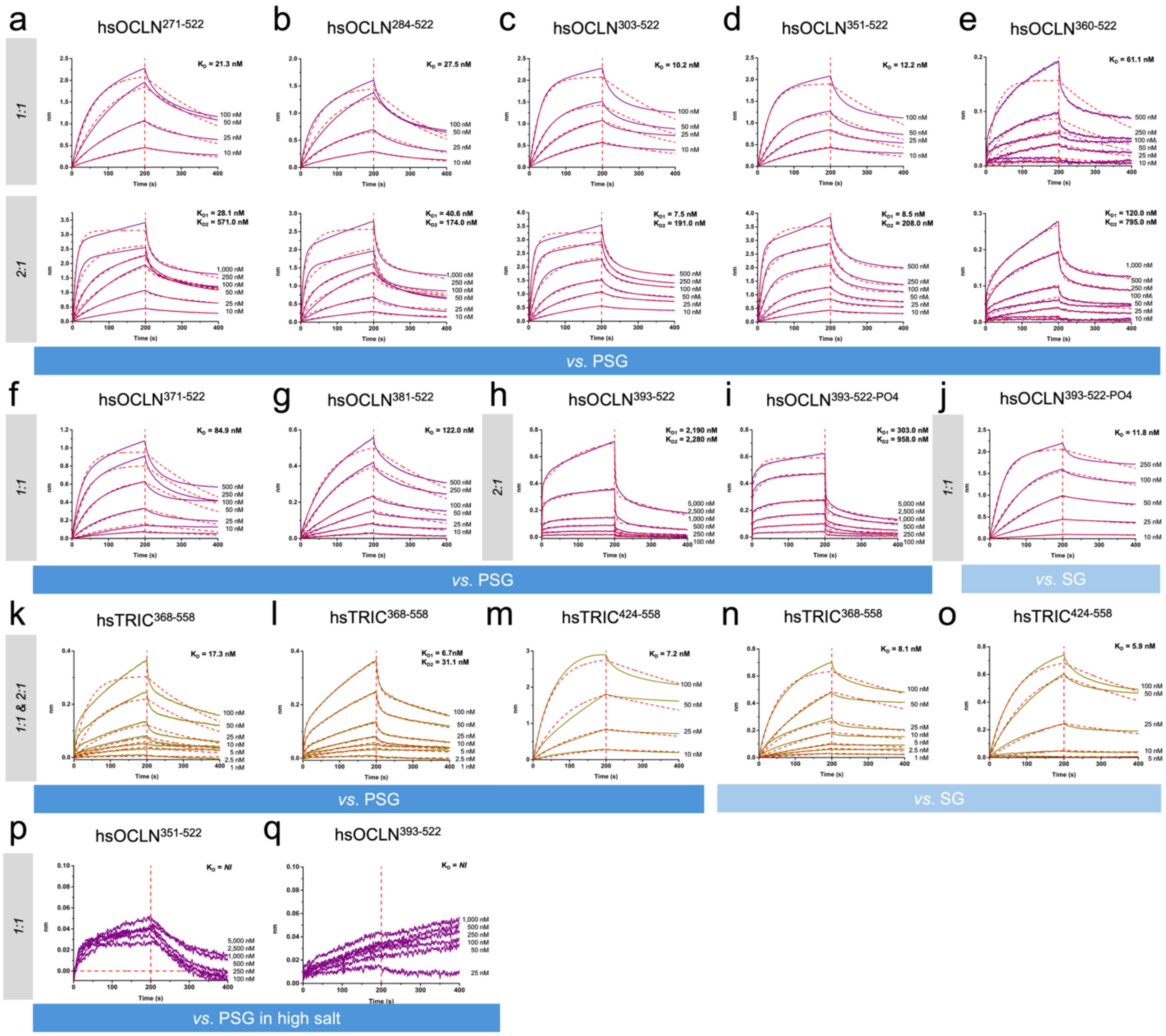


**Fig. S4. Biophysical interactions of PSG and SG *versus* TAMPs using BLI.** (a-j) BLI sensorgrams depicting binding interactions to occludin constructs. (k-o) BLI sensorgrams depicting binding interactions to tricellulin constructs. (p-q) BLI sensorgrams depicting binding interactions of PSG to occludin constructs in 500 mM NaCl. Experimental data is shown as thick line for occludin (purple) and tricellulin (green) with kinetic fits shown as dashed lines (red). Two $K_{D's}$ are reported where a heterogeneous 2:1 fit was used.

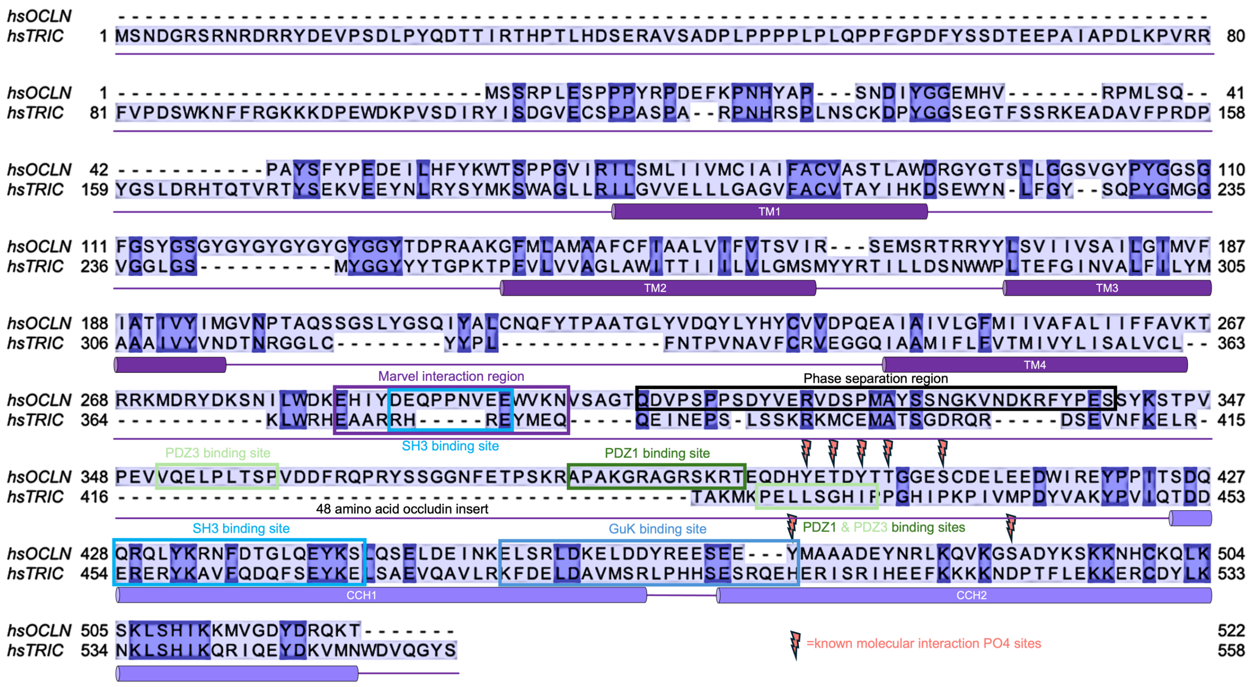


**Fig. S5. Annotated sequence alignment of human TAMPs.** Aligned sequences of human occludin (OCLN) and tricellulin (TRIC). Boxes represent approximate or experimentally determined binding sites of ZO-1 subdomain and other regions where molecular interactions of biological interest occur. Lightning bolts (red) represent phosphorylation sites determined from previous studies to alter ZO-1 binding and tight junction organization. Interestingly, ZO-1 domain binding sites overlap with phosphorylation sites and/or intra-molecular TAMP binding sites. This and all subsequent alignments were made in ClustalOmega[2] and visualized using Jalview.[3]

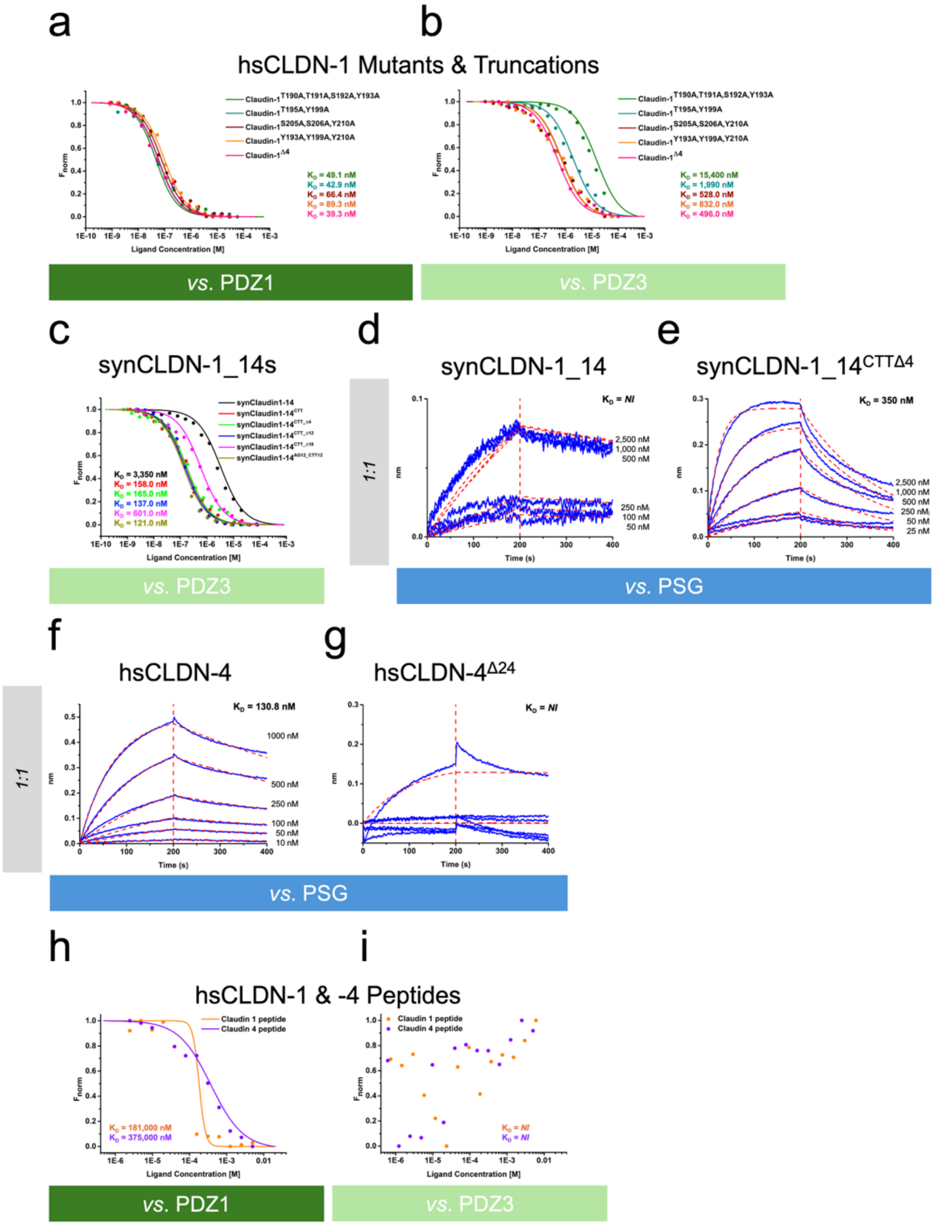


**Fig. S6. Biophysical interactions between ZO-1 subdomains and claudins.** (a-b) Binding of claudin-1 mutants or truncations against PDZ1 and PDZ3 using MST. (c-e) Binding of syntheticClaudin-1 (synCLDN-1) constructs against PDZ3 and PSG using MST and BLI. (f-g) Binding of claudin-4 and a truncation against PSG using BLI. (h-i) Binding of C-terminal peptides from claudin-1 and -4 against PDZ1 and PDZ3 using MST.

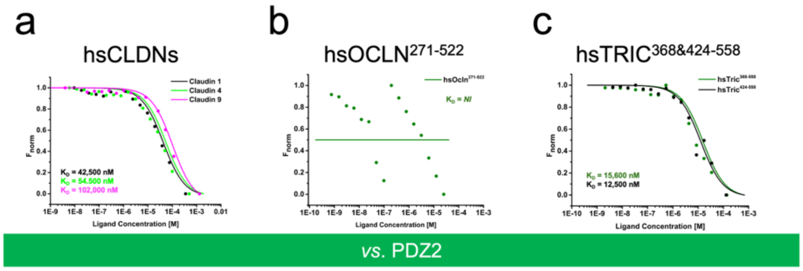


**Fig. S7. PDZ2 binding to TJMPs.** Binding of (a) human claudins -1, -4, and -9; (b) human occludin CTD; and (c) human tricellulin CTD against the PDZ2 domain of ZO-1.

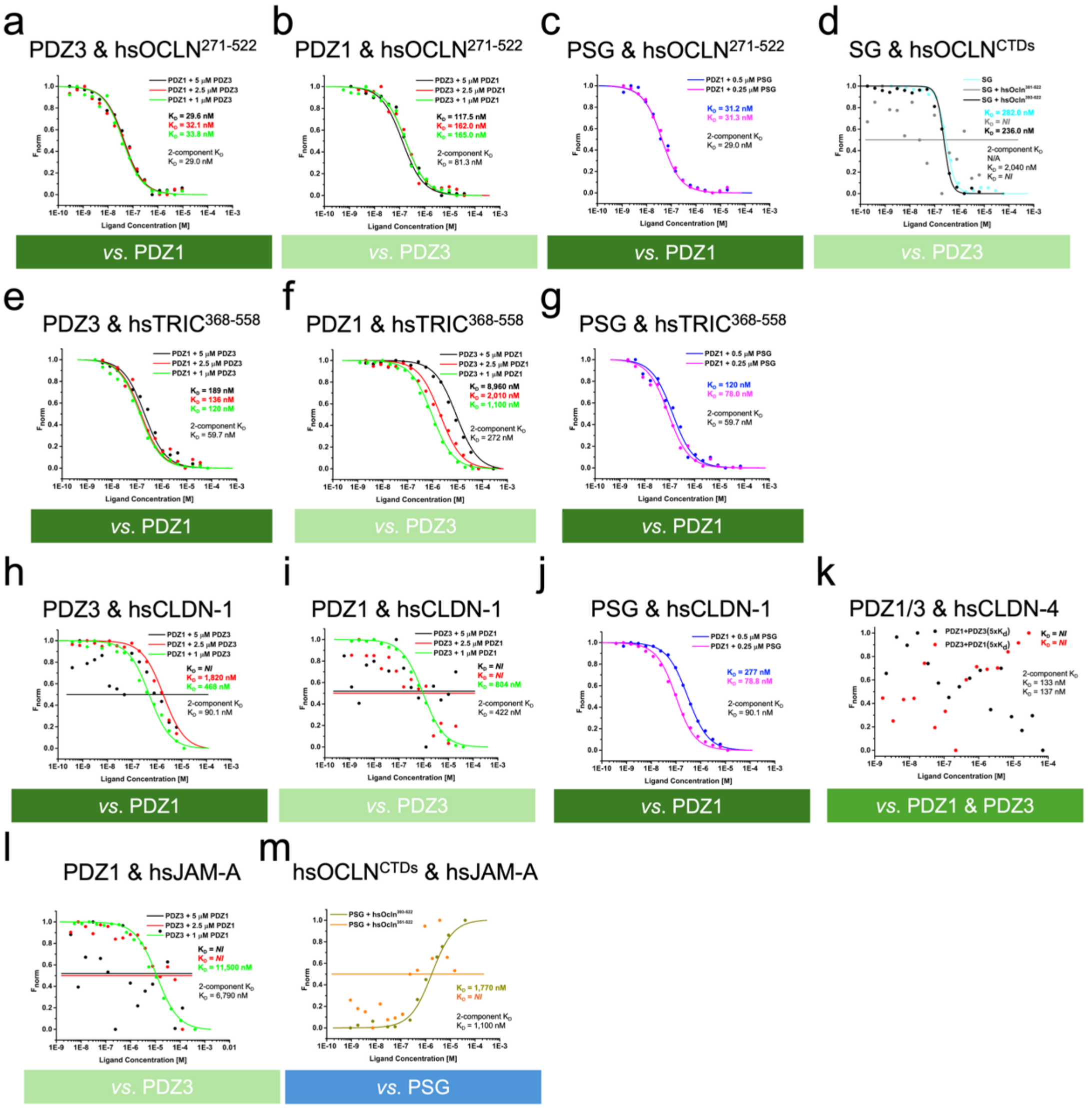

**Fig. S8. Three-component binding studies.** ZO-1 subdomains were pre-incubated with one analyte at a concentration above its $K_D$, then a second analyte was added to this mixture in various concentrations. Experiments were conducted with two ZO-1 subdomains against a second TJMP analyte that included (a-d) human occludin CTD; (e-g) human tricellulin CTD; (h-j) human claudin-1 and (k) claudin-4; and (l) human JAM-A. For (m), two TJMPs were used as analytes against PSG.

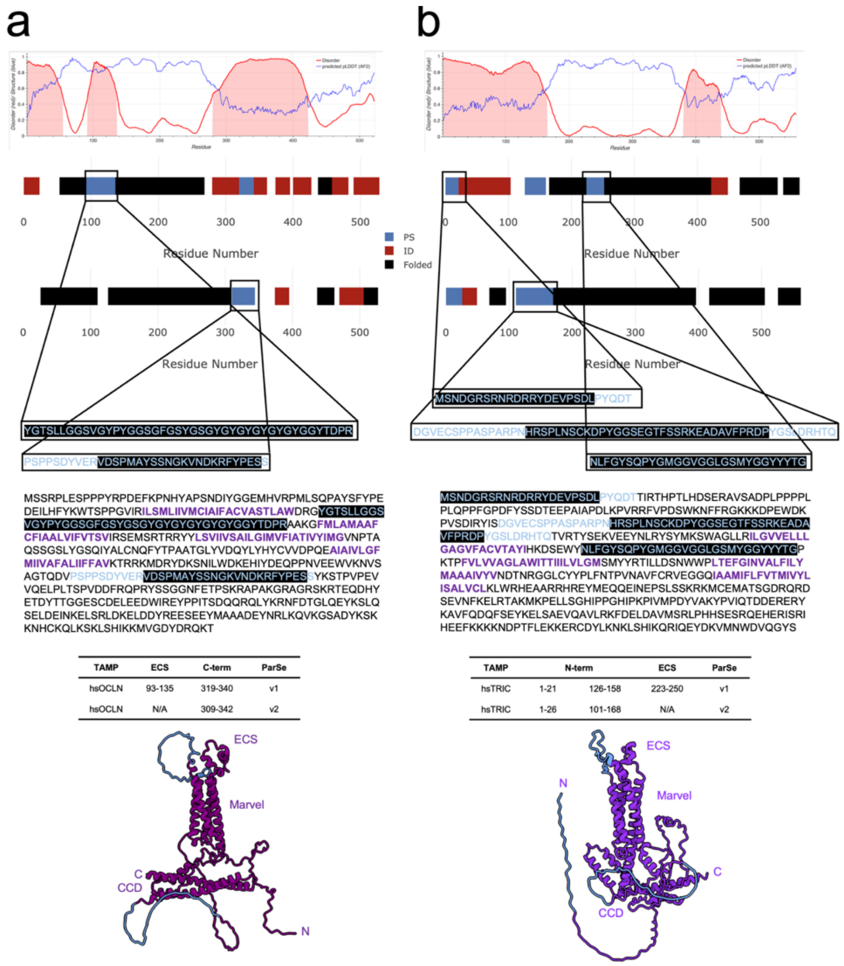


**Fig. S9. Predicted phase transition behavior of TAMPs using the ParSe servers.** The ParSe servers versions 1[4] and 2[5] were used to predict phase separation potential of human (a) occludin and (b) tricellulin. Disorder predictions (MetaPredict 3.0)[6] are plotted above ParSe results where phase separation (blue), intrinsically disordered regions (red) and folded domains (black) are represented by boxes. For phase separating regions, we have provided the associated sequences, which can then be compared to their positions within the full primary sequence. These regions are highlighted (black) with text (blue). Tables represent a summary of the results from each server. Predicted 3D structures from AlphaFold[7] are provided showing occludin (purple) and tricellulin (grape) with phase separating regions colored (lt blue).

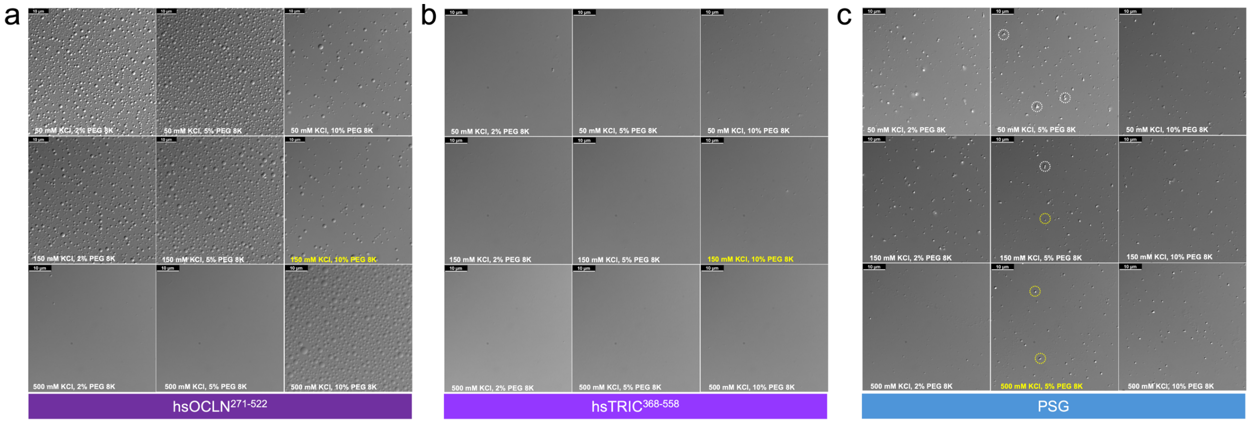


**Fig. S10. Buffer screening for conditions that yield phase separation of occludin, tricellulin, and PSG.** We screened constructs (a) hsOCLN$^{271-522}$, (b) hsTRIC$^{368-558}$, and (c) PSG at 10 μM in nine buffer conditions that included KCl, polyethylene glycol 8,000 (PEG 8K), and 20 mM HEPES pH 7.4, known from other systems to yield phase separation. Results show robust droplet formation for hsOCLN$^{271-522}$ but not hsTRIC$^{368-558}$. For PSG, low KCl concentrations yielded mixed species of aggregates (white circle) and droplets (yellow circle) whereas high KCl shifted this equilibrium toward droplet formation. The preferred buffer conditions for each protein are highlighted (orange).

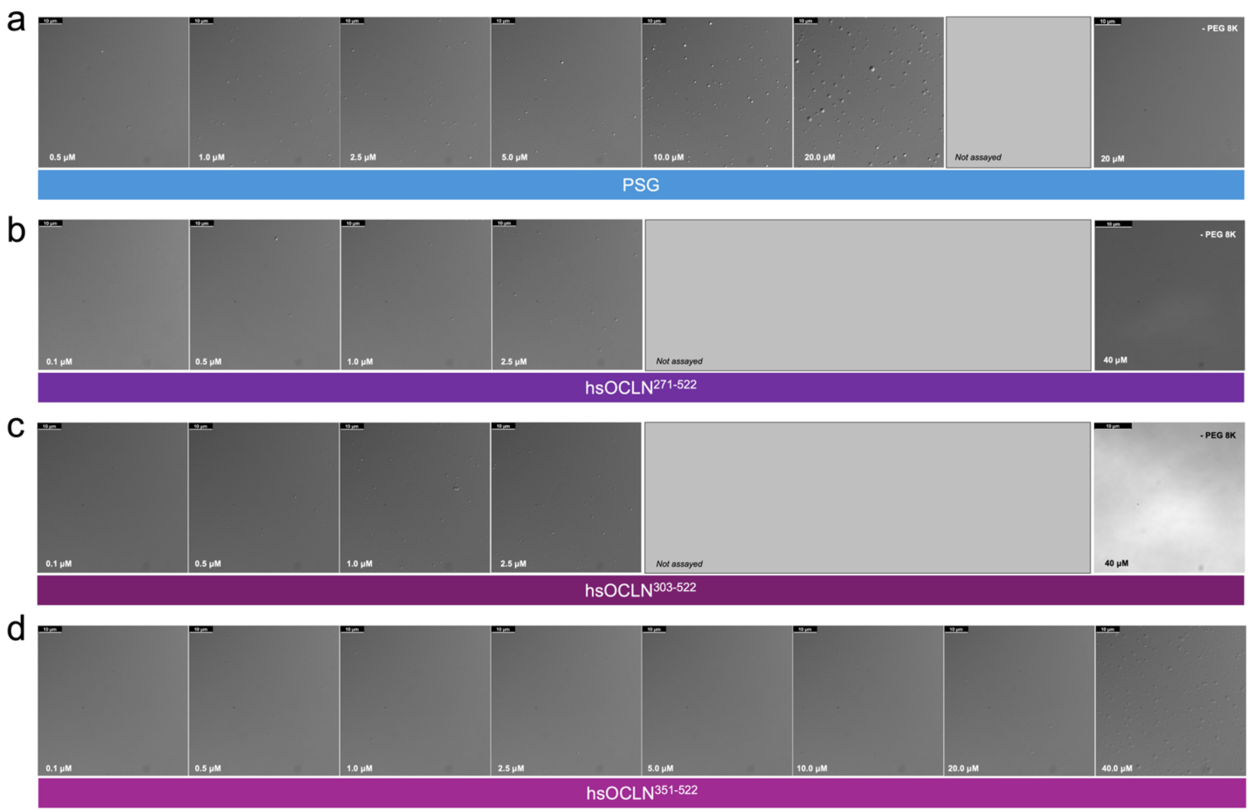


**Fig. S11. Phase separation of PSG and occludin at physiological concentrations.** We used buffers most amenable for phase separation to screen concentrations of (a) PSG (b) $hsOCLN^{271-522}$ (c) $hsOCLN^{303-522}$ (d) $hsOCLN^{351-522}$ that yielded phase separation. For PSG, 20 mM HEPES pH 7.4, 500 mM KCl, and 10% PEG 8K showed that it forms phase condensates at >1 μM. For $hsOCLN^{303-522}$ and $hsOCLN^{351-522}$, 20 mM HEPES pH 7.4, 500 mM KCl, and 10% PEG 8K showed that it forms phase condensates between 1-2.5 μM while for $hsOCLN^{351-522}$, it required >20 μM to form droplets. This verified that residues 271-350 were required for phase separation. As controls, in 20 mM HEPES pH 7.4, 150 mM NaCl without PEG 8K we found neither PSG nor occludin formed droplets.

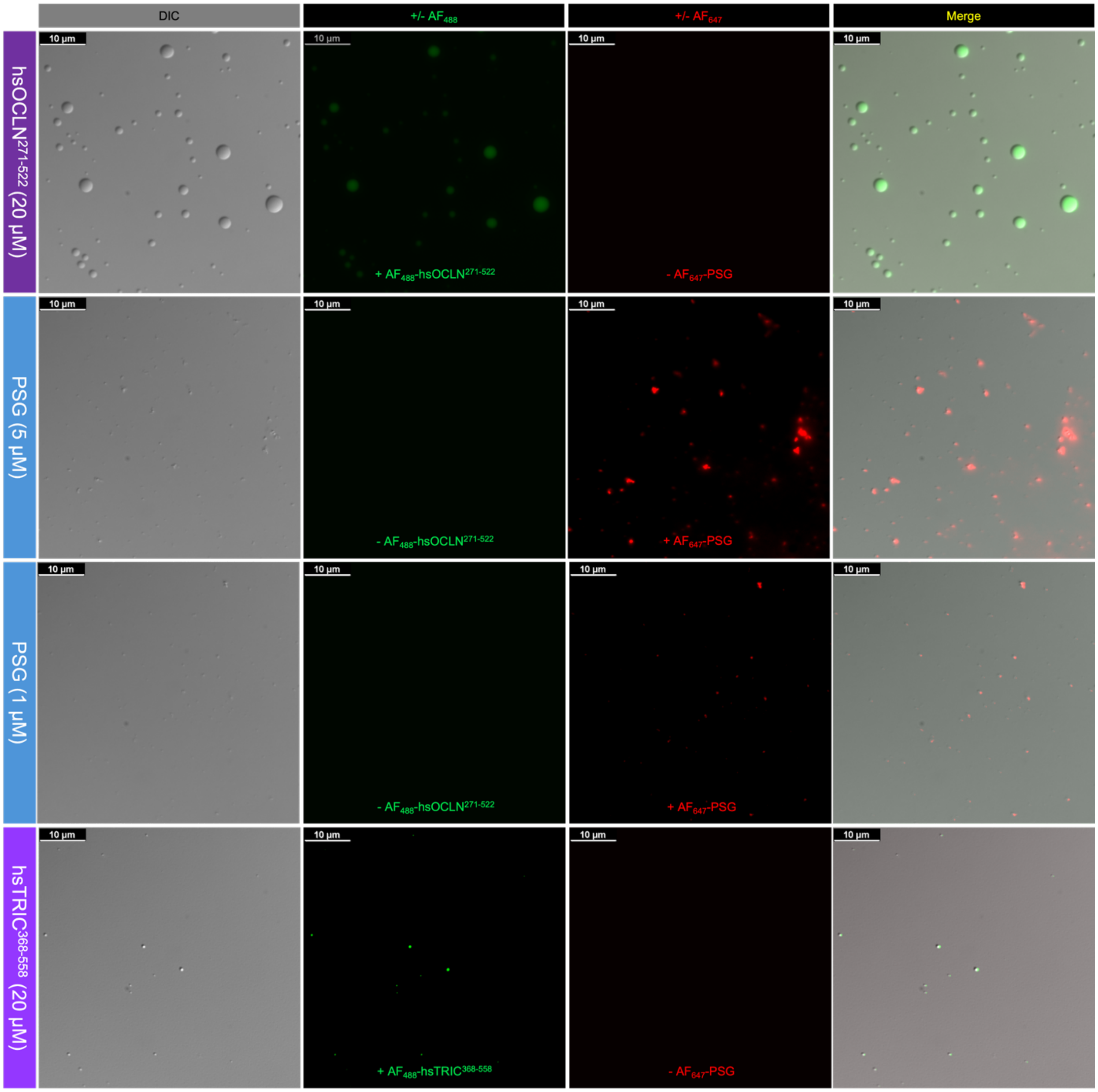


**Fig. S12. Controls for client partition assay in buffer 20 mM HEPES pH 7.4, 100 mM NaCl, 10% PEG 8K.** 20 µM hsOCLN$^{271-522}$ containing 1% Alexa Fluor 488 labeled hsOCLN$^{271-522}$ phase separated. 5 µM Alexa Fluor 647 labeled PSG showed aggregation. 1 µM Alexa Fluor 647 labeled PSG showed aggregation. 20 µM hsTRIC$^{368-558}$ containing 1% Alexa Fluor 488 labeled hsTRIC$^{368-558}$ showed very low phase separation.

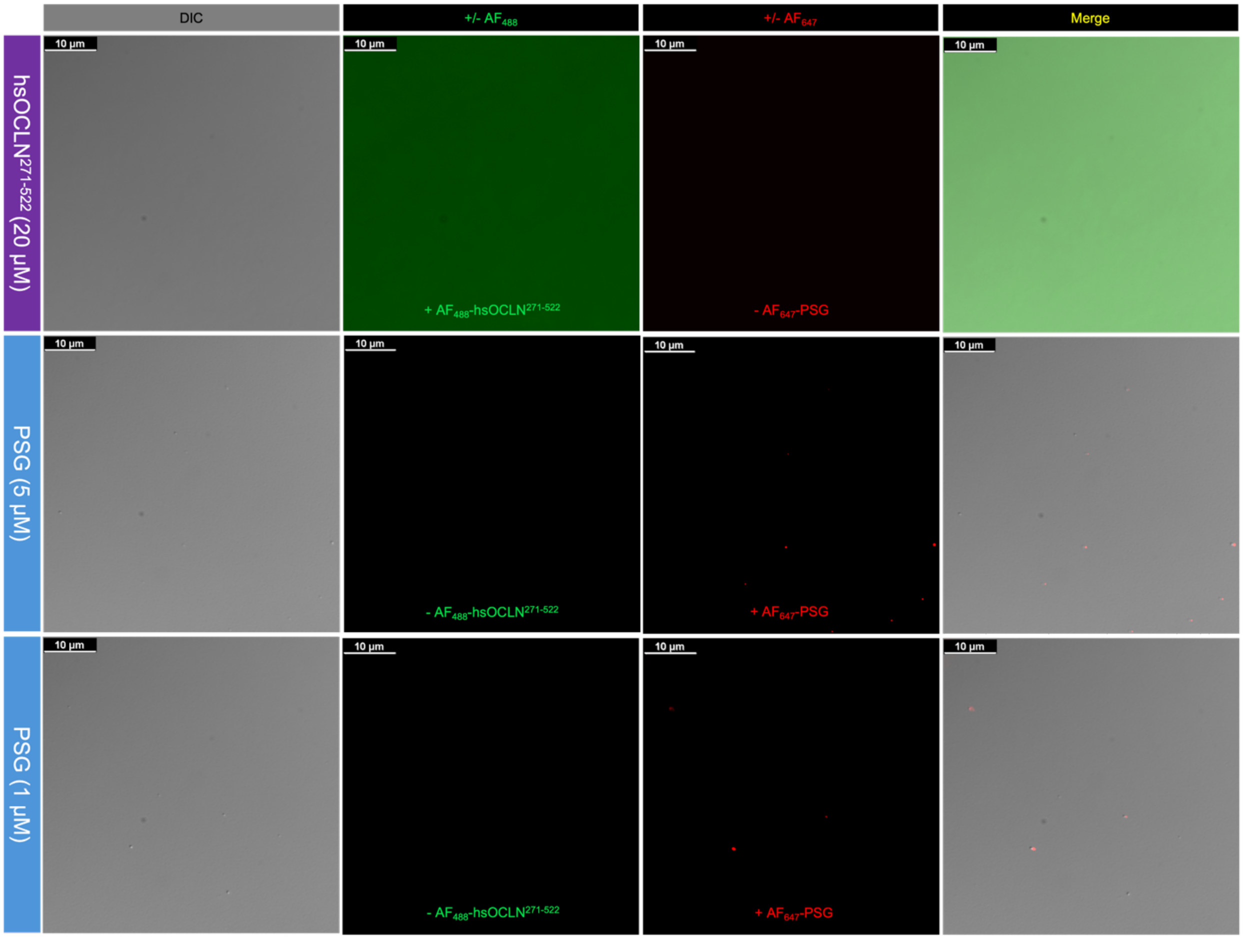


**Fig. S13. Controls for client partition assay in buffer 20 mM HEPES pH 7.4, 500 mM NaCl, 5% PEG 8K.** 20 µM hsOCLN$^{271\text{-}522}$ containing 1% Alexa Fluor 488 labeled hsOCLN$^{271\text{-}522}$ showed no phase separation. 5 µM Alexa Fluor 647 labeled PSG showed aggregation. 1 µM Alexa Fluor 647 labeled PSG showed phase separation.

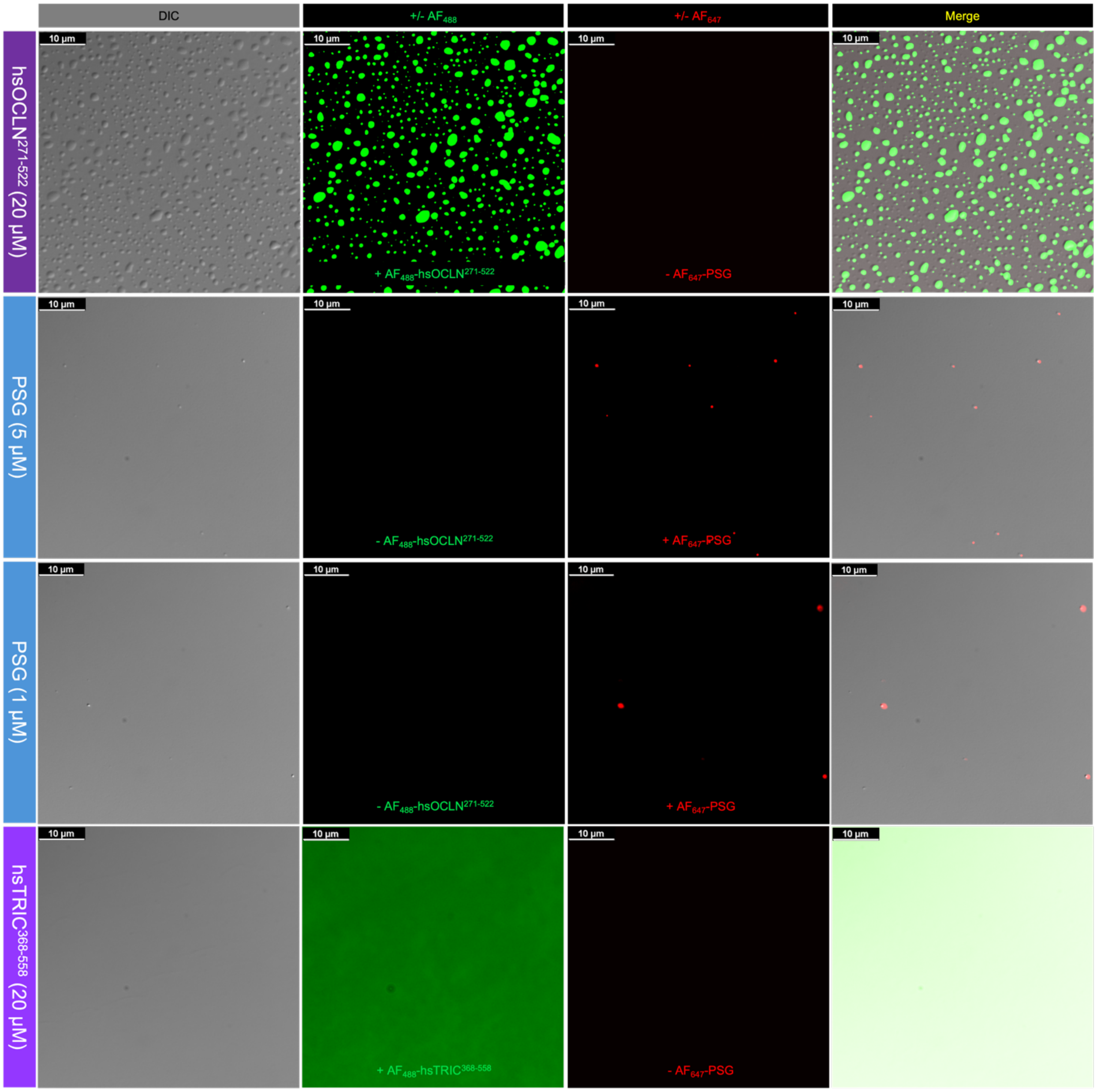


**Fig. S14. Controls for client partition assay in buffer 20 mM HEPES pH 7.4, 500 mM NaCl, 10% PEG 8K.** 20 µM hsOCLN$^{271\text{-}522}$ containing 1% Alexa Fluor 488 labeled hsOCLN$^{271\text{-}522}$ showed phase separation. 5 µM Alexa Fluor 647 labeled PSG showed phase separation. 1 µM Alexa Fluor 647 labeled PSG showed phase separation. 20 µM hsTRIC$^{368\text{-}558}$ containing 1% Alexa Fluor 488 labeled hsTRIC$^{368\text{-}558}$ showed no phase separation.

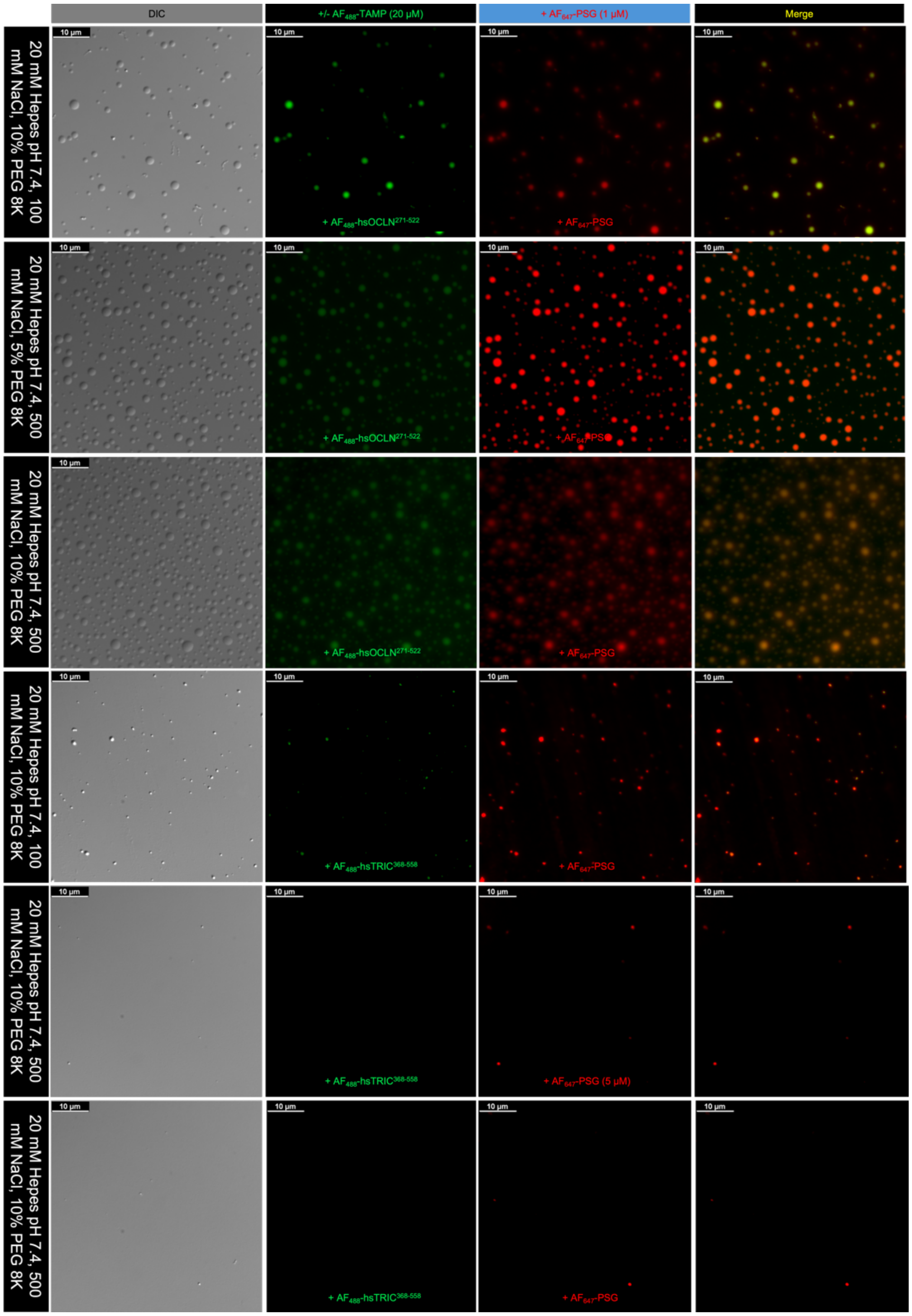


**Fig. S15. Client partition assay confirms that hsOCLN$^{271\text{-}522}$ and PSG cohabit the same droplets.** Phase condensates of 20 μM Alexa Fluor 488 labeled hsOCLN$^{271\text{-}522}$ or hsTRIC$^{368\text{-}588}$ in the presence of 1 μM Alexa Fluor 647 labeled PSG in various buffers. Note that in all buffers and salts tested that occludin and PSG cohabited the same drops and that only in 100 mM NaCl did tricellulin perform similarly. High salt abrogated tricellulin/PSG cohabitation.

**Table S1. Phase separation prediction of TJMPs using ParSe servers**

| Protein | PS | Region | Sequence | Domain |
|---|---|---|---|---|
| Claudin-1 | No | N/A | N/A | N/A |
| Claudin-2 | No | N/A | N/A | N/A |
| Claudin-3 | Yes | 202-220 | RSTGPGASLGTGYDRKDYV | IC, C-term |
| Claudin-4 | No | N/A | N/A | N/A |
| Claudin-5 | No | N/A | N/A | N/A |
| Claudin-6 | Yes | 177-207 | GGGLLCCTCPSGGSQGPSHYMARYSTSAPAISRGPSEYPTKNYV | IC, C-term |
| Claudin-7 | No | N/A | N/A | N/A |
| Claudin-8 | No | N/A | N/A | N/A |
| Claudin-9 | No | N/A | N/A | N/A |
| Claudin-10 | No | N/A | N/A | N/A |
| Claudin-11 | No | N/A | N/A | N/A |
| Claudin-12 | No | N/A | N/A | N/A |
| Claudin-14 | No | N/A | N/A | N/A |
| Claudin-15 | Yes | 206-228 | MPVATSDQEGDSSFGKYGRNAYV | IC, C-term |
| Claudin-16 | No | N/A | N/A | N/A |
| Claudin-17 | No | N/A | N/A | N/A |
| Claudin-18 | Yes | 231-261 | FGSNTKNKKIYDGGARTEDEVQSYPSKHDYV | IC, C-term |
| Claudin-19 | No | N/A | N/A | N/A |
| Claudin-20 | No | N/A | N/A | N/A |
| Claudin-23 | No | N/A | N/A | N/A |
| Claudin-25 | No | N/A | N/A | N/A |
| JAM-A | No | N/A | N/A | N/A |
| JAM-B | Yes | 166-191 | YTWFKDGIRLLENPRLGSQSTNSSYT | EC, N-term |
| JAM-C | Yes | 276-297 | KQDGESYKNPGKPDGVNYIRTD | IC, C-term |
| Occludin | Yes✓ | 93-135 | YGTSLLGGSVGYPYGGSGFGSYGSGYGYGYGYGYGYGGYTDPR | EC, ECS1 |
| | Yes✓ | 319-340 | VDSPMAYSSNGKVNDKRFYPE | IC, C-term |
| Tricellulin | Yes | 1-21 | MSNDGRSRNRDRRYDEVPSDL | IC, N-term |
| | Yes | 126-158 | HRSPLNSCKDPYGGSEGTFSSRKEADAVFPRDP | IC, N-term |
| | Yes | 223-250 | NLFGYSQPYGMGGVGGLGSMYGGYYYTG | EC, ECS1 |
| MarvelD3-1 | Yes | 50-82 | NRRRDGDRDPERDQERDGNRDRNRDRERERERER | IC, N-term |
| *@tri-cell* | Yes | 391-410 | SREQPGSYSDAPEYLWSGTL | IC, C-term |
| MarvelD3-2 | Yes | 50-82 | NRRRDGDRDPERDQERDGNRDRNRDRERERERER | IC, N-term |
| *@bi-cell* | Yes | 222-250 | STGGYTGITSLGGIYYYQFGGAYSGFDG | EC, ECS1 |
| ZO-1 | Yes✓ | 323-345 | SDHSRHSPQQPSNGSLRSRDEER | IC, U3 motif |

ParSe server versions 1[4] and 2[5] were used in analyses.
IC (intracellular); EC (extracellular)
✓ indicates phase separation has been validated experimentally.

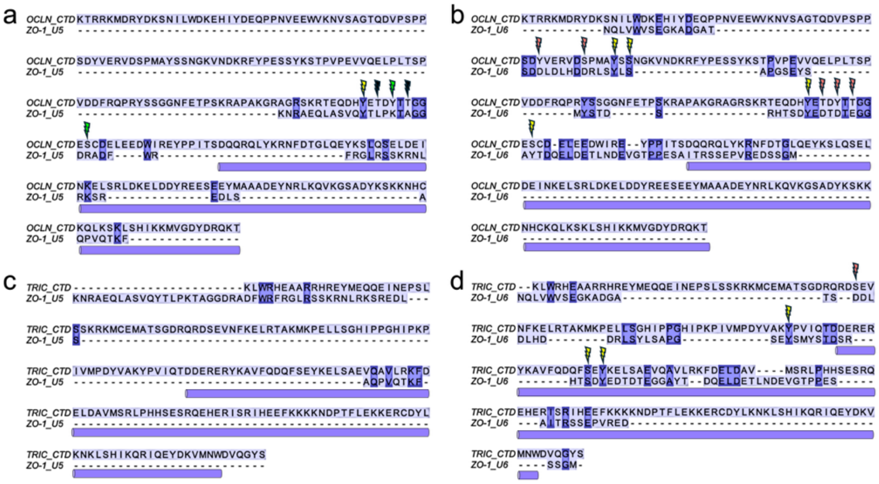

e

| Protein | pI |
|---|---|
| Occludin CTD | 5.5 |
| Occludin IDR | 5.0 |
| Occludin CCD | 7.7 |
| Tricellulin CTD | 8.7 |
| Tricellulin IDR | 9.4 |
| Tricellulin CCD | 6.5 |
| PDZ3 | 5.2 |
| SH3 | 8.2 |
| PS | 7.0 |
| U5 | 11.3 |
| SG | 9.8 |
| GuK | 9.5 |
| PSG | 9.6 |
| U6 | 3.8 |
| Occludin U5 homolog | 5.1 |
| Occludin U6 homolog | 4.1 |
| Tricellulin U5 homolog | 9.7 |
| Tricellulin U6 homolog | 7.2 |

**Fig. S16. Hidden sequence conservation between ZO-1 U motifs and the C-terminal domains of TAMPs.** Sequence alignments between ZO-1's U5 and U6 motifs against (a-b) occludin's CTD and (c-d) tricellulin's CTD. Lightning bolts represent residues on TAMPs known to alter ZO-1 binding or tight junction organization. Red bolts indicate residues where phosphorylation of the TAMP would yield the same charge as the U motif residue; Yellow bolts indicate residues that could be phosphorylated in either the TAMP or U motif to yield similar charges; Green bolts indicate residues where phosphorylation of the TAMP would yield the opposite charge as the U motif residue; and Black bolts indicate residues where phosphorylation of the TAMP would yield a charged residue while U motif would yield a hydrophobic one. (e) Sequences representing each protein or domain had their isoelectric points (pI) calculated and are reported here. Note how the IDR regions of TAMPs and U motifs of ZO-1 influence the pI of the larger domains or modules. Interestingly, the U5 homologous region of occludin and the U6 homologous region of tricellulin have very different pIs than their counterparts despite sequence homology. For instance, U5 is very basic at 11.3 while the U5 homology region of occludin is acidic at 5.1 This would yield antagonistic binding/unbinding events in favor of regulation.

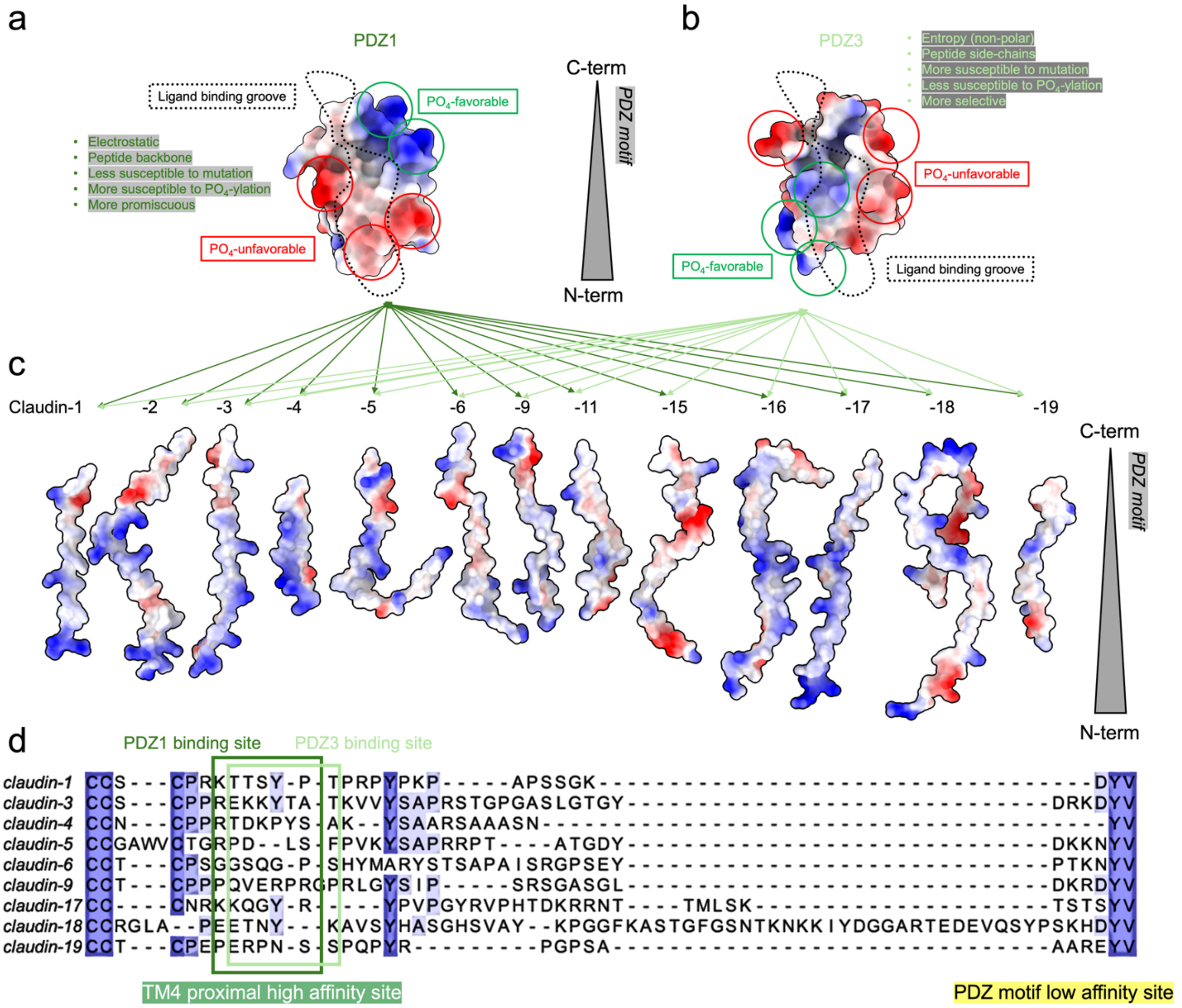


**Fig. S17. PDZ-specific mechanisms for binding claudins.** Electrostatic surfaces of the ligand binding grooves of (a) PDZ1 and (b) PDZ3, as shown in **Fig. S3**. Negative (red) and positive (blue) charges are colored accordingly. Based on crystal structures, ligands bind the grooves from C-terminus (top) to N-terminus (bottom). (c) Electrostatic surfaces of various claudin C-terminal domains are shown to highlight electrostatic similarities and differences. Very few regions show electrostatic homology apart from the N-terminus (TM4 proximal region) which in most claudins is positively charged followed by non-polar residues. This basic patch could bind the complimentary acidic patch at the base of the ligand binding groove of PDZ1 and position residues downstream of it toward the non-polar and basic regions of PDZ1. As electrostatics do not play a role in PDZ3 binding side chain shape complementarity likely plays a greater role and fit within the ligand binding groove of PDZ3 is the same orientation. (d) Based on the results of these analyses and those from binding studies, we propose the overlapping PDZ1 and PDZ3 binding sites on the intracellular domains of claudins.

### References


1. Li, Y., Fanning, A.S., Anderson, J.M., and Lavie, A. (2005). Structure of the conserved cytoplasmic C-terminal domain of occludin: identification of the ZO-1 binding surface. J Mol Biol *352*, 151-164. 10.1016/j.jmb.2005.07.017.
2. Madeira, F., Madhusoodanan, N., Lee, J., Eusebi, A., Niewielska, A., Tivey, A.R.N., Lopez, R., and Butcher, S. (2024). The EMBL-EBI Job Dispatcher sequence analysis tools framework in 2024. Nucleic Acids Res *52*, W521-W525. 10.1093/nar/gkae241.
3. Waterhouse, A.M., Procter, J.B., Martin, D.M., Clamp, M., and Barton, G.J. (2009). Jalview Version 2--a multiple sequence alignment editor and analysis workbench. Bioinformatics *25*, 1189-1191. 10.1093/bioinformatics/btp033.
4. Paiz, E.A., Allen, J.H., Correia, J.J., Fitzkee, N.C., Hough, L.E., and Whitten, S.T. (2021). Beta turn propensity and a model polymer scaling exponent identify intrinsically disordered phase-separating proteins. J Biol Chem *297*, 101343. 10.1016/j.jbc.2021.101343.
5. Wilson, C., Lewis, K.A., Fitzkee, N.C., Hough, L.E., and Whitten, S.T. (2023). ParSe 2.0: A web tool to identify drivers of protein phase separation at the proteome level. Protein Sci *32*, e4756. 10.1002/pro.4756.
6. Emenecker, R.J., Griffith, D., and Holehouse, A.S. (2021). Metapredict: a fast, accurate, and easy-to-use predictor of consensus disorder and structure. Biophys J *120*, 4312-4319. 10.1016/j.bpj.2021.08.039.
7. Varadi, M., Anyango, S., Deshpande, M., Nair, S., Natassia, C., Yordanova, G., Yuan, D., Stroe, O., Wood, G., Laydon, A., et al. (2022). AlphaFold Protein Structure Database: massively expanding the structural coverage of protein-sequence space with high-accuracy models. Nucleic Acids Res *50*, D439-D444. 10.1093/nar/gkab1061.
